# Omics-scale capture of electrophilic metabolites

**DOI:** 10.64898/2026.08.07.743530

**Authors:** Allen F. Schroeder, Yan Yu, Ali Turk, Henry H. Le, Matthew LeClair, Marissa A. Fontaine, Nicholas W. Cheng, Saeideh Azad, Christopher N. Parkhurst, Jason Pan, David Artis, Meng Wang, Frank C. Schroeder

## Abstract

The majority of metabolic pathways rely on the production of activated electrophilic intermediates, e.g., coenzyme-A esters, and their chemical structures and abundances are central to understanding enzyme function and biochemical mechanisms. However, most electrophilic metabolites are lost in traditional metabolomic analysis and thus remain poorly characterized. Here we introduce a biochemical probe, *O*-(trimethylammoniobutyl)-hydroxylamine (TAMOHA), that enables comprehensive profiling of electrophilic species such as coenzyme-A esters, ketones, and aldehydes. TAMOHA incorporates a highly nucleophilic hydroxyl amine that reacts quickly with electrophilic species upon tissue lysis, trapping them as stable derivatives that feature a tetraalkylammonium moiety whose constitutive charge and characteristic MS^2^ fragmentation fingerprint enable their highly sensitive detection. Using TAMOHA to survey the electrophilomes of *E. coli, C. elegans*, and mouse revealed several thousand electrophilic metabolites, most of which have not been characterized. We then demonstrate trapping of electrophilic metabolites with TAMOHA in the context of specific biochemical pathways, confirming previously proposed functions of two fatty acid metabolism enzymes, detecting formaldehyde production in mice, and providing new insights into the biosynthesis of ascaroside pheromones in *C. elegans*. We anticipate that use of TAMOHA for the profiling of electrophilic species will help clarify enzyme function and uncover previously elusive biochemical mechanisms in a wide range of biological systems.

**Table of Contents artwork:** 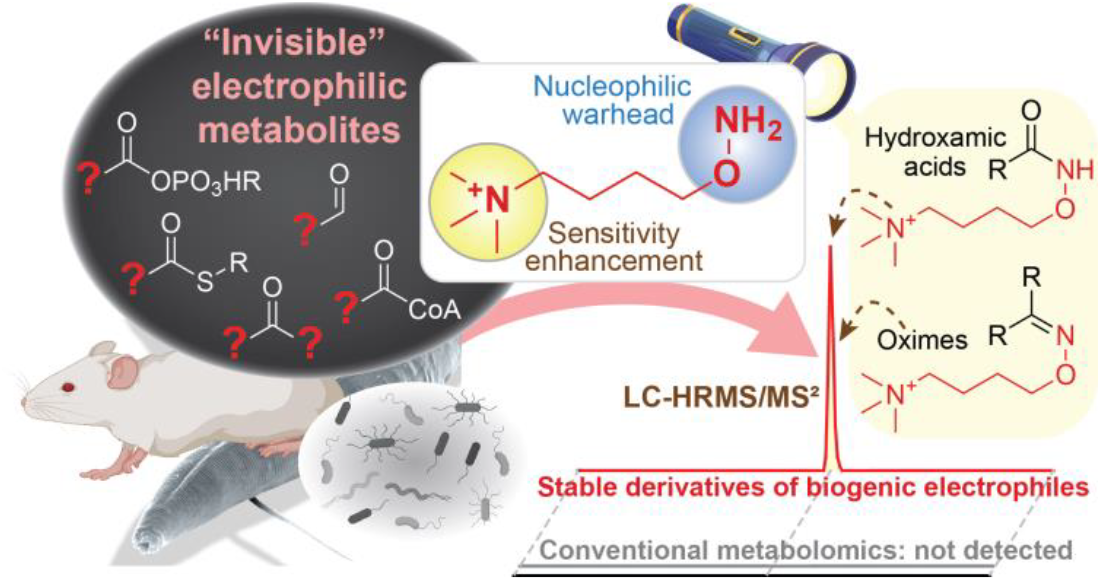

## INTRODUCTION

Recent advances in mass spectrometry have demonstrated that the metabolomes of even the best characterized model systems remain incompletely characterized^1^. This vast and chemically diverse space of unknown metabolites hints at the existence of complex networks of yet incompletely characterized biochemical pathways, many of which may rely on the generation of reactive electrophilic intermediates^2^. Such reactive metabolites include coenzyme-A (CoA) thioesters and acyl-AMP conjugates, which play central roles in metabolism and serve as building blocks for the biosynthesis of larger, more complex metabolites, but also have been shown to serve as cellular signals^2, 3^. Other electrophilic metabolites, for example aldehydes and ketones, are of interest because they are markers of oxidative damage^4^, contribute to protein and DNA damage^5^, and serve as important markers of disease^6^.

Identifying the structures and monitoring the abundances of reactive, electrophilic metabolites therefore seems essential for a comprehensive understanding of biosynthetic networks. However, the vast majority of electrophilic metabolites in living systems – the electrophilome^7^ – remain poorly characterized *in vivo*, because they undergo hydrolysis or react with biogenic nucleophiles at time scales that are incompatible with their characterization using conventional metabolomic analyses^2, 8^. Moreover, many electrophilic species exist only at very low abundances in biological systems^8^. Therefore, the characterization of electrophilic metabolites, though centrally important, remains as perhaps the most significant challenge in metabolomics.

Previous efforts to trap electrophilic metabolites as stable derivatives suitable for metabolomic analysis have employed hydroxylamine (HA)^9, 10, 11^. HA is uniquely well suited for trapping reactive thioesters, aldehydes, ketones and other electrophilic species as it is highly nucleophilic but not a strong base, and because the resulting hydroxamic acids and oximes are sufficiently stable for subsequent metabolomic analysis. However, the scope of HA-based metabolomic profiling is limited, given the low abundance of most electrophilic intermediates and because the resulting oximes and hydroxamic acids lack specific mass spectrometric features that would facilitate their detection or recognition in complex metabolomes, which are dominated by stable metabolic end products.

Aiming to develop a probe that would enable comprehensive profiling of activated metabolites, we were inspired by previous studies that take advantage of constitutively charged quaternary ammonium ions as ionization tags^12^. Here we describe O-(trimethylammoniobutyl)-hydroxylamine (TAMOHA, **1**), which traps reactive electrophilic metabolites by converting them into stable derivatives, dramatically enhances LC-MS sensitivity by including an ionization tag, and confers an easily recognizable MS^2^ fragmentation signature that distinguishes TAMOHA derivatives from unmodified metabolites (Fig. 1a). We demonstrate that using TAMOHA in combination with comparative mass spectrometric analysis^13, 14^ can reveal a vast space of electrophilic metabolites in diverse biological systems (Fig. 1b, c), most of which would not be detected in conventional metabolomic analyses. We then interrogate the ability of TAMOHA to trap two major classes of electrophilic metabolites in biological systems, CoA thioesters as well as aldehydes and ketones.

**Figure 1.**
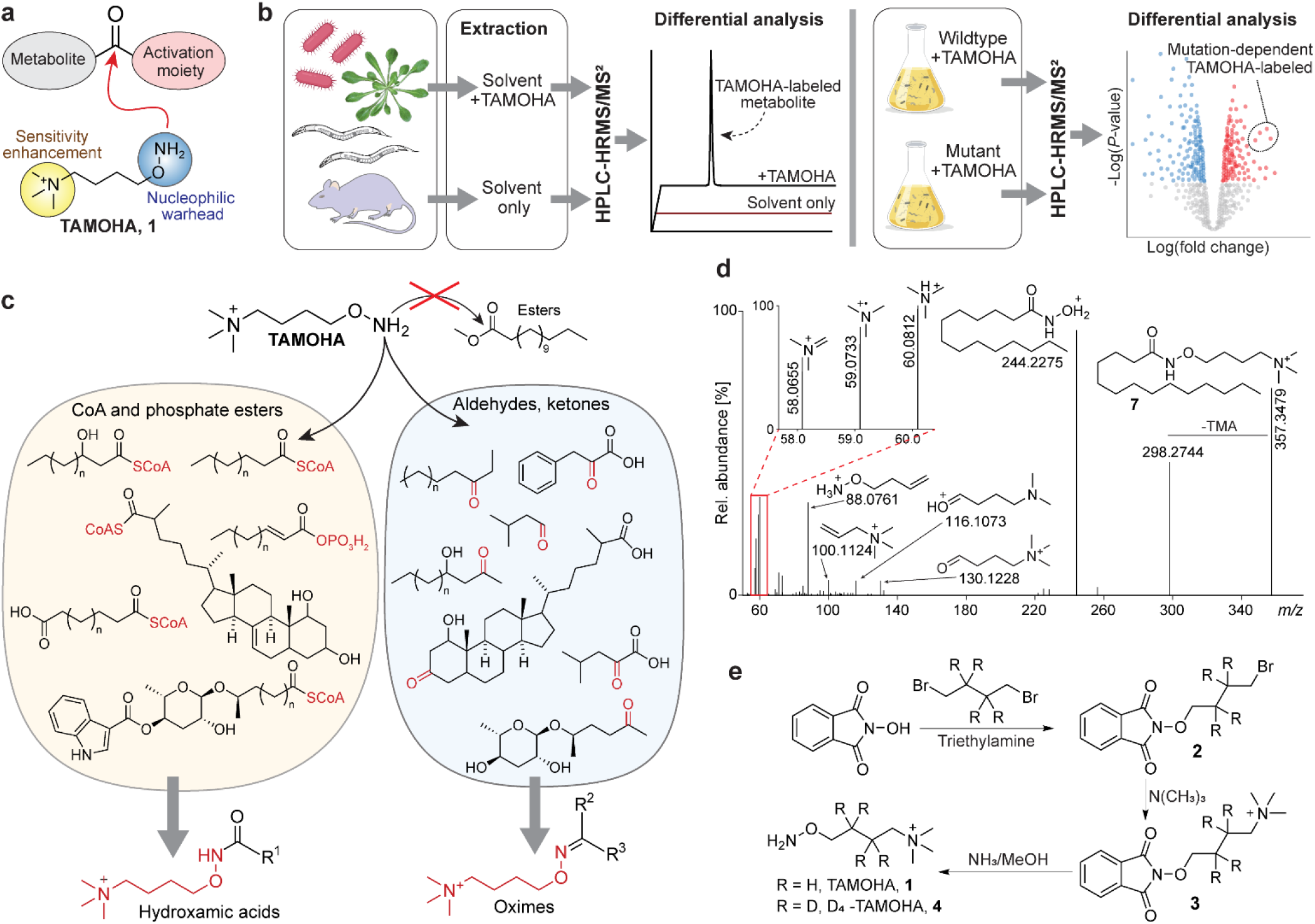
TAMOHA Synthesis and model studies. (a) Schematic of TAMOHA labeling strategy. (b) Scheme for the use of TAMOHA for global profiling of activated metabolites (left) or comparative metabolomic analyses of wildtype and mutant organisms to investigate enzyme function (right). (c) Overview of the scope of TAMOHA reactivity. (d) Annotated MS^2^ spectrum of the TAMOHA conjugate of tetradecanoic acid (**7**). (e) Synthesis of TAMOHA (**1**) and TAMOHA D_4_. (**4**).

## RESULTS

### Development of the TAMOHA probe

We set out to develop an HA-based probe that would enable (i) rapid trapping of electrophilic species upon cell lysis in the form of stable derivatives, (ii) improved sensitivity via enrichment or by enhancing the ionization efficiency of the trapped metabolites, and (iii) recognition of trapped metabolites by providing a unique fragmentation pattern in MS^2^ analysis. Toward this goal, we initially explored the utility of (azidoalkyl)-hydroxylamine derivatives (Fig. S1) for copper-catalyzed azide-alkyne cycloaddition-(CuAAC-) based enrichment^15^. CuAAC has been employed extensively to enrich proteins and peptides^10, 16^; and more recently to interrogate metabolism^17^. However, we found that the use of copper-based click-chemistry cocktails resulted in partial reduction of the N-O bond in the hydroxylamine moiety, limiting enrichment efficiency (Fig. S1a, b).

As an alternative strategy we explored ionization enhancement, linking HA with a constitutively charged tetraalkylammonium tag (Fig 1a). Probe design further aimed to ensure that conjugates resulting from trapping common electrophilic species such as thioesters, aldehydes and ketones would be sufficiently stable for subsequent indepth metabolomic analysis. This was a concern for *N-* alkyl-linked HA derivatives, which would likely offer superior nucleophilicity^18^; however, their reaction with aldehydes and ketones would result in formation of potentially reactive nitrones^19^ instead of the more stable oximes that reaction with *O*-alkyl HA derivatives would generate (Fig. S1c). Ultimately, we settled on the *O*-alkyl HA derivative TAMOHA (**1**), featuring an n-butyl chain linking HA and the trimethylammonium tag, which also allowed for facile integration of a stable-isotope label, as in D_4_-TAMOHA (**4**, Fig. 1e). Next, we conducted a series of model reactions to demonstrate that TAMOHA can effectively trap reactive electrophiles, such as CoA-thioesters and ketones, whereas less reactive electrophilic species, e.g., esters, are inert. As intended, TAMOHA treatment converted fatty acyl CoA-thioesters instantly into the corresponding hydroxamic acids and ketones into the corresponding oximes, whereas a fatty acyl methyl ester did not react (Fig. 1c and S2).

Inspection of the MS^2^ spectra of products from TAMOHA model reactions indicated that TAMOHA-labeled entities fragment in a highly characteristic manner. TAMOHA derivatives lose trimethylammonium in both homo- and heterolytic fashion to produce reporter ions at *m/z* 59.0733 and 60.0812. In addition, TAMOHA conjugates exhibit characteristic fragmentation about the N-O bond to produce several larger diagnostic fragments (Figs. 1d S3). Combined detection of this set of fragments enables highly confident and sensitive identification of TAMOHA conjugates. Detection of TAMOHA-conjugates can be further facilitated using D_4_-TAMOHA (**4**) in a SILAC-like workflow where samples are split three ways, treated with TAMOHA, D_4_-TAMOHA, or solvent only, and analyzed by HPLC-HRMS^20^. TAMOHA labeled features are then identified by their absence in solvent-treated controls, presence in TAMOHA treated samples, and presence in D_4_-TAMOHA treated samples with a mass shift of 4.0251 amu (Fig. S4), as demonstrated with specific examples below.

### TAMOHA labeling increases sensitivity

To test to what degree TAMOHA improves sensitivity for the detection of trapped metabolites relative to unmodified HA^10, 11^, we compared MS peak intensities of synthetic samples of corresponding HA and TAMOHA derivatives. We found that inclusion of the ionization tag in TAMOHA increased the MS response dramatically, for example, roughly 1000-fold comparing the oleic acid TAMOHA and HA derivatives (Fig. 2a). Ionization enhancement conferred by TAMOHA varied with compound structure and was somewhat lower for metabolites that, in their unmodified form, ionize better than oleic acid (Fig. S5). Importantly, TAMOHA conjugates generally produced much sharper elution profiles on conventional C18 reverse-phase columns compared to the corresponding HA derivatives (see e.g., Fig. 2a), further aiding in their detection.

**Figure 2.**
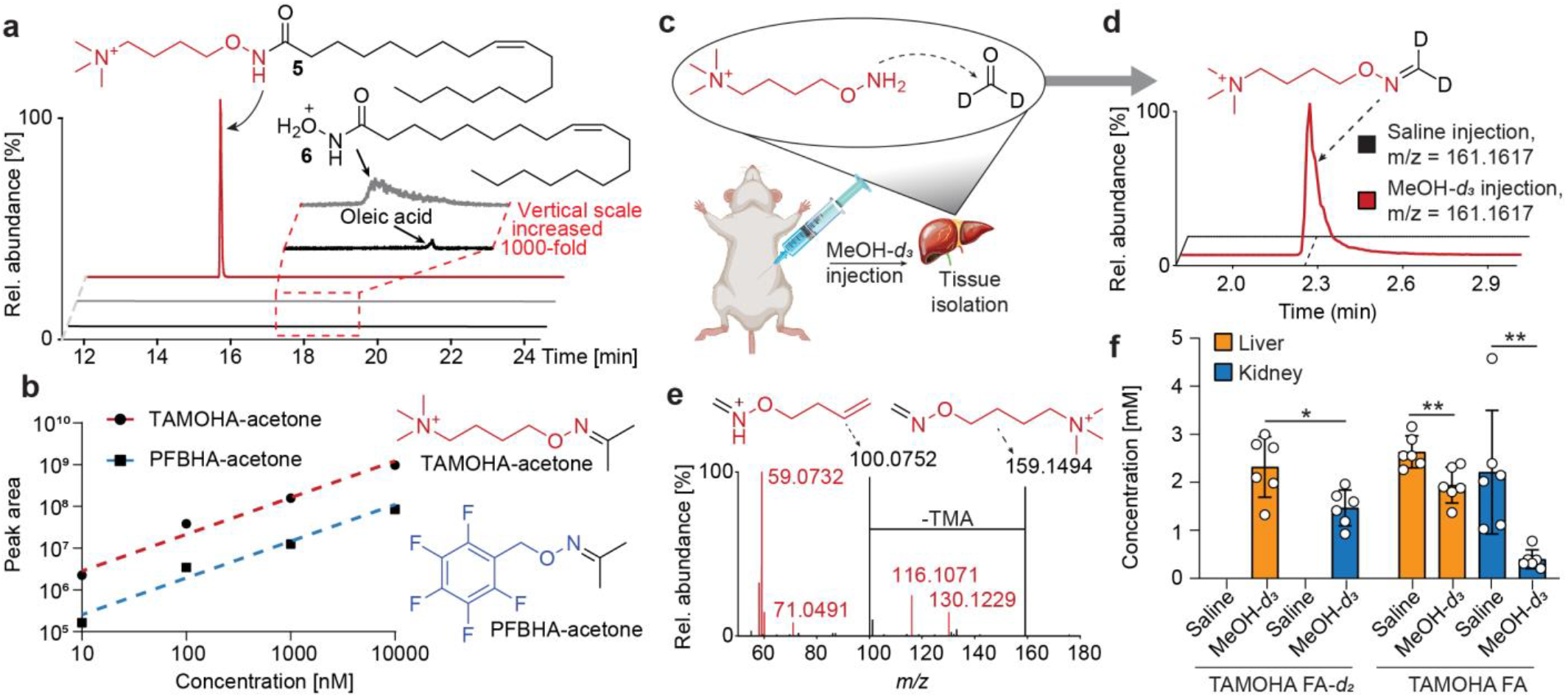
TAMOHA greatly increases sensitivity. (a) Ion chromatograms of synthetic standards of the oleic acid TAMOHA conjugate (**5**) (top, positive electrospray ionization mode (ESI+)), *N*-hydroxyoleamide (**6**) (middle, ESI+), and oleic acid (bottom, ESI-), each at 10 µM. (b) HPLC-HRMS-detected peak area for the TAMOHA-(red) or PFBHA-(blue) acetone conjugate at different concentrations. (c) Scheme for the detection of methanol-*d*_*3*_-derived formaldehyde in mouse TAMOHA formaldehyde labeling. (d) Ion chromatograms representing TAMOHA-labeled formaldehyde-*d*_*2*_ obtained from mice injected intraperitoneally with vehicle (top) or MeOH-*d*_*3*_ (bottom). (e) MS^2^ of TAMOHA-labeled formaldehyde. (f) Quantification of TAMOHA-labeled formaldehyde-*d*_*2*_ (TAMOHA-FA-*d*_*2*_, left) and TAMOHA-labeled endogenous (non-deuterated) formaldehyde (TAMOHA-FA) right) in liver or kidney from animals injected intraperitoneally with vehicle or MeOH-*d*_*3*_. Note that levels of endogenous, unlabeled formaldehyde are reduced in MeOH-*d*_*3*_-injected animals, indicating that formaldehyde production or detoxification are perturbed as a result of MeOH-*d*_*3*_-injection.

To assess to what extent TAMOHA facilities detection of aldehydes and ketones, we compared the utility of TAMOHA and *O*-(2,3,4,5,6-pentafluorobenzyl)hydroxylamine (PFBHA), a derivatization reagent previously used to trap formaldehyde and other carbonyl species via GC-MS^21^. For this comparison we selected formaldehyde and acetone, which both play important roles in biological systems but are generally difficult to detect, due to their volatility and tendency to react with diverse nucleophiles. Formaldehyde is generated in many biological contexts, but highly toxic, causing DNA lesions and protein cross-linking^22^, and the resulting cellular damage is suspected to play a role in the etiology of a wide range of human diseases^23^. Measurement of acetone levels is relevant, e.g., in the context of diabetic ketoacidosis and other metabolic diseases^24^.

In the case of acetone, both TAMOHA and PFBHA yielded the expected oximes, with the TAMOHA oxime producing more than an order of magnitude higher signal than the PFBHA oxime across the tested concentration range (Fig. 2b). Previous efforts using PFBHA and GC-MS enabled detection of formaldehyde down to low micromolar levels, e.g., in mouse serum^25, 26^. Using TAMOHA and HPLC-HRMS, formaldehyde could be detected down to ~5 nM levels, whereas only trace amounts of the formaldehyde PFBHA derivative were detected using HPLC-HRMS, even at the highest tested concentrations (Fig. S6a)

Lastly, we developed a labeling strategy in which fresh cell or tissue samples are lysed in the presence of TAMOHA to enable immediate trapping of activated metabolites at the time of harvest. We then tested the utility of TAMOHA for the detection of formaldehyde *in vivo*, using mice with a loss-of-function mutation in the alcohol dehydrogenase *adh*5, which plays a central role in formaldehyde detoxification^25^. *adh5* mutant mice were injected intraperitoneally with either methanol-*d*_*3*_ or saline control, based on prior work that showed that intraperitoneal methanol is metabolized into formaldehyde^27^ (Fig. 2c). HPLC-HRMS analysis of liver and kidney extracts harvested 24 h later demonstrated sensitive detection of TAMOHA-formaldehyde-*d*_*2*_ in samples obtained from methanol-*d*_*3*_-injected animals, in addition to the TAMOHA-derivatives of unlabeled, endogenous formaldehyde and other short-chain aldehydes (Fig. 2d, e, Fig. S6b). Taken together, these results show that TAMOHA labeling confers substantial sensitivity gains for the detection of thioesters and other reactive carbonyl species^25^.

### The scope of TAMOHA labeling

To get a rough estimate for the number of electrophilic species that TAMOHA labeling could reveal in living systems, we selected cell and tissue samples from three model systems, *E. coli* (cell pellet) *C. elegans* (whole body) and mouse (intestine). Following extraction and HPLC-HRMS analysis, TAMOHA-labeled features were identified by comparing the TAMOHA-treated and untreated control samples using the xcms and Metaboseek platforms^13^ and then validated based on the diagnostic MS^2^ fragmentation pattern of the TAMOHA moiety (see Methods). TAMOHA treatment uncovered a vast space of electrophilic species in these organisms. After removal of artifacts and obvious fragments from the MS data, we detected 3644, 9109, and 9801 TAMOHA labeled mass features in the *E. coli, C. elegans*, and mouse samples, respectively. This corresponds to more than a third of the number of non-TAMOHA-labeled mass features detected in the untreated control samples (Fig. 3a), consistent with the idea that active metabolism involves large numbers of electrophilic intermediates from diverse biochemical pathways.

**Figure 3.**
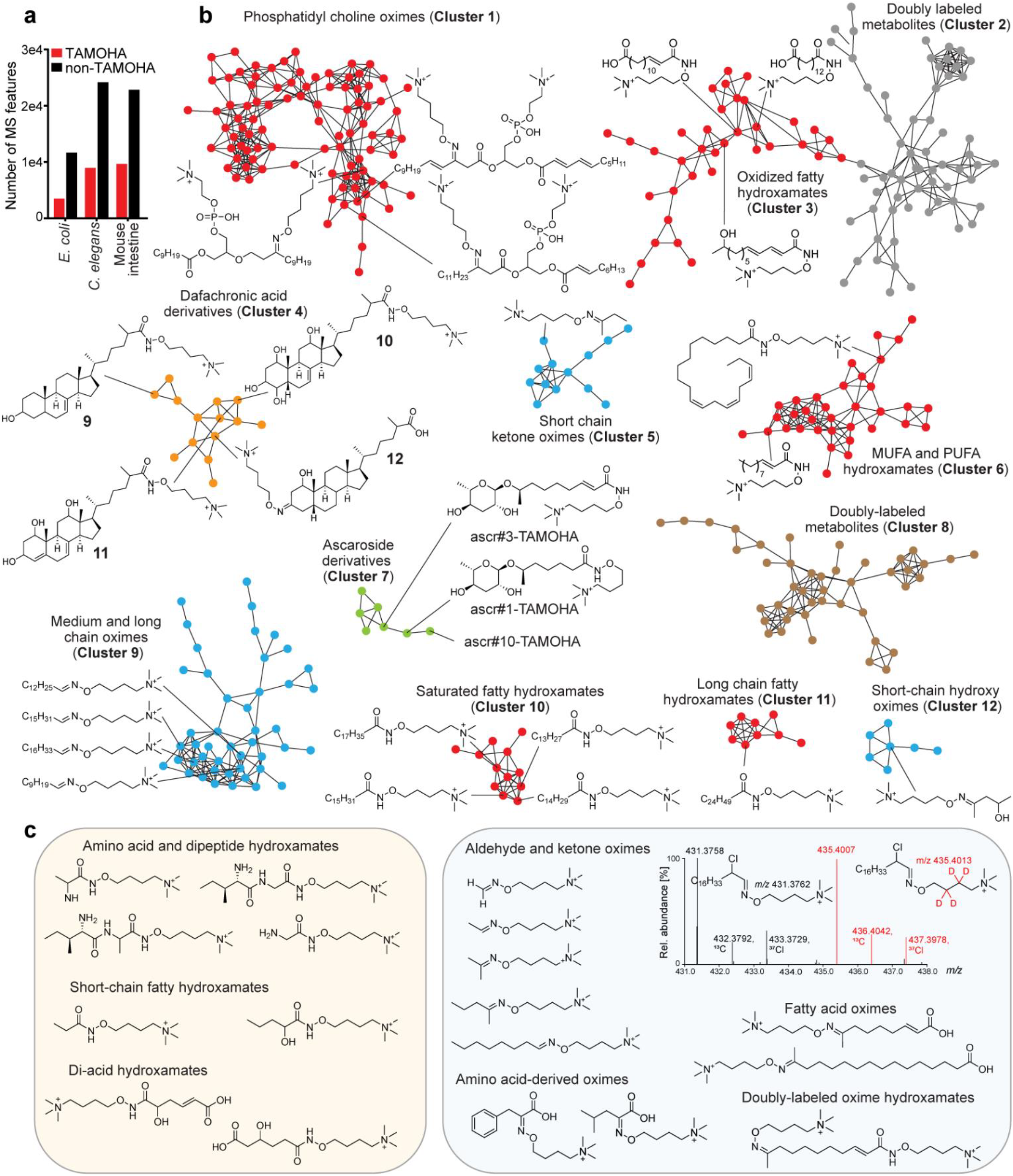
The scope of TAMOHA labeling. (a) Numbers of TAMOHA labeled mass features (red) identified in three different model organisms compared to non-TAMOHA labeled mass features (black). Mass features must be present in each of five biological replicates to be included in this quantification (see Methods). (b) Partial MS^2^ network of TAMOHA labeled metabolites. Shown structures were proposed based on analysis of the MS^2^ spectra and, if available, confirmed by comparison with authentic standards. See Figure S8 for the full MS^2^ network. (b) Additional examples for detected TAMOHA hydroxamates and oximes, including superimposed isotope patterns of mass spectra for the TAMOHA conjugate of α-chlorooctadecanal.

To gain insight in the chemical diversity of the TAMOHA-labeled metabolites, we acquired MS^2^ spectra for more than 1,000 TAMOHA conjugates detected in *C. elegans*. Networking parsed the MS^2^ spectra into more than 50 clusters (Fig. 3b). Analysis of representative MS^2^ spectra selected from several of the largest clusters indicated that they represent TAMOHA fatty hydroxamates, which could be derived, e.g., from acyl-thioester intermediates in fatty acid biosynthesis or β-oxidation. This includes several series of saturated straight chain and iso-branched fatty hydroxamates (Cluster 10, and see Fig. S7), mono- and poly-unsaturated fatty hydroxamates (Cluster 6), as well as hydroxamates of hydroxylated or otherwise oxidized fatty acids (Cluster 3).

In addition, we observed several MS^2^ clusters that correspond to TAMOHA oximes of long-chain ketones or aldehydes (Cluster 9), representing a facet of fatty acid metabolism that has been difficult to characterize but is increasingly recognized as supporting diverse signaling functions^28^. This includes short and long chain simple aliphatic oximes (Clusters 5 and 9, Fig, 3b, c) as well as diverse, more highly oxygenated species (Cluster 12, Fig. 3b). We further detected a series of α-chlorofatty aldehydes (Fig. 3c), electrophilic metabolites that in human and mouse are involved in neutrophil activation during immune responses^29^ but challenging to detect due to their generally low concentrations^30^.

We further detected a large MS^2^ cluster of doubly-charged species, representing TAMOHA oximes of phosphatidylcholines (Cluster 1). MS^2^ fragmentation spectra of these mass features revealed a diagnostic fragment ion at *m/z* 184.0733 corresponding to the phosphatidylcholine head group, in addition to the characteristic TAMOHA-derived fragments (Fig. 3b). TAMOHA thus allows to profile keto-phospholipids, another class of lipids that were recently shown to modulate diverse signaling pathways and influence inflammatory responses but have been difficult to track with conventional metabolomics^31^.

Another family of metabolites that serve as potent signaling molecules but due to low abundance or poor ionization properties are often difficult to detect and quantify is represented by keto-functionalized cholesterol derivatives, e.g., cholestanone derivatives and keto-functionalized bile acids. In *C. elegans*, 3-keto-functionalized cholestanoic acids are ligands of the nuclear hormone receptor DAF-12, a vitamin D and liver-X receptor homologue that plays a central role in larval development, fat metabolism, and lifespan in *C. elegans*^32^. Our MS^2^ networking revealed a prominent cluster sharing a 27-carbon skeleton with at least 5 degrees of unsaturation, consistent with TAMOHA derivatives of dafachronic acids (Cluster 4). Fragmentation patterns suggest that features in this cluster represent both oximes, derived from 3-keto-cholestanoic acids (e.g., **12**), as well as hydroxamates (e.g., **9, 10**, and **11**), indicating that corresponding thioesters may play a role in dafachronic acid degradation, e.g., via β-oxidation (Fig. 3b and S9a, b). Some of the mass features in this cluster appear to represent further oxidized dafachronic acid derivatives, hinting at additional yet unexplored pathways of steroid metabolism in *C. elegans*. We also detect doubly charged mass features likely derived from TAMOHA addition at both the 3-oxo position and the thioester terminus of a dafachronic acid-CoA conjugate (Fig. S9c).

Other MS^2^ clusters representing nematode-specific metabolite families include TAMOHA derivatives of ascarosides (Cluster 7), a family of signaling molecules that play a central role in regulating *C. elegans* development and behavior^33, 34^. These include ascr#1-TAMOHA, ascr#3-TAMOHA, and ascr#10-TAMOHA plausibly derived from the corresponding acyl-CoA thioesters, which are intermediates of ascaroside biosynthesis via peroxisomal β-oxidation (Fig. 3b)^35^. Targeted analysis revealed additional TAMOHA labeled ascarosides, of which we confirmed the structure of ascr#18-TAMOHA via coinjection with a synthetic sample (Fig. S10). Lastly, we detected TAMOHA conjugates of diverse thioesters, aldehydes, and ketones derived from non-lipid primary metabolism. For example, we detected mass features that correspond to TAMOHA hydroxamic acid derivatives of amino acids and di-peptides as well as TAMOHA oxime derivatives of intermediates in phenylalanine and leucine metabolism (Fig. 3b and S11).

Although we propose structures for many TAMOHA derivatives based on their HRMS and MS^2^ spectra, several hundred detected TAMOHA derivatives remain unidentified, hinting at a large reservoir of yet uncharacterized electrophilic species (see Fig. S8). Taken together, these data demonstrate that TAMOHA labels diverse electrophilic metabolites, enabling or simplifying characterization of molecular species that due to lack of chemical stability, low abundance, or poor ionization properties are difficult to detect with conventional MS-based metabolomics.

### Profiling Coenzyme-A esters in *E. coli*

Next, we aimed to demonstrate that TAMOHA can be applied in complex biological systems to detect changes in the production of specific electrophilic species, e.g., due to inactivation of enzymes involved in thioester metabolism. As an initial test for the utility of TAMOHA to investigate biochemical pathways, we selected fatty acid (FA) metabolism in *E. coli*. Fatty acyl-coenzyme-A thioesters (FA-CoAs) play a central role in FA biosynthesis and degradation, and thus we envisioned that TAMOHA could be used to profile changes in FA-CoA profiles in response to, e.g., mutations in enzymes involved in fatty acid metabolism. We started by comparing the profiles of free FAs and FA-CoAs in wildtype *E. coli* (BW25113). For this purpose, *E. coli* cultures were split into two samples, which were treated with either a solution of TAMOHA in PBS buffer or PBS buffer only (as a control), and were then analyzed by HPLC-HRMS.

MS^2^ analyses of TAMOHA-treated *E. coli* revealed several homologous series of mass features whose molecular formulae and fragmentation patterns suggested that they represent TAMOHA derivatives of electrophilic intermediates in FA biosynthesis or degradation, including putative TAMOHA conjugates of saturated, unsaturated, and hydroxylated FAs. We identified mass features consistent with TAMOHA conjugates of saturated FA ranging in length from C_3_ to C_24_, and we confirmed the identity of TAMOHA conjugates of saturated FAs from C_7_ to C_19_ via co-injection with authentic synthetic samples (Figs. 4a and S12a).

**Figure 4.**
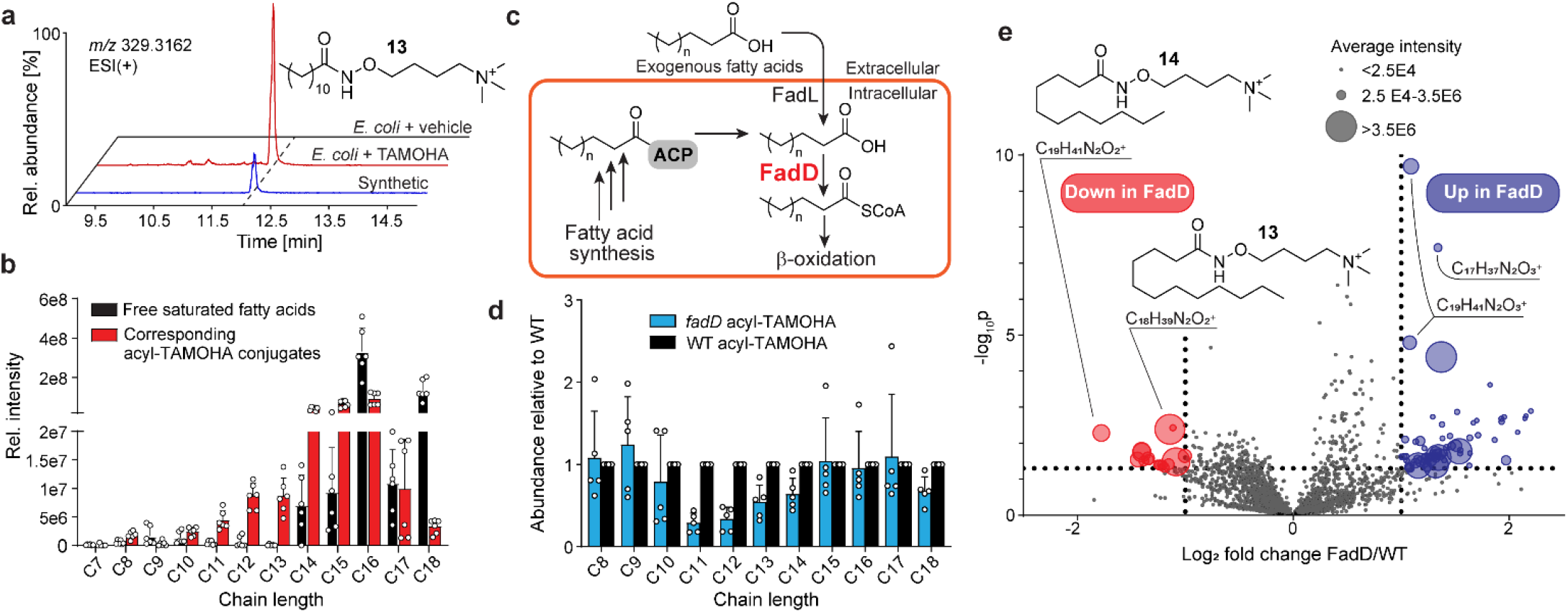
TAMOHA reveals substrate range of a fatty acid metabolism enzyme in *E. coli*. (a) Ion chromatograms representing TAMOHA-labeled dodecanoic acid obtained from *E. coli* treated with vehicle (top) or TAMOHA (middle), compared with a synthetic standard (bottom). (b) Profiles of saturated, straight chain fatty acids (black) compared to their TAMOHA-labeled conjugates (red) in *E. coli*. (c) Proposed role of FadD in *E. coli* fatty acid β-oxidation. (d) Relative abundances of TAMOHA-labeled straight chain, saturated fatty acids in FadD mutant and wildtype *E. coli*. (e) Volcano plot of mass features detected by HPLC-HRMS (ESI+) in TAMOHA-treated wildtype and FadD mutant *E. coli*.

Comparing the relative abundances of FA-TAMOHA conjugates to the corresponding free fatty acids revealed starkly different profiles. For example, free medium-chain FAs (C_10_-C_14_) were 10-100-fold less abundant relative to longer chained free FAs, whereas differences between the abundances of medium- and long-chain TAMOHA conjugates were much smaller (Fig. 4b). These observations indicate that pools of activated FAs are biased toward shorter chain lengths compared to the corresponding free FAs, which seems plausible, given that profiles of activated FAs include FA-CoA esters from various stages of chain-shortening via β-oxidation as well as fatty acylated Acyl Carrier Protein (ACP) from different stages of chain elongation during FA biosynthesis. In addition, comparison of relative abundances of the TAMOHA-conjugates and free FAs in *E. coli* revealed that the TAMOHA conjugate of pentadecanoic acid was similarly abundant as tetradecanoic and hexadecanoic acid (Fig. 4b). This high abundance of the TAMOHA-derivative of saturated C_15_ was surprising, given that odd saturated FAs are generally much less abundant than even-chain FAs in *E. coli* and most other bacteria^36^. Among TAMOHA conjugates of monounsaturated FAs, C18:1 was most abundant, derived from activated cis-vaccenic acid^37^. In addition, we detected C19:1 and C17:1 TAMOHA conjugates, which represent activated cyclopropyl-containing FAs derived from bacterial lactobacillic acid (LBA) (Fig. S12b).

These results demonstrate that TAMOHA treatment can provide a comprehensible inventory of fatty acyl CoA species. Next, we explored the potential of TAMOHA labeling for the *in vivo* study of FA metabolism enzymes. Acyl-CoA synthetases (ACSs) play a central role in metabolism by converting carboxylic acids into the corresponding acyl-CoA thioesters, activating them for subsequent chain extension, breakdown via β-oxidation, or as substrates for other acyl transferase enzymes^38, 39^. Importantly, the substrate scope of most ACSs in model systems and humans have only been characterized *in vitro*, but not *in vivo*^39-41^. We reasoned that combining TAMOHA labeling and untargeted comparative metabolomics of wildtype and ACS-deficient mutants would reveal putative *in vivo* substrates as TAMOHA-labeled features whose abundances are reduced or are abolished in the mutant compared to wildtype.

As a test case we selected the *E. coli* ACS, FadD, which has been reported to activate exogenous FAs for subsequent β-oxidation (Fig. 4c)^42, 43^. Previous *in vitro* studies indicate that FadD exhibits a preference for FAs ranging from C_10_ to 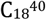, and that expression of FadD can rescue growth of *E. coli* supplemented with C_10_ FA as its sole carbon source^43^. To investigate the *in vivo* substrate scope of FadD, we treated wildtype and FadD mutant *E. coli* with TAMOHA, followed by comparative analysis via HPLC-HRMS to identify differential TAMOHA labeled FAs. Untargeted analysis revealed significantly reduced abundances of saturated TAMOHA labeled FAs from C_11_ to C_14_ in FadD mutants compared to wildtype (Figs. 4d, e and S12c), as well as accumulation of TAMOHA labeled C_10_ and C_12_ hydroxy FAs in FadD mutants (Fig. 4e). In contrast, profiles of unsaturated FA-TAMOHA conjugates were largely unchanged. These findings suggest that FadD exhibits a preference for saturated medium chain fatty acids from undecanoic to tetradecanoic acid *in vivo*. Although it is possible that compensatory metabolic processes work to maintain fatty acid homeostasis in the absence of FadD and thereby mask the effect of FadD deletion on other fatty acid chain lengths, our results show that TAMOHA can provide detailed insight in mutant-dependent changes in CoA ester profiles.

### TAMOHA corroborates substrate of a *C. elegans* β-oxidation enzyme

Encouraged by these results, we employed TAMOHA to probe a recently discovered branch of fat metabolism in *C. elegans*. In *C. elegans, acdh-11* is an unusual acyl-CoA dehydrogenase homolog that has been implicated in the metabolism of bacterial cyclopropyl fatty acids and thus is of great interest in the context of host microbe interactions. Cyclopropyl fatty acids are not endogenously produced by *C. elegans* but are abundant in *E. coli*, the standard laboratory food for *C. elegans*, and are also increasingly found in the human diet^44^. Previous work indicated that food-derived cyclopropyl fatty acids are chain-shortened via fatty acid β-oxidation, producing a β-cyclopropyl C_11_ fatty acid, named becyp#1 that is unsuitable for processing by the canonical β-oxidative machinery (Fig. 5a)^45^. Further metabolism of becyp#1 was shown to require *acdh-11*^45, 46^, based on the observation that diverse oxidized shunt metabolites of becyp#1 accumulate in *acdh-11* mutant animals. These shunt metabolites could plausibly be derived from accumulation of becyp#1-CoA as the putative substrate of ACDH-11.

**Figure 5.**
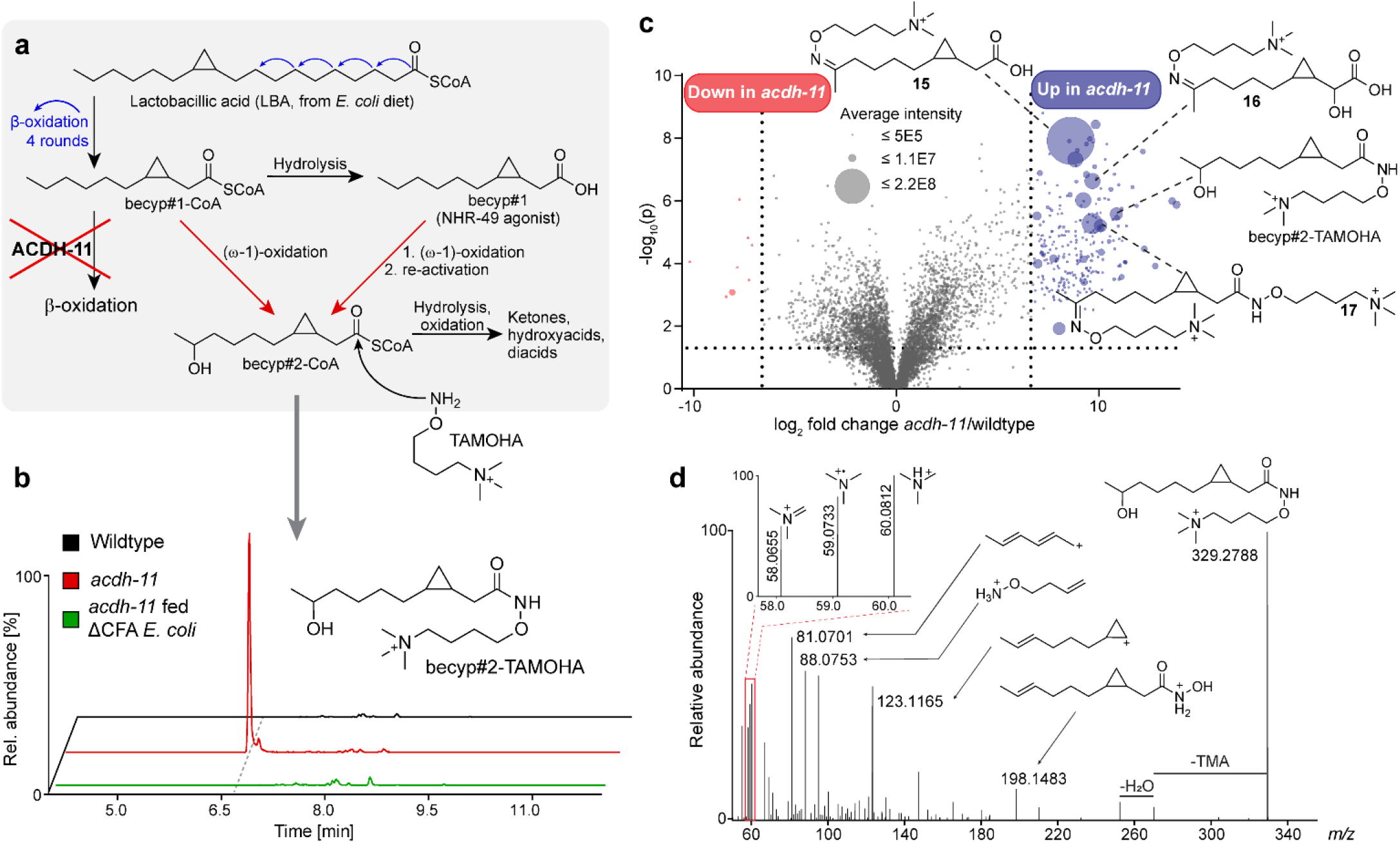
TAMOHA reveals putative substrate of the acyl-CoA dehydrogenase ACDH-11. (a) Current model for the metabolism of bacterial cyclopropyl fatty acids in *C. elegans* and the proposed role of ACDH-11 (b) Ion chromatograms of becyp#2-TAMOHA in wildtype (top), *acdh-11* mutant (middle), and *acdh-11* mutant *C. elegans* fed cyclopropane fatty acid synthase deficient *E. coli* (ΔCFA). (c) Volcano plot of mass features detected by HPLC-HRMS (ESI+) in TAMOHA-treated wildtype and *acdh-11* mutant *C. elegans*, highlighting features significantly increased (red) or decreased (blue) more than 100-fold in the *acdh-11* mutant. (d) MS^2^ fragmentation pattern of TAMOHA-labeled becyp#2.

To test whether the putative substrate of ACDH-11, becyp#1-CoA and perhaps related CoA species accumulate in *acdh-11* mutants, we applied TAMOHA treatment to *C. elegans* wildtype and *acdh-11* mutants, using two different *E. coli* strains as food, *E. coli* BW25113 (WT), and *E. coli* JW1653, a mutant that does not produce cyclopropane fatty acids (ΔCFA). Untargeted HPLC-HRMS analysis revealed a cluster of mass features enriched more than 100-fold in *acdh-11* mutant animals compared to wildtype. These differential features were absent in samples derived from animals raised on cyclopropane-deficient bacteria, indicating that they must be derived from cyclopropane fatty acids (Fig. 5b).

The MS^2^ spectra of the differential mass features indicated that they represent TAMOHA-derivatives of C_11_-FAs with at least one degree of unsaturation and varying oxygenation patterns, consistent with their origin from becyp#1 and becyp#1-CoA (Fig. 5c, d, S13, S14). Among the *acdh-11*-dependent mass features, becyp#1-TAMOHA was much less abundant than most oxygenated derivatives (Fig. S13), e.g., becyp#2-TAMOHA, **15, 16**, and **17**, suggesting that accumulating becyp#1-CoA is rapidly modified, e.g., by ω-hydroxylases^47^ to yield oxidized derivatives, consistent with previous studies^45^. Taken together, our results support that processing of C_11_-CoA substrates such as becyp#1-CoA by ACDH-11 is required for the metabolism of bacteria-derived cyclopropyl fatty acids in *C. elegans*.

### TAMOHA provides insights in the role of an acyl-CoA synthetase

Lastly, we employed TAMOHA to investigate the biosynthetic role of the *C. elegans* acyl-CoA-synthetase ACS-7, which is required for the production of the aggregation pheromone, icas#9^48, 49^ (Fig. 6a) In *acs-7* mutants, icas#9 production is abolished, whereas levels of icas#10, which differs only in the length of the hydrocarbon side chain, are increased^48^. Further, it was shown that icas#9 is derived from icas#10 via two rounds of peroxisomal β-oxidation, which shortens the 9-carbon side chain of icas#10 to 5 carbons as in icas#9^49^. In turn, icas#10 is derived from attachment of the indolecarboxyl moiety to ascr#10 via the lysosomal carboxylesterase CEST-3^34^. To explain abolishment of icas#9 production in *acs-7* mutants, it was proposed that ACS-7 converts icas#10 to icas#10-CoA for subsequent peroxisomal β-oxidation to afford icas#9-CoA^49^ (Fig. 6a). Therefore, we expected to find that in *acs-7* mutants, levels of icas#10-CoA and icas#9-CoA would be reduced or abolished.

**Figure 6.**
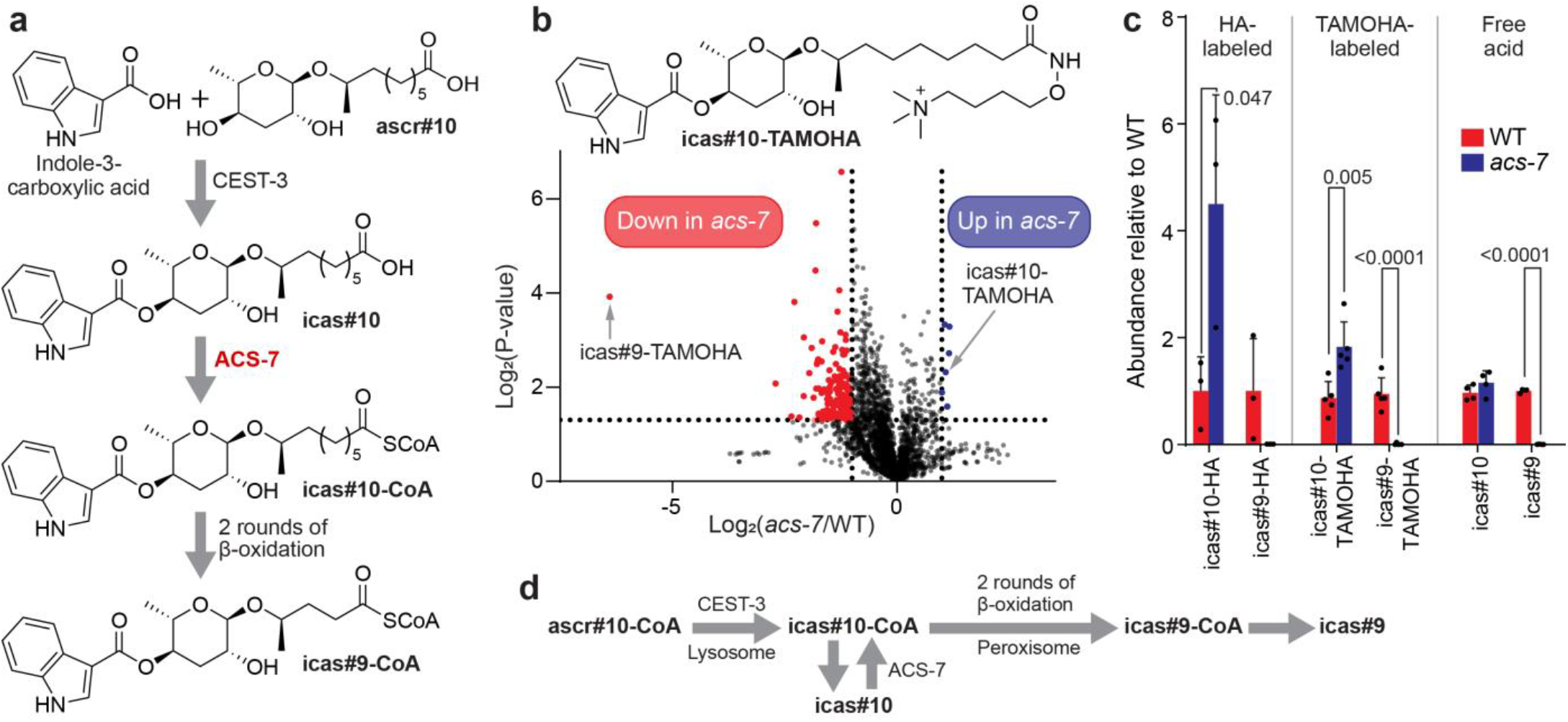
TAMOHA-based analysis of the role of the acyl-CoA synthetase ACS-7 in ascaroside pheromone biosynthesis. (a) Current model for the biosynthesis of the aggregation pheromone icas#9^48, 49^. (b) Volcano plot of mass features detected by HPLC-HRMS (ESI+) in TAMOHA-treated wildtype and *acs-7* mutant *C. elegans*, highlighting features significantly decreased (red) or increased (blue) more than 10-fold in *acs-7* mutants. (c) Relative abundances of icas#9 and icas#10 as well as their HA- and TAMOHA-labeled conjugates in wildtype and *acs-7* mutants. P-values were calculated using a two-tailed Student’s *t*-test. (d) Alternative model for icas#9 biosynthesis.

To test this, we treated wildtype and *acs-7* mutant cultures with TAMOHA, followed by HPLC-HRMS analysis as above. Targeted analyses of TAMOHA-treated samples revealed the presence of both icas#9-TAMOHA and icas#10-TAMOHA in wildtype, and almost complete abolishment of icas#9-TAMOHA in *acs-7* mutants, as expected. However, abundances of icas#10-TAMOHA were increased relative to wildtype in *acs-7* mutants, not reduced (Fig. 6b, and S15). To confirm these unexpected results, we additionally compared wildtype and *acs-7* mutants using hydroxylamine (HA) instead of TAMOHA, which revealed the same trend, i.e., abolishment of icas#9-HA but increased levels of icas#10-HA (Fig. 6c), although significance was lower than in the TAMOHA study, likely due to the worse peak shapes and ionization of the HA derivatives. Since levels of activated icas#10 are not abolished in *acs-7* mutants, as suggested by current icas#9 biosynthesis models^49^, but instead increased, there must exist other mechanisms for producing activated icas#10. This may involve activation by other acyl-CoA synthetases, e.g., in the cytosol. Alternatively, CEST-3 may operate on ascr#10-CoA, not free ascr#10, and thus icas#10-CoA could represent the direct product of CEST-3 (Fig. 6d). Distinguishing between these possibilities will require further study; however, our results demonstrate that TAMOHA is a useful tool for tracking biosynthetic intermediates and hypothesis generation.

## DISCUSSION

Metabolism represents a complex network of thousands of interconnected reactions, many of which involve the generation of transiently stable, reactive electrophiles. Detailed knowledge of their structures and abundances is of central importance for understanding metabolic pathways, but due to their inherent reactivity most electrophilic metabolites do not survive routine metabolomic analyses. Moreover, some electrophilic species such as aliphatic aldehydes and ketones do not ionize well, further complicating their detection via mass spectrometric analyses. The TAMOHA probe addresses this limitation by (i) trapping activated metabolites as stable derivatives, (ii) enhancing sensitivity by attaching a constitutively charged moiety, and (iii) providing a unique MS^2^ fingerprint for identification. Our model studies demonstrate that TAMOHA reacts with CoA thioesters, aldehydes, and ketones, but not with esters and provides as much as 1000-fold sensitivity enhancement. We further show that TAMOHA facilitates comprehensive and sensitive profiling of short-chain aldehydes and ketones, which due to their volatility and reactivity are difficult to track in complex biological samples.

TAMOHA labeling of samples from *E. coli, C. elegans*, and mouse revealed a vast space of structurally diverse TAMOHA conjugates, derived from all corners of metabolism. Our results thus suggest that conventional metabolomics captures a rather incomplete picture of chemical diversity in living systems. While many of the detected TAMOHA conjugates could be assigned to known biochemical pathways, it should be noted that we confirmed the identities of only a few representative examples with authentic standards, and that, based on their MS^2^ spectra, for several hundred of the detected TAMOHA conjugates it was not obvious to what biochemical context they may belong, suggesting significant discovery potential for metabolism and biosynthesis studies.

Further, we show that TAMOHA can be an effective tool to study enzyme function *in vivo*. TAMOHA treatment of an *E. coli* mutant of the ACS enzyme FadD^42, 43^ confirmed the proposed function of this enzyme, and further showed that the profile of activated fatty acids is distinct from the corresponding profile of free fatty acids in wildtype *E. coli*. Similarly, the results from TAMOHA treatment of *C. elegans* mutants of the acyl-CoA dehydrogenase *acdh-11*^45, 46^ support the proposed function of this enzyme in the degradation of bacteria derived β-cyclopropyl fatty acids, providing a striking example for the adaptation of host biosynthetic machinery to enable processing of microbiota-derived lipids that were recently shown to regulate fat metabolism via conserved nuclear receptor signaling^45^. In contrast, our results for the acyl-CoA synthetase *acs-7* were unexpected and suggest that current models for the biosynthesis of modular ascarosides are incomplete. More work is required to clarify the origin of icas#10-CoA and the role of *acs-7* in icas#9 biosynthesis^50^; however, this example demonstrates that using TAMOHA to measure the abundance of biosynthetic intermediates *in vivo* can provide unexpected insights that complement results from conventional metabolomics and *in vitro* studies of enzyme function.

The above examples suggest that TAMOHA can be applied to profile electrophilic species in a wide range of biological contexts, e.g., to elucidate biosynthetic functions, uncover previously invisible sets of metabolites, or to investigate flux through metabolic pathways and how stable metabolic end products and electrophilic intermediates are differentially regulated. In addition, TAMOHA may enable targeted approaches to investigate electrophilic. protein post translational modifications (PTMs, Fig. 7). In a pilot study, we recently showed that TAMOHA can be used to profile protein iminylation derived from the reaction of the ε-amino group of lysine with cellular aldehydes^51^ (Fig. 7). Furthermore, TAMOHA may facilitate profiling and discovery of other types of electrophilic PTMs^52^, e.g., thioe-ster-linked PTMs, including biosynthetic intermediates on polyketide synthetases or non-ribosomal peptide synthases^53^.

**Figure 7.**
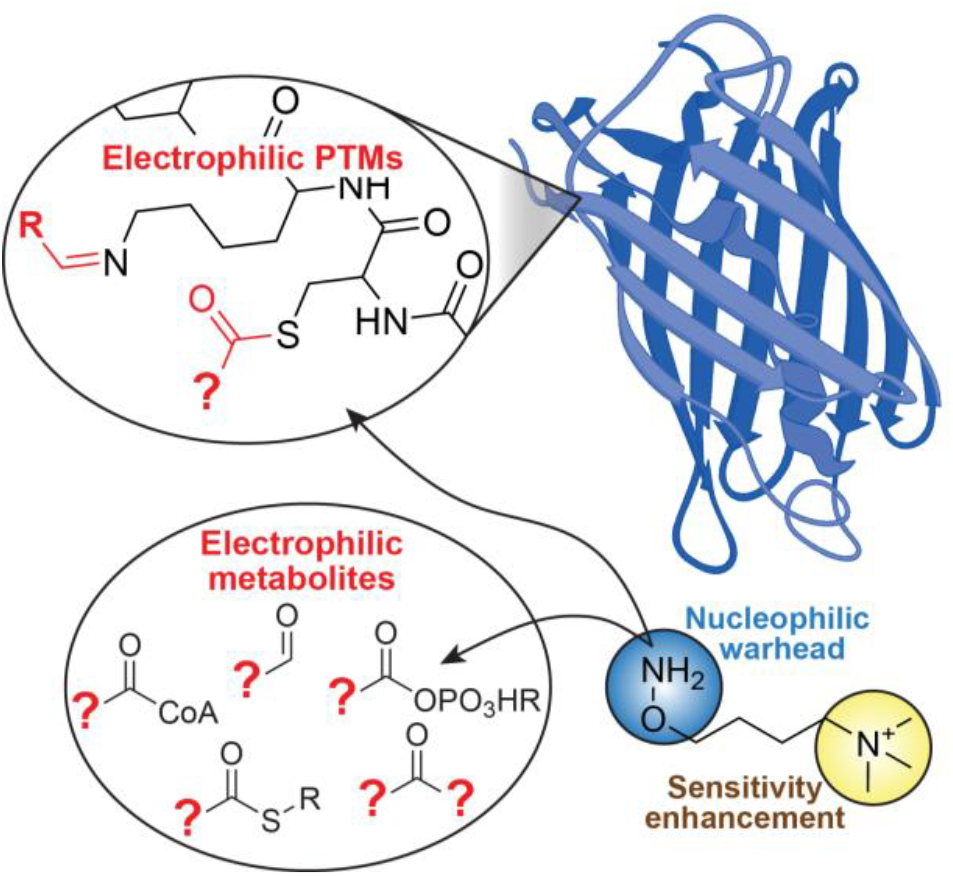
In addition to applications in metabolomics, TAMOHA can be employed to trap electrophilic PTMs, e.g., protein iminylation derived from reaction of lysine with aliphatic aldehydes (R = methyl, ethyl, propyl, isopropyl)^51^. In addition, TAMOHA may facilitate characterization of thioester-based PTMs.

## ASSOCIATED CONTENT

### Supporting Information

This material is available free of charge at https://pubs.acs.org

General methods, TAMOHA treatment conditions, methods for metabolomics, synthetic procedures with NMR spectroscopic data, Figures S1-S16, NMR spectra.

## AUTHOR INFORMATION

### Author Contributions

All authors have given approval to the final version of the manuscript. ‡These authors contributed equally.

### Notes

F.C.S. is a cofounder of Ascribe Bioscience and Holoclara Inc. The other authors declare no competing interests.

## ACKNOWLEDGMENT

Some strains used in this work were provided by the CGC which is funded by the NIH Office of Research Infrastructure Programs (P40 OD010440). We thank Ivan Keresztes and Dimitrios Koumoulis for assistance with NMR spectroscopy. Ascr#18 was provided as a gift Ascribe Bioscience.

## ABBREVIATIONS

CoA: coenzyme A
HA: hydroxylamine
ESI+: positive electrospray ionization mode
TAMOHA: *O*-(trimethylammoniobutyl) hydroxylamine
CuAAC: copper catalyzed azide-alkyne cycloaddition
HPLC: high pressure liquid chromatography
HRMS: high resolution mass spectrometry
PFBHA: pentafluorobenzyl)hydroxylamine
FA: fatty acid
ACP: acyl carrier protein
LBA: lactobacillic acid
ACS: acyl-CoA synthase
ΔCFA: cyclopropane fatty acid synthase deficient bacteria
WT: wildtype
PTM: post translational modification
NRPS: non-ribosomal peptide synthase.

