## Supporting Information for "Omics-scale capture of electrophilic metabolites"

[1]Boyce Thompson Institute and Department of Chemistry and Chemical Biology, Cornell University, Ithaca, New York 14853, United States. [2]Division of Nutritional Sciences, Cornell University, Ithaca, New York 14853, United States. [3]Jill Roberts Institute for Research in Inflammatory Bowel Disease, Division of Gastroenterology and Hepatology, Joan and Sanford I. Weill Department of Medicine, Weill Cornell Medicine, New York, NY, United States.

### Table of Contents

**1. Methods** 2
**2. Synthetic procedures** 6
**2.1. General synthetic methods** 6
**2.2 Synthesis of TAMOHA and TAMOHA-$D_4$** 7
**2.3 TAMOHA quality assurance** 9
**2.4 Synthesis of TAMOHA derivatives** 9
**3. Supporting Figures** 11
**4. NMR spectra appendix** 27
**5. Supporting references** 46

### 1. Methods

**General information.** Unless noted otherwise, reagents were purchased from Sigma-Aldrich. *C. elegans* metabolites are referred to by their SMID identifiers (a search-compatible Small Molecule IDentifier) e.g., "icas#9" or "bemeth#1". The SMID database (www.smid-db.org) is an electronic resource maintained in collaboration with WormBase (www.wormbase.org). A complete list of SMIDs can be found at www.smid-db.org/browse.

**Bacterial strains and maintenance.** Bacteria were maintained on sealed Lennox Broth (LB) agar plates supplemented with appropriate antibiotics and stored at 4 °C for up to 4 weeks. The following bacterial strains were used, *E. coli* OP50, BW25113 (wildtype), JW1653 (Δcfa), and JW1794 (FadD). BW25113, JW1653, and JW1794 were purchased from the Keio collection of *E. coli* knockouts managed by Horizon Discovery. *E. coli* OP50 was maintained with supplemental streptomycin, JW1653 and JW1794 were maintained with supplemental kanamycin.

***C. elegans* strains and maintenance.** Unless noted otherwise, worms were maintained on Nematode Growth Medium (NGM) 6 cm diameter petri dish plates seeded with *E. coli* OP50, under standard growth conditions[1]. The following nematode strains were used in comparative metabolomics experiments: *C. elegans* Bristol N2 ("Wildtype"), *C. elegans* DMS441 *acdh-11(n5878)*, *C. elegans* FCS-10 *acs-7(tm6781)*.

***E. coli* suspension for nematode cultures.** *E. coli* was streaked on Lennox Broth (LB) agar plates and grown at 37 °C for 20 hours. Plates were then sealed and stored at 4 °C for up to 4 weeks. Liquid cultures were generated by inoculating a single colony into 125 mL Erlenmeyer flasks containing 30 mL LB. Cultures were grown at 37 °C shaking at 200 RPM for 20 hours. 10 mL of this culture was diluted in 2 L LB in a 4 L Erlenmeyer flask, and the resultant culture was grown at 37 °C shaking at 180 RPM for 20 hours. Bacteria were collected by centrifugation (2500xg, 4 °C, 20 min), supernatant was removed, and the bacterial pellet was resuspended in M9 solution to yield a 1 g/mL suspension of bacteria in M9.

***C. elegans* liquid cultures.** 6 cm maintenance plates of *C. elegans* were transferred to 10 cm diameter petri dish plates seeded with *E. coli* OP50 and grown to confluence. Confluent plates were washed with 5 mL S-complete medium into a 125 mL Erlenmeyer flask with 20 mL S-complete medium and fed with 2 mL of 1 g/mL *E. coli* suspension in M9. Mixed stage cultures were grown for 72 hours at 20 °C shaking at 180 RPM. After 72 hours, mixed stage cultures were pelleted, washed with 25 mL water, and treated with alkaline bleach to yield a suspension of eggs, which were washed two times with 25 mL M9 buffer and rocked overnight in 5 mL M9 solution to yield synchronized, starved L1 larvae. Cultures of synchronized adults were prepared by adding 70,000 synchronized L1 larvae, obtained from alkaline bleach treatment, to 25 mL of S-complete medium in a 125 mL Erlenmeyer flask to which was added 2 mL of 1 g/mL *E. coli* suspension in M9. Cultures were incubated at 20 °C shaking at 180 RPM for 60-63 hours until animals were gravid.

**TAMOHA model reactions.** Myristoyl coenzyme A ester, myristoyl methyl ester, or 2-octanone was solubilized in methanol at a concentration of 1 mM. To aliquots of these solutions, equivalent volumes of a 1 mM (1:1 treatment), 10 mM (1:10 treatment), or 100 mM (1:100 treatment) solution of TAMOHA in methanol was added at room temperature. Addition of methanol served as a control. These reactions were immediately sonicated using a Qsonica Q700 equipped with a horn cup (Process time = 2 min, Amplitude = 100, Pulse-ON time = 2 s, Pulse-OFF time = 2 s), diluted, cooled to 0 °C until analysis by high pressure liquid chromatography coupled to high-resolution mass spectrometry (HPLC-HRMS) on the same day.

**TAMOHA and PFBHA comparison.** To 100 µL of a 20 µM solution of acetone in water were added 100 µL of a 0.1 M solution of either TAMOHA or PFBHA in phosphate buffer at pH 4 and room temperature. Addition of water served as a control. Reactions were incubated overnight at room temperature, serially diluted, and stored at 2 °C until analysis by HPLC-HRMS on the same day. Analogous reactions using formaldehyde instead of acetone yielded the expected formaldehyde-

TAMOHA conjugate, whereas only trace amounts of the formaldehyde-PFBHA conjugate were detected by HPLC-HRMS.

**TAMOHA treatment of bacterial cultures.** *E. coli* was streaked on LB agar plates and grown at 37 °C for 20 hours. Plates were then sealed and stored at 4 °C for up to 4 weeks. Liquid cultures were generated by inoculating a single colony into 125 mL Erlenmeyer flasks containing 30 mL LB. Cultures were grown at 37 °C shaking at 200 RPM for 20 hours. Bacterial cultures were split into 5 mL aliquots and centrifuged (2850xg, 4200 °C, 25 min) using a Beckman Coulter Allegra X-15R Centrifuge. Supernatant was removed, and bacterial pellets were suspended in either 1 mL of 0.1 M TAMOHA in 0.1X phosphate-buffered saline (PBS), pH = 4.0 or 1 mL of 0.1X PBS alone. Samples were sonicated using a Qsonica Q700 equipped with a horn cup (Process time = 2 min, Amplitude = 100, Pulse-ON time = 2 s, Pulse-OFF time = 2 s), immediately snap frozen in liquid nitrogen, and stored at –20 °C until analysis by HPLC-HRMS. To ensure that TAMOHA is used at a concentration that would trap electrophilic metabolites quantitatively in a complex biological matrix, we treated *E. coli* cultures with a range of different TAMOHA concentrations (Fig. S14). We determined that treatment with 20 mM TAMOHA maximizes electrophilic metabolite trapping, but is low enough so that remaining unreacted TAMOHA does not interfere with subsequent HPLC analysis.

**Mouse tissue harvest for TAMOHA analysis.** Fresh mouse ileum samples were prepared as described[2]. All additional mice were housed in specific pathogen-free conditions within the Cornell University Weill Hall Barrier Facility and handled in accordance with Cornell Institutional Animal Care and Use Committee (IACUC) protocol 2023-0054. $Adh5^{-/-}$ mice ($Adh5^{tm1Stam}$; MGI ID: 3033711) maintained in C57BL/6J background were kindly gifted from Dr. Jonathan Stamler (Liu *et al.*, 2004). Male $Adh5^{-/-}$ mice aged 7.5 to 14.5 weeks old received a single intraperitoneal injection with either 1.5 g/kg methanol-$d_4$ (Oakwood Chemical) prepared in 0.9% saline or 0.9% saline control. Following 24 hours, the mice were euthanized by $CO_2$ exposure and terminal venepuncture. Liver and kidney were collected into cryovials and immediately snap frozen on dry ice for downstream analysis. Samples were stored at -80 °C until TAMOHA treatment.

**TAMOHA treatment of mouse illeum.** Approximately 100 mg of frozen sections of ileum were powdered on dry ice using a mortar and pestle, and the resultant powder was partitioned into two Eppendorf tubes. 0.25 mM TAMOHA in 0.1X PBS at pH = 4 or 0.1X PBS only were added at 20 µL per mg of tissue. Samples were then sonicated using a Qsonica Q700 equipped with a horn cup (Process time = 2 min, Amplitude = 100, Pulse-ON time = 5 s, Pulse-OFF time = 2 s) and concentrated to dryness in an SC250EXP Speedvac Concentrator coupled to an RVT5105 Refrigerated Vapor Trap (Thermo Scientific). The resulting powder was suspended in 20 µL of methanol per mg of tissue and extracted for 18 hours at 4 °C with gentle rocking. After extraction, samples were centrifuged (2500Xg, 10 min, 20 °C), the clarified extract was transferred to a 4 mL scintillation vial, and samples were concentrated to dryness in an SC250EXP Speedvac Concentrator coupled to an RVT5105 Refrigerated Vapor Trap (Thermo Scientific). The resulting powder was suspended in 1 µL of methanol per mg of tissue by vigorous vortex and 10 minutes of sonication (Mettler Electronics Cavitator ultrasonic cleaner). The resulting suspension was transferred to a 1 mL Eppendorf tube, centrifuged (10,000Xg, 5 min, 20 °C) in an Eppendorf 5417 R centrifuge and the clarified extract was transferred to HPLC vial inserts (SureSTART 6PME03C1SSP). Samples were stored at -20 °C until analysis.

**TAMOHA treatment of mouse Liver and Kidney.** Approximately 50 mg of frozen liver, or 20 mg of frozen kidney samples were powdered on dry ice using a mortar and pestle, and the resultant powder was partitioned into two Eppendorf tubes. 0.25 mM TAMOHA in 0.1X PBS at pH 4 or 0.1X PBS only were added at 10 µL per mg of tissue. Samples were then sonicated using a Qsonica Q700 equipped with a horn cup (Process time 2 min, Amplitude 100, Pulse-ON time 5 s, Pulse-OFF time 2 s), centrifuged (10,000Xg, 5 min, 20 °C) in an Eppendorf 5417 R centrifuge and the clarified extract was transferred to HPLC vial inserts (SureSTART 6PME03C1SSP). Samples were stored at -20 °C until analysis.

**Synthesis of TAMOHA fatty acid standards.** Straight chain, saturated fatty acids from $C_7$ to $C_{19}$ (5 mg each) were combined and solubilized in 1 mL dichloromethane (DCM). This mixture was diluted 1:10 in DCM, and 4-(*N*,*N*-dimethyl)aminopyridine (DMAP, 0.303 mmol), *N*-(3-Dimethylaminopropyl)-*N*'-ethylcarbodiimide (EDC 0.303 mmol), and TAMOHA (0.303 mmol) were added. The reaction was left at room temperature for 18 hours then diluted in methanol and injected on HPLC−HRMS.

**Synthesis of ascr#18-TAMOHA.** ascr#18 was prepared as described[3]. ascr#18 (1.5 mg, 0.0045 mmol) was solubilized in 1.5 mL DCM and EDC (0.0067 mmol, 1.5 eq.) was added. The reaction was left at room temperature for 15 minutes. Now, TAMOHA (**1**, 0.0067 mmol, 1.5 eq.) and DMAP (0.0067 mmol, 1.5 eq.) were added as a solution in EtOH (0.5 mL). The reaction was left at room temperature for 30 minutes and concentrated *in vacuo*. After flash column chromatography on silica, fractions were analyzed by HPLC-HRMS.

**TAMOHA treatment of *C. elegans* liquid cultures.** Cultures of 70,000 synchronized gravid adult worms were transferred to 50 mL falcon tubes and settled for ten minutes. 10 mL of supernatant was transferred to a separate 50 mL falcon tube, snap frozen in liquid nitrogen, and lyophilized to dryness (exo-metabolome). The remaining worm pellet was suspended in 25 mL 0.1X PBS solution and split equally into two 50 mL falcon tubes. Now, each pellet was washed two more times with 0.1X PBS. After washing, 1 mL of 0.1M TAMOHA in 0.1X PBS, pH = 4.0 was added to one of the paired worm pellets, addition of 1 mL of 0.1X PBS served as a control. Samples were then sonicated using a Qsonica Q700 equipped with a horn cup (Process time = 2 min, Amplitude = 100, Pulse-ON time = 2 s, Pulse-OFF time = 2 s), and following sonication, each sample was snap frozen in liquid nitrogen (endo-metabolome) and stored at -20 °C until sample preparation.

***C. elegans* and *E. coli* sample preparation for HPLC-HRMS.** Snap frozen samples were lyophilized to dryness using a VirTis BenchTop 4 K Freeze Dryer. The resultant powder was suspended in 10 mL methanol and samples were allowed to rock at room temperature for 18 hours. After extraction, samples were centrifuged (2500Xg, 10 min, 20 °C), the clarified extract was transferred to a 20 mL scintillation vial, and samples were concentrated to dryness in an SC250EXP Speedvac Concentrator coupled to an RVT5105 Refrigerated Vapor Trap (Thermo Scientific). The resulting powder was suspended in 1 mL of methanol by vigorous vortex and 10 minutes of sonication (Mettler Electronics Cavitator ultrasonic cleaner). The resulting suspension was transferred to a 1 mL Eppendorf tube, centrifuged (10,000Xg, 5 min, 20 °C) in an Eppendorf 5417 R centrifuge and the clarified extract was transferred to an HPLC vial (Thermo Scientific cat. no. 6PSV9-1P). Samples were concentrated to dryness as above, suspended in 100 µL of methanol with vigorous vortex, transferred to a 1 mL Eppendorf tube, centrifuged (18,000Xg, 10 min, 20 °C) in an Eppendorf 5417 R centrifuge, and 70 µL was transferred to HPLC vial inserts (SureSTART 6PME03C1SSP). Samples were stored at -20 °C until analysis.

**Mass spectrometric analysis.** HPLC−HRMS analysis was performed on a ThermoFisher Scientific Vanquish Horizon UHPLC System controlled by Chromelion software (Thermo Fisher Scientific) coupled with a Thermo Q Exactive HF hybrid quadrupole-orbitrap high-resolution mass spectrometer, controlled by the same software, and equipped with a HESI ion source. Metabolites were separated using a water-acetonitrile gradient on an Agilent Zorbax Eclipse XDB-C18 column (150 mm × 2.1 mm, particle size 1.8 µm) maintained at 40 °C. Solvent A: 0.1% formic acid in water; Solvent B: 0.1% formic acid in acetonitrile. A/B gradient started at 1% B for 5 min after injection and increased linearly to 99% B at 20 min, using a flow rate 0.5 mL/min. Mass spectrometer parameters: spray voltage 3.0 kV, capillary temperature 380 °C, probe heater temperature 300 °C; sheath, auxiliary, and spare gas 60, 20, and 2, respectively; S-lens RF level 50, resolution 240,000 at *m/z* 200, AGC target $3\times10^6$. The instrument was calibrated with positive and negative ion calibration solutions (ThermoFisher). Each sample was analyzed in positive and negative modes using an *m/z* range 70 to 1000. Tandem mass spectrometry parameters: Resolution 45,000, AGC target $5\times10^4$, Maximum IT 80 ms, loop count 10, TopN 10, isolation window 1.0 *m/z*, stepped NCE 25 to 50, minimum AGC target $8\times10^3$, intensity threshold $1\times10^5$, no apex trigger, no charge exclusion, no peptide match, exclud18he isotopes on, dynamic exclusion 1.0 s, if idle, do not pick others. HPLC-HRMS RAW data were converted into mzXML file format using MSConvert (v.3.0, ProteoWizard) and were analyzed using Metaboseek v.0.9.9[4].

**Quantifying TAMOHA dependent mass features.** MzXML files were filtered using Metaboseek[4] to include mass features with a mean intensity of at least 1e4, that has a mass to charge ratio greater than *m/z* = 150, that were at least ten-fold more abundant in TAMOHA treated samples than PBS treated samples, and that scored between 0.9 and 1 using the Metaboseek fast peak shapes analysis. These filtering conditions were defined as TAMOHA dependent mass features. This was compared to mass features detected in PBS controls. Mass features in PBS controls were filtered for those with a mean intensity of at least 1e4, that were ten-fold more abundant in PBS treated samples compared to methanol blank samples, and that scored between 0.9 and 1 using the Metaboseek fast peak shapes analysis. Of the remaining mass features, only features present in all five TAMOHA-treated biological replicates and absent in all five PBS-treated control samples were included in the mass feature quantification in Fig. 3a.

**Generation of volcano plots.** MzXML files were filtered using Metaboseek[4] to include mass features with a mean intensity threshold of 5e4 in all replicates of at least one TAMOHA treatment condition, that had a mass to charge ratio between *m/z* 150 and 1200, that had a retention time on C18 between 180 s and 1600 s, and that were at least five-fold more abundant in TAMOHA treated samples than in non-TAMOHA treated samples. P-values were calculated using a two-sample independent Student's *t*-test, two tailed, assuming equal variance. P-values and fold over control were then log transformed and plotted using GraphPad Prism 9.0.0. P-value of $P \leq 0.05$ was used as a significance threshold. The most significant and differential mass features were then individually inspected and artefacts were removed. The volcano plot in figure 5 was generated by filtering MzXML files using the Metaboseek software to include mass features with a mean intensity of 9e3 in all replicates of at least one TAMOHA treatment condition, that had a mass to charge ratio between *m/z* 150 and 1200, that had a retention time on a reversed phase (C18) HPLC column between 180 s and 1600 s, that were at least ten-fold more abundant in TAMOHA treated samples than in TAMOHA $D_4$ treated samples, and that scored between 0.98 and 1 using the Metaboseek fast peak shapes analysis. P-values were calculated using a two-sample independent Student's *t*-test, two tailed, assuming equal variance. P-values and fold over control were then log transformed and plotted using GraphPad Prism 9.0.0. P-value of $P \leq 0.05$ was used as a significance threshold. Differential mass features were then individually inspected and mass features that did not bear characteristic TAMOHA fragments, or that were present in PBS controls, were removed.

**$MS^2$ molecular networking.** $MS^2$ spectra were converted to mzXML file using MSConvertGUI. Then mzXML files were uploaded to gnps.ucsd.edu for networking. Networking parameters: PAIRS MIN COSINE 0.5, ANALOG SEARCH 0, tolerance.PM tolerance 2.0, tolerance.Ion tolerance 0.5, MIN MATCHED PEAKS 4, TOPK 10, CLUSTER MIN SIZE 2, MAXIMUM COMPONENT SIZE 100, MIN PEAK INT 0.0, FILTER STDDEV PEAK INT 0.0, RUN MSCLUSTER on, FILTER PRECURSOR 8 WINDOW 1, FILTER LIBRARY 1, WINDOW FILTER 1, SCORE THRESHOLD 0.5, MIN MATCHED PEAK SEARCH 4. MAX SHIFT MASS 100. After networking was complete, the file was processed with Cytoscape software.

.

### 2. Synthetic procedures

#### 2.1. General synthetic methods

All oxygen and moisture-sensitive reactions were carried out under argon (Ar) atmosphere in flame-dried glassware. Solutions and solvents sensitive to moisture and oxygen were transferred via standard syringe and cannula techniques. Reactions were cooled with ice water or heated with mineral oil baths depending on reaction temperature. Mixtures (reaction or from chromatography) were concentrated using a Buchi rotary evaporator. Unless stated otherwise, all chemicals were purchased from Sigma Aldrich. 4-(*N,N*-dimethyl)aminopyridine (DMAP), *N*-(3-dimethylaminopropyl)-*N'*-ethylcarbodiimide (EDC) and dioxane was purchased from TCI. Methanolic ammonia (7N) was purchased from Acros Organics. Oleic acid was purchased from Cayman Chemicals. 4-Methylmorpholine was purchased from Fluka Chemical Corporation. Acetonitrile (ACN), water ($H_2O$), dichloromethane (DCM), ethyl acetate (EtOAc), hexanes, methanol (MeOH), dimethylformamide (DMF), were purchased form Fisher Scientific. Thin layer chromatography (TLC) was performed using J. T. Baker Silica Gel IB2F plated with analysis via UV and *p*-anisaldehyde, phosphomolybdic acid, and potassium permanganate stains. Flash chromatography was performed using Teledyne Isco CombiFlash systems and Teledyne Isco RediSep Rf silica and C18 columns. All deuterated solvents were purchased from Cambridge Isotopes. Nuclear Magnetic Resonance (NMR) spectra were recorded on Varian INOVA 600 (600 MHz) or Bruker (500 MHz) AVIII with BBO Prodigy cryoprobe spectrometer at Cornell University's NMR facility. $^{1}H$ NMR chemical shifts are reported in ppm (δ) relative to residual solvent peaks (3.31 ppm for methanol-$d_4$, 4.79 ppm for $D_2O$, and 7.26 ppm for chloroform-*d*). $^{13}C$ NMR chemical shifts are reported in ppm (δ) relative to residual solvent peaks (49.00 ppm for methanol-$d_4$, 49.00 ppm for residual methanol signal for $D_2O$, and 77.2 ppm for chloroform-*d*). All NMR data processing was done using MNOVA 15.0.0 (https://mestrelab.com/).

### 2.2 Synthesis of TAMOHA and TAMOHA-$D_4$

#### 2.2.1 Synthesis of 4-(aminooxy)-*N,N,N*-trimethylbutan-1-aminium bromide (TAMOHA, 1)

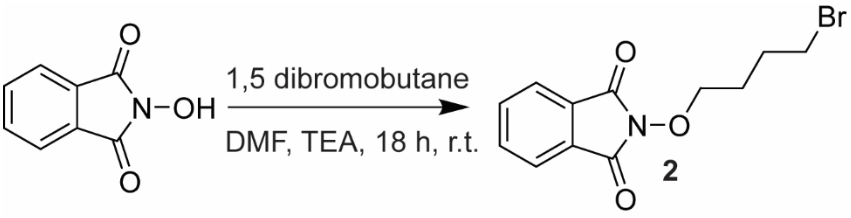


**2-(4-bromobutoxy)isoindoline-1,3-dione (2).** To a stirred solution of 1,4-dibromobutane (36.7 mmol, 3 eq.) and *N*-hydroxyphthalimide (12.2 mmol, 1 eq.) in DMF (20 mL) was added triethylamine (18.3 mmol, 1.5 eq.) at room temperature. After 16 hours the reaction was diluted in water and extracted with EtOAc three times. The combined organic phase was washed with brine and concentrated *in vacuo*. Flash column chromatography on silica using a gradient of 0-50% EtOAc/hexanes afforded **2** (2.71 g, 74.4%).

**$^1$H NMR (500 MHz, chloroform-*d*):** δ (ppm) 7.84 (m, 2H), 7.76 (m, 2H), 4.25 (t, *J* = 6.1 Hz, 2H), 3.55 (t, *J* = 6.5 Hz, 2H), 2.16 (m, 2H), 1.95 (m, 2H).

**$^{13}$C NMR (126 MHz, chloroform-*d*):** δ (ppm) 163.80, 134.71, 129.09, 123.74, 33.52, 28.98, 26.93, 1.49, 1.20.

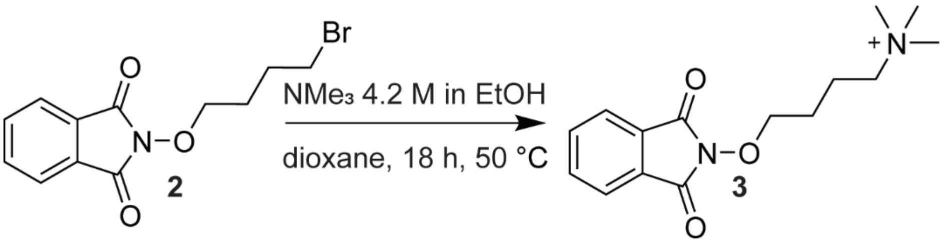


**4-((1,3-dioxoisoindolin-2-yl)oxy)-*N,N,N*-trimethylbutan-1-aminium bromide (3).** **2** (2.0 g, 6.7 mmol) was dissolved in dioxane (10 mL), and trimethylamine (35 wt.% in methanol) (3.0 mL, 17.7 mmol, 2.65 eq.) was added. The reaction was stirred overnight under reflux at 60 °C and concentrated *in vacuo*. Flash column chromatography on silica using a gradient of 0-100% MeOH/DCM afforded **3** (2.0 g, 5.6 mmol, 84%).

**$^1$H NMR (500 MHz, $D_2O$):** δ (ppm) 7.85 (m, 4H), 4.30 (t, *J* = 6.1 Hz, 2H), 3.44 (m, 2H), 3.18 (s, 9H), 2.10 (m, 2H), 1.84 (m, 2H).

**$^{13}$C NMR (126 MHz, $D_2O$):** δ (ppm) 165.25, 135.97, 130.29, 124.42, 78.24, 67.36, 53.65, 25.83, 20.78.

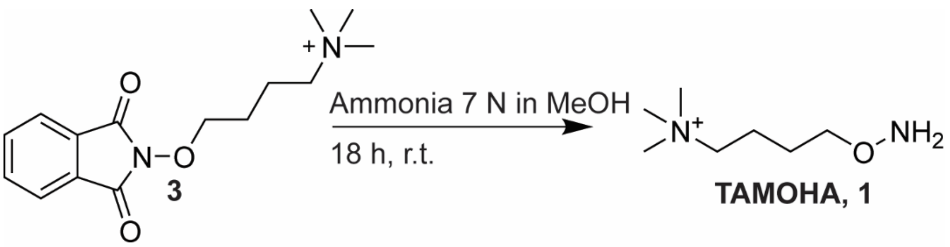


**4-(aminooxy)-*N,N,N*-trimethylbutan-1-aminium bromide (TAMOHA, 1).** This protocol was adapted from[5]. In this synthesis of TAMOHA (**1**), the use of ammonia and heat instead of the more commonly employed hydrazine for phthalimide cleavage was preferred, given difficulties encountered to ensure complete removal of excess hydrazine, which could compete with TAMOHA in trapping electrophiles. To a stirring solution of **3** (2.0 g, 5.6 mmol) in methanol (15 mL) was added a solution of ammonia in methanol (7 N $NH_3$ in MeOH, 15 mL, 105 mmol). The reaction vessel was capped with a rubber septum, and the reaction mixture was stirred until judged complete by TLC (~18 hours). The reaction mixture was then diluted with water (30 mL) and concentrated to dryness *in vacuo*. The resulting oil was redissolved in water (~30 mL), celite was added and the resulting mixture was concentrated to dryness *in vacuo* for dry loading. Reverse phase-flash chromatography (C18 silica gel, 0-5% ACN in $H_2O$) afforded **TAMOHA** (**1**) (1.12 g, 88%) as a white solid.

**$^1$H NMR (500 MHz, $D_2O$):** δ (ppm) 3.77 (t, 2H), 3.34 (m, 2H), 3.11 (s, 9H), 1.85 (m, 2H), 1.65 (m, 2H).

**$^{13}$C NMR (126 MHz, $D_2O$):** δ (ppm) 74.86, 66.35, 52.92, 24.41, 19.37.

**HRMS (ESI) *m/z*:** $[M]^+$ calcd. for $C_7H_{19}N_2O^+$ 147.1491, found 147.1493.

#### 2.2.2 Synthesis of 4-(aminooxy)-*N*,*N*,*N*-trimethylbutan-1-aminium-2,2,3,3-$D_4$ bromide (TAMOHA-$D_4$, 2)

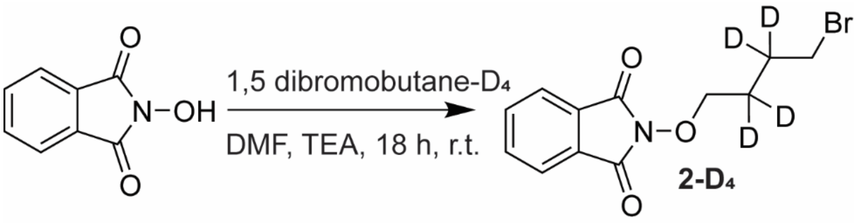


**2-(4-bromobutoxy-2,2,3,3-$d_4$)isoindoline-1,3-dione (2-$D_4$).** To a stirred solution of 1,4-dibromobutane-2,2,3,3-$d_4$ (4.63 mmol, 1.2 eq.) in DMF (5 mL) was added triethylamine (4.63 mmol, 1.2 eq.). Now, *N*-hydroxyphthalimide (3.85 mmol, 1 eq.) as a solution in DMF (2.6 mL) was added dropwise over 30 minutes at room temperature. After 16 hours the reaction was diluted in water and extracted with EtOAc three times. The combined organic phase was washed with brine and concentrated *in vacuo*. Flash column chromatography on silica using a gradient of 0-50% EtOAc/hexanes afforded **2-$D_4$** (647 mg, 2.14 mmol, 46.2%).

**$^1$H NMR (600 MHz, chloroform-*d*):** δ (ppm) 7.83 (m, 2H), 7.75 (m, 2H), 4.22 (s, 2H), 3.55 (s, 2H).

**$^{13}$C NMR (126 MHz, chloroform-*d*):** δ (ppm) 163.80, 134.72, 129.10, 123.75, 33.29, f28.11, 26.09, 1.21.

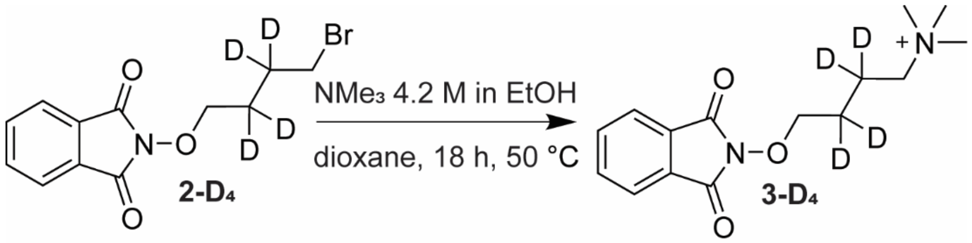


**4-((1,3-dioxoisoindolin-2-yl)oxy)-*N*,*N*,*N*-trimethylbutan-1-aminium-2,2,3,3-D4 bromide (3 $D_4$). 2-$D_4$** (2.132 mmol, 1 eq.) was dissolved in dioxane (2.12 mL) and trimethylamine (4.2M in methanol) (2.5 mL, 10.6 mmol, 5 eq.) was added. The reaction was stirred overnight under reflux at 60 °C and concentrated *in vacuo*. Flash column chromatography on silica using a gradient of 0-100% MeOH/DCM afforded **3-$D_4$** (592 mg, 2.105 mmol, 98.6%).

**$^1$H NMR (600 MHz, methanol-*$d_4$*):** δ (ppm) 7.86 (m, 4H), 4.27 (s, 2H), 3.49 (s, 2H), 3.2 (s, 9H).

**$^{13}$C NMR (126 MHz, methanol-*$d_4$*):** δ (ppm) 165.26, 135.98, 130.30, 124.43, 78.11, 67.22, 53.63.

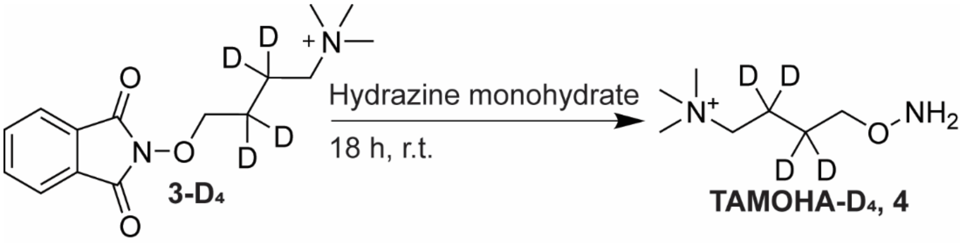


**4-(aminooxy)-*N,N,N*-trimethylbutan-1-aminium-2,3,3,3-D4 bromide (TAMOHA-$D_4$, 4). 3-$D_4$** (590 mg, 2.096 mmol) was dissolved in water (4.2 mL) and 590 µL of hydrazine monohydrate was added. The reaction was stirred overnight, acidified with conc. HCl, and filtered to remove precipitate. The filtrate was then basified with 10 M NaOH to near neutral pH, frozen, and lyophilized to dryness. Flash column chromatography on silica using a gradient of 0-100% MeOH/DCM afforded TAMOHA-$D_4$ (**4**, 200 mg, 1.322 mmol, 63%).

**$^1$H NMR (600 MHz, $D_2O$):** δ (ppm) 3.77 (s, 2H), 3.33 (s, 2H), 3.11 (s, 9H).

**$^{13}$C NMR (126 MHz, $D_2O$):** δ (ppm) 74.72, 66.20, 52.92.

**HRMS (ESI) *m/z*:** $[M]^+$ calcd. for $C_7H_{15}D_4N_2O^+$ 151.1743, found 151.1738.

### 2.3 TAMOHA quality assurance

**TAMOHA quality assurance.** TAMOHA condensation with ketones is immediate and irreversible. Therefore, care should be taken to not expose TAMOHA to traces of acetone. We recommend handling TAMOHA in flame-dried glassware and in a fume hood that does not have open containers of acetone. In addition, before each application of the TAMOHA probe, we recommend preparing two NMR samples, one that contains the TAMOHA probe alone, and one that contains the TAMOHA probe with 5 µL of acetone. The $^1$H NMR spectrum of the sample to which acetone was added should show complete conversion of TAMOHA to the corresponding oxime, as indicated by two methyl singlets at 1.891 and 1.877 ppm. These peaks should be absent in the NMR sample that contains only TAMOHA. This test ensures that the TAMOHA sample is pure and retains its reactivity (loss of reactivity can occur e.g., due to reduction of the nitrogen oxygen bond).

### 2.4 Synthesis of TAMOHA derivatives

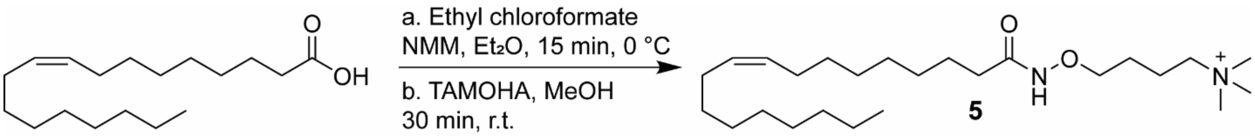


**Synthesis of the TAMOHA derivative of oleic acid (*N*,*N*,*N*-trimethyl-4-(oleamidooxy)butan-1-aminium, 5).** This protocol was adapted from[6]. To a stirred solution of oleic acid (0.248 mmol, 1 eq.) in diethyl ether (500 µL) at 0 °C was added ethyl chloroformate (0.223 mmol, 0.9 eq.) followed by 4-methylmorpholine (0.248 mmol, 1 eq.). An immediate white precipitate formed, and the reaction was stirred for 15 minutes. The reaction mixture was then passed through a pad of celite into a stirred solution of TAMOHA (0.273 mmol, 1.1 eq.) in methanol (500 µL). The reaction was stirred at room temperature for 30 minutes and concentrated *in vacuo.* Flash column chromatography on silica using a gradient of 0-40% MeOH/DCM afforded **5** (51 mg, 0.123 mmol, 50%).

**$^1$H NMR (600 MHz, methanol-*$d_4$*):** δ (ppm) 5.2 (m, 2H), 3.76 (t, 2H), 3.32 (m, 2H), 3.02 (s, 9H), 1.96 (t, 2H), 1.92 – 1.79 (m, 6H), 1.57 (M, *2*H), 1.46 (M, 2H), 1.27 – 1.09 (m, 20H), 0.76 (t, 3H).

**$^{13}$C NMR (126 MHz, methanol-*$d_4$*):** δ 172.94, 130.88, 130.75, 76.17, 67.37, 53.55, 33.82, 33.05, 30.83, 30.81, 30.61, 30.45, 30.34, 30.30, 30.21, 30.17, 28.13, 26.64, 25.69, 23.74, 20.94, 14.50.

**HRMS (ESI) *m/z*:** $[M]^+$ calcd. for $C_7H_{15}D_4N_2O^+$ 151.1743, found 151.1738.

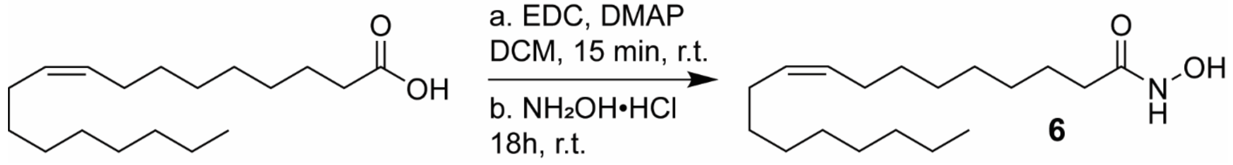

**Synthesis of *N*-hydroxyoleamide (6).** To a stirred solution of oleic acid (0.0885 mmol, 1 eq.) in DCM (885 µL) was added EDC (0.0885 mmol, 1 eq.) and DMAP (0.2046 mmol, 2.31 eq.). The reaction was left to stir for 15 minutes at room temperature and hydroxylamine hydrochloride (0.0885 mmol, 1 eq.) was added. The reaction was left to stir at room temperature for 18 hours and concentrated *in vacuo.* Flash column chromatography on silica using a gradient of 0-30% EtOAc/hexanes afforded **6** (12 mg, 0.0403 mmol, 45.58%)

**$^{1}$H NMR (500 MHz, methanol-*$d_4$*)** δ 5.35 (s, 2H), 2.31 (t, 2H), 2.03 (q, 4H), 1.60 (m, 2H), 1.40 – 1.25 (m, 20H), 0.90 (t, 3H).

**$^{13}$C NMR** (**126 MHz, methanol-*$d_4$***) δ 176.00, 130.91, 130.78, 51.95, 34.80, 33.07, 30.84, 30.76, 30.61, 30.46, 30.34, 30.25, 30.18, 30.14, 28.12, 28.09, 26.02, 23.74, 14.44.

**HRMS (ESI) *m/z*:** $[M+H]^+$ calcd. for $C_{18}H_{36}NO_2^+$ 298.2740, found 298.2733.

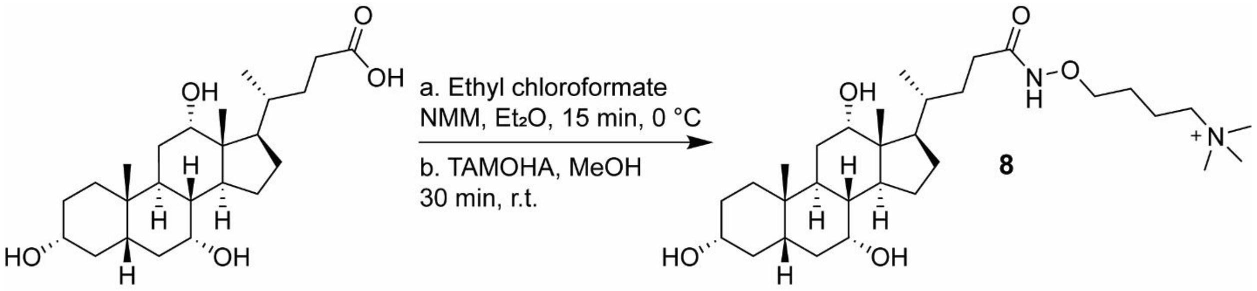

**Synthesis of the TAMOHA derivative of cholic acid *(N,N,N*-trimethyl-4-(((*R*)-4-((3*R*,5*S*,7*R*,8*R*,9*S*,10*S*,12*S*,13*R*,14*S*,17*R*)-3,7,12-trihydroxy-10,13-dimethylhexadecahydro-1H-cyclopenta[a]phenanthren-17-yl)pentanamido)oxy)butan-1-aminium, 8).** This protocol was adapted from[6]. To a stirred solution of cholic acid (0.085 mmol, 1 eq.) in diethyl ether (500 µL) with minimal DMF for solubility at 0 °C was added ethyl chloroformate (0.34 mmol. 4 eq.) followed by 4-methylmorpholine (0.34 mmol, 4 eq.). An immediate white precipitate formed, and the reaction was stirred for 15 minutes. The reaction mixture was then passed through a pad of celite into a stirred solution of TAMOHA (0.34 mmol, 4 eq.) in methanol (500 µL). The reaction was stirred at room temperature for 30 minutes and concentrated *in vacuo.* Flash column chromatography on silica using a gradient of 0-40% MeOH/DCM afforded **8** (26 mg, 0.048 mmol, 56.8%)

**$^{1}$H NMR (500 MHz, methanol-*$d_4$*):** δ (ppm) 3.95 (t, 1H), 3.90 (t, 2H), 3.80 (m, 1H), 3.45 (m, 2H), 3.38 (m, 1H), 3.16 (s, 9H), 2.33 – 2.22 (m, 2H), 2.17 (m, 1H), 2.07 – 1.93 (m, 5H), 1.93-1.85 (m, 2H), 1.85 – 1.77 (m, 2H), 1.77 – 1.68 (m, 3H), 1.68 – 1.63 (m, 1H), 1.63 – 1.50 (m, 5H), 1.47-1.33 (m, 4H), 1.33 – 1.23 (m, 1H), 1.12 (m, 1H), 1.03 (d, 3H), 1.02 – 0.93 (m, 1H), 0.92 (s, 3H), 0.71 (s, 3H).

**$^{13}$C NMR** (**126 MHz, methanol-*$d_4$***)**:** δ (ppm) 173.59, 76.15, 73.96, 72.83, 68.98, 67.42, 67.40, 67.38, 53.60, 53.57, 53.54, 47.87, 47.46, 43.16, 43.03, 40.98, 40.45, 36.85, 36.46, 35.89, 35.88, 32.94, 31.17, 30.78, 29.60, 28.71, 27.88, 25.71, 24.20, 23.16, 20.97, 17.70, 12.97.

**HRMS (ESI) *m/z*:** $[M]^+$ calcd. for $C_{31}H_{57}N_2O_5^+$ 537.4262, found 537.4258.

### 3. Supporting Figures

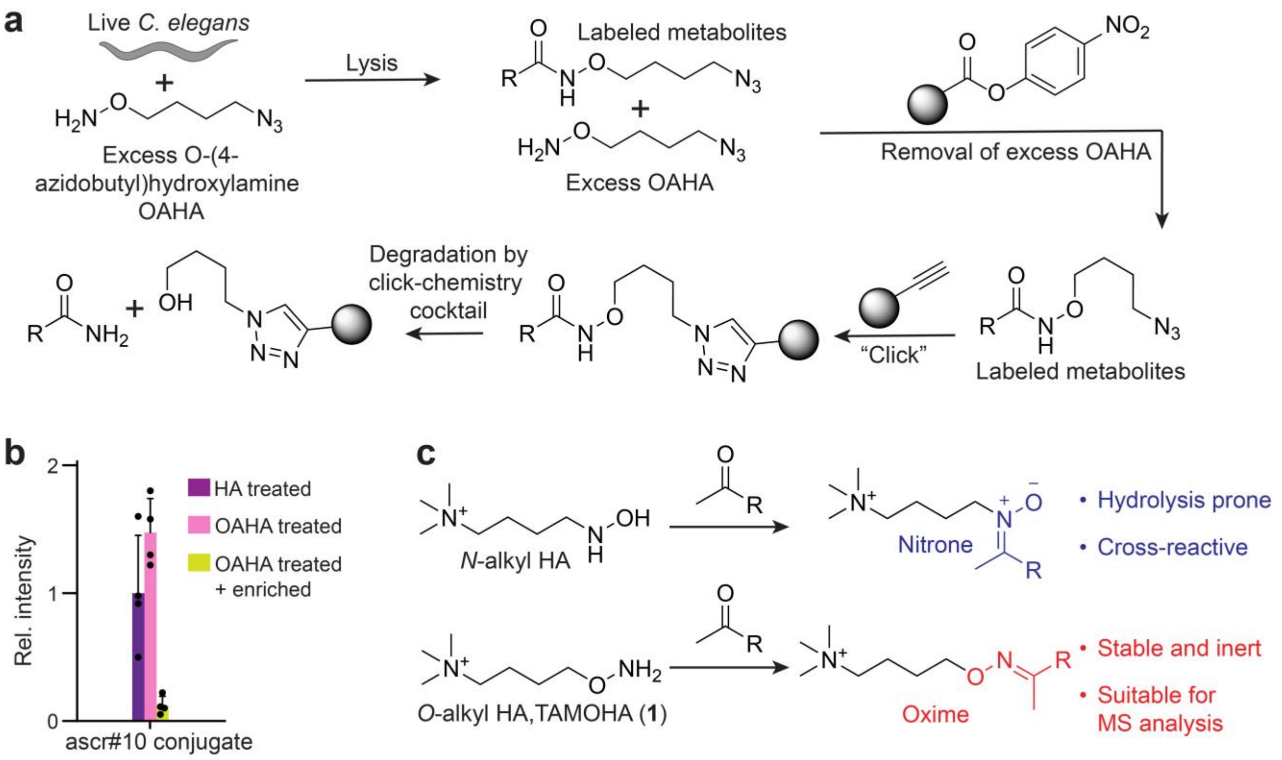


**Figure S1.** Initial approaches to capture and enhance detection of the activated metabolome. (a) Scheme of OAHA labeling strategy and degradation by the click chemistry cocktail. (b) Relative intensity of HA-labeled (purple), OAHA labeled (pink), and OAHA labeled ascr#10 followed by click chemistry mediated enrichment (green). (c) Scheme of *N*-alkyl and *O*-alkyl hydroxylamine derivative reactivity and product stability.

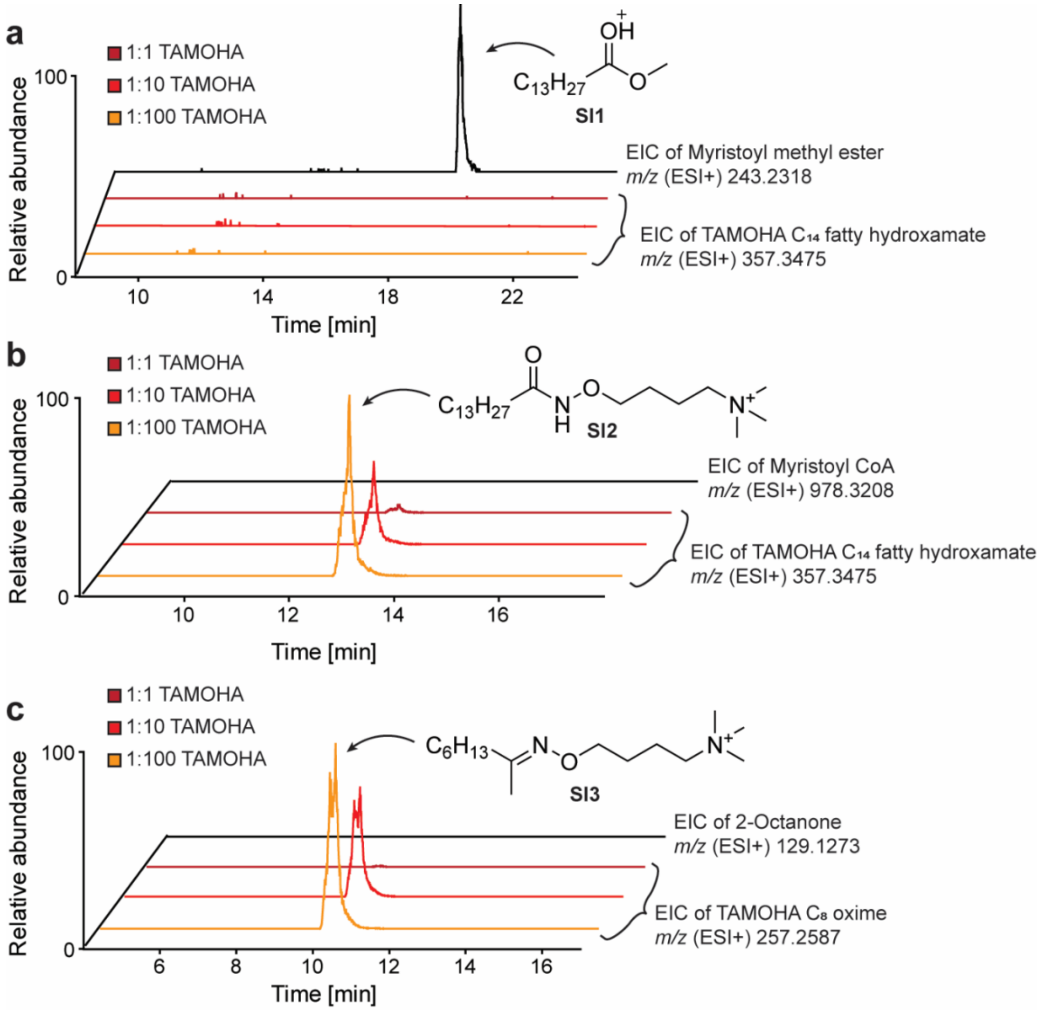


**Figure S2.** TAMOHA model reactions. (a) Ion chromatograms of myristoyl methyl ester (**SI1**) (top) or $C_{14}$ TAMOHA fatty hydroxamate (**SI2**) in samples of myristoyl methyl ester incubated with a 1:1 (maroon), 1:10 (red), or 1:100 (orange) molar ratio of TAMOHA. (b) Ion chromatograms of myristoyl coenzyme A ester in methanol (top) or $C_{14}$ TAMOHA fatty hydroxamate (**SI2**) in samples of myristoyl coenzyme A ester incubated with a 1:1 (maroon), 1:10 (red), or 1:100 (orange) molar ratio of TAMOHA. (c) Ion chromatograms of 2-octanone in methanol (top) or $C_8$ TAMOHA ketone oxime (**SI3**) in samples of 2-octanone incubated with a 1:1 (maroon), 1:10 (red), or 1:100 (orange) molar ratio of TAMOHA.

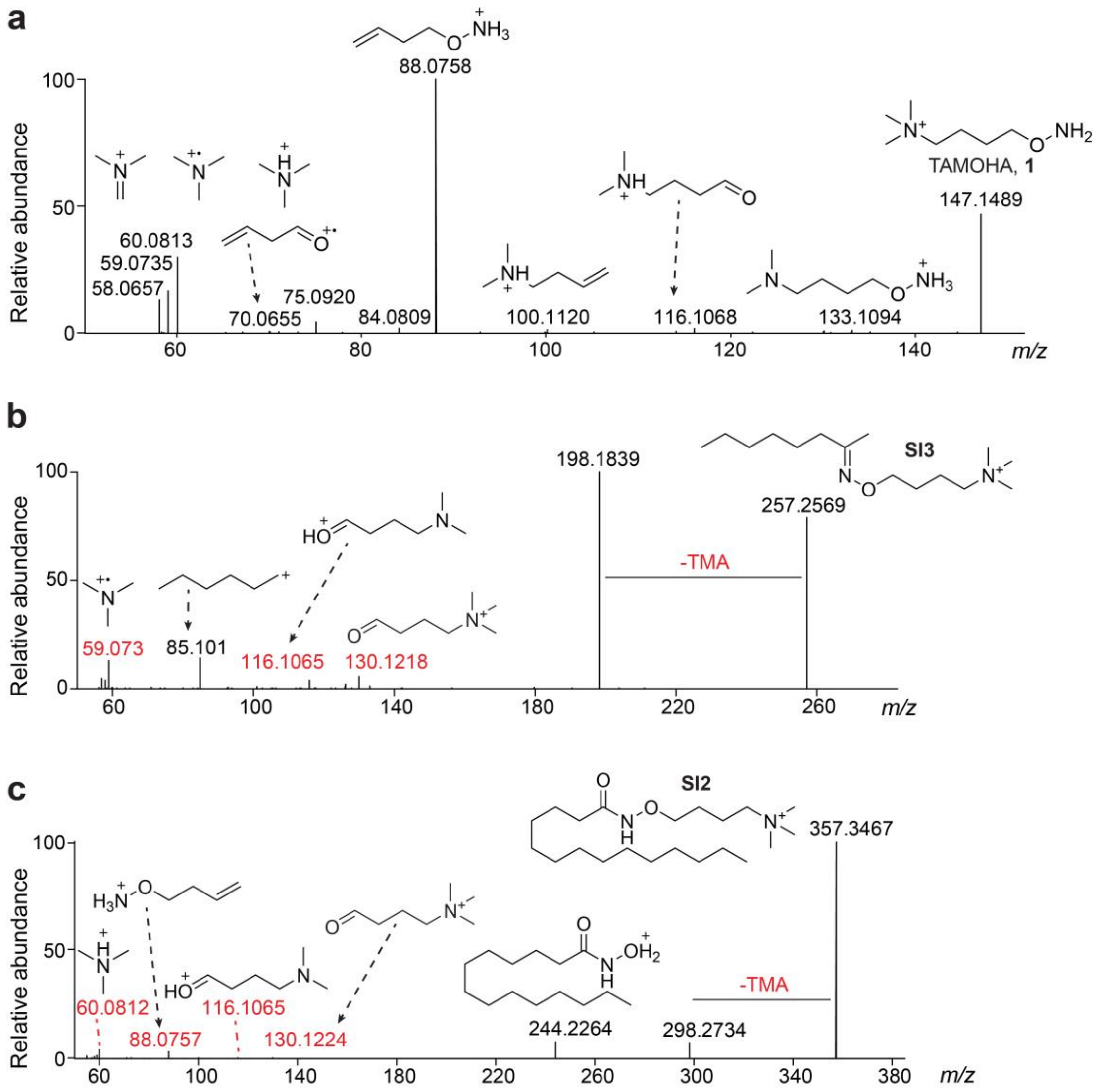


**Figure S3.** Fragmentation of TAMOHA and TAMOHA labeled metabolites. (a) Annotated $MS^2$ Spectrum of TAMOHA (**1**). (b) Annotated $MS^2$ spectrum of $C_8$ TAMOHA ketone oxime (**SI3**) from TAMOHA model studies (above). (c) Annotated $MS^2$ spectrum of $C_{14}$ TAMOHA fatty hydroxamate (**SI2**) from TAMOHA model studies (above).

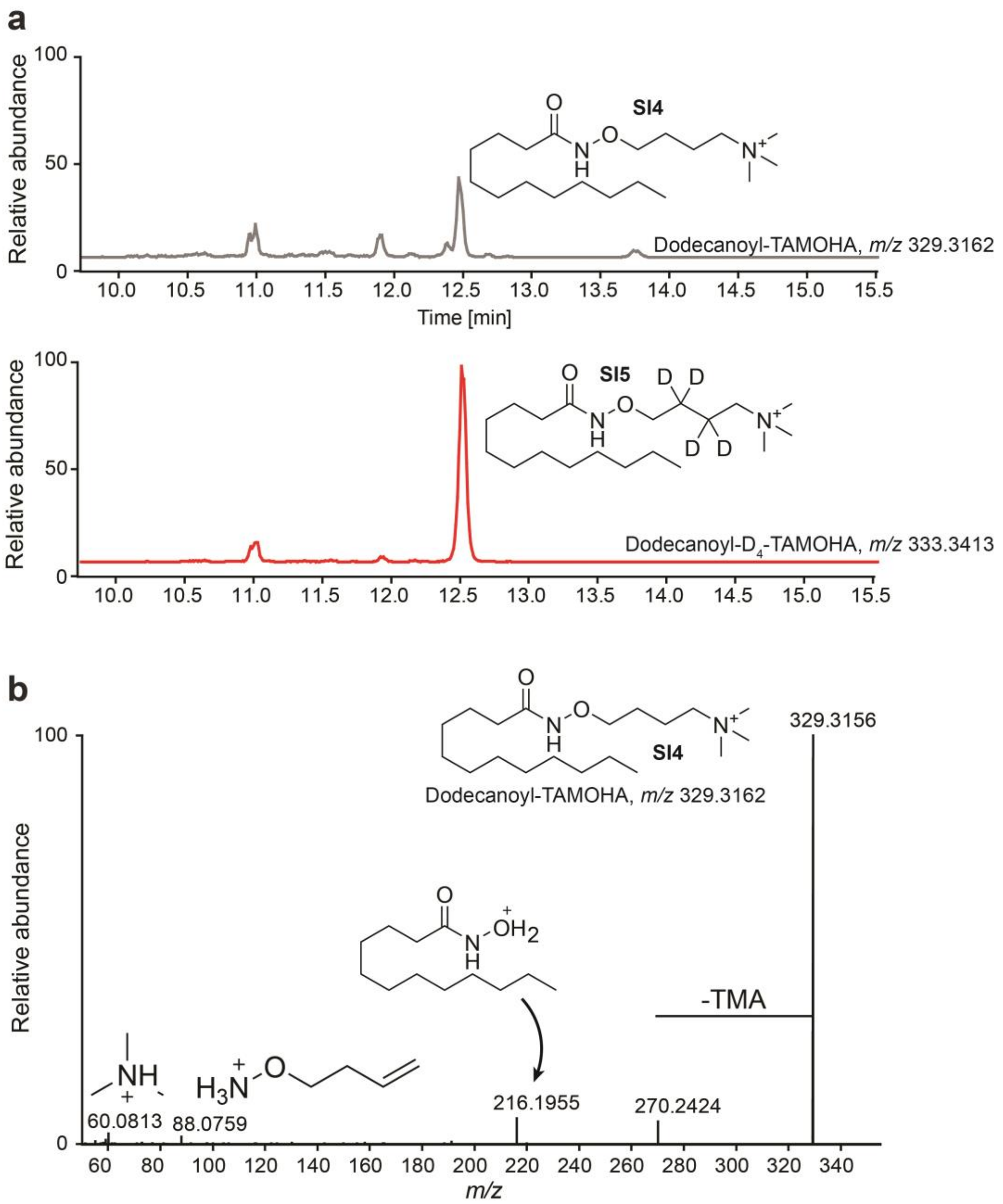


**Figure S4.** Use of $D_4$-TAMOHA to confirm TAMOHA-labeled mass features. (a) Ion chromatograms of dodecanoyl-TAMOHA (**SI4**) (top) and dodecanoyl-$D_4$-TAMOHA (**SI5**) (bottom). (b) Annotated $MS^2$ spectrum of dodecanoyl-TAMOHA (**SI4**).

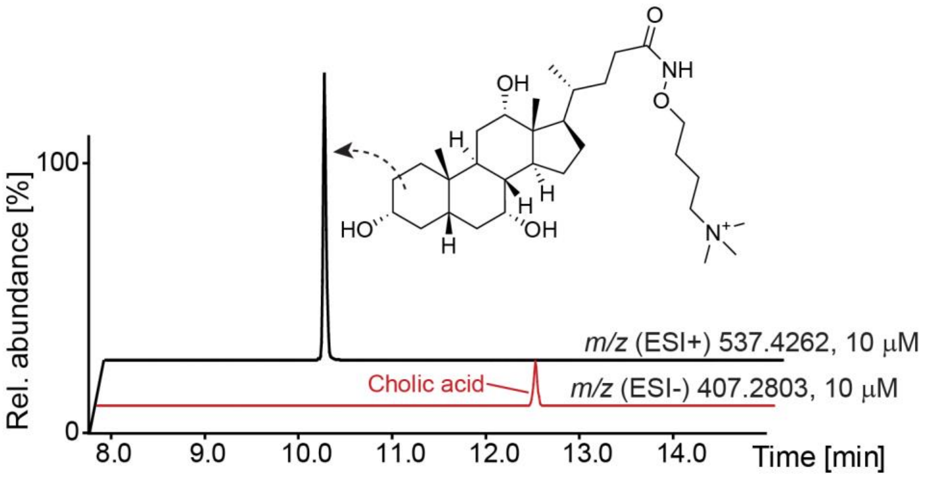


**Figure S5.** TAMOHA ionization enhancement. (a) Ion chromatograms of synthetic standards of the cholic acid TAMOHA conjugate (top, ESI+), and cholic acid (bottom, ESI-).

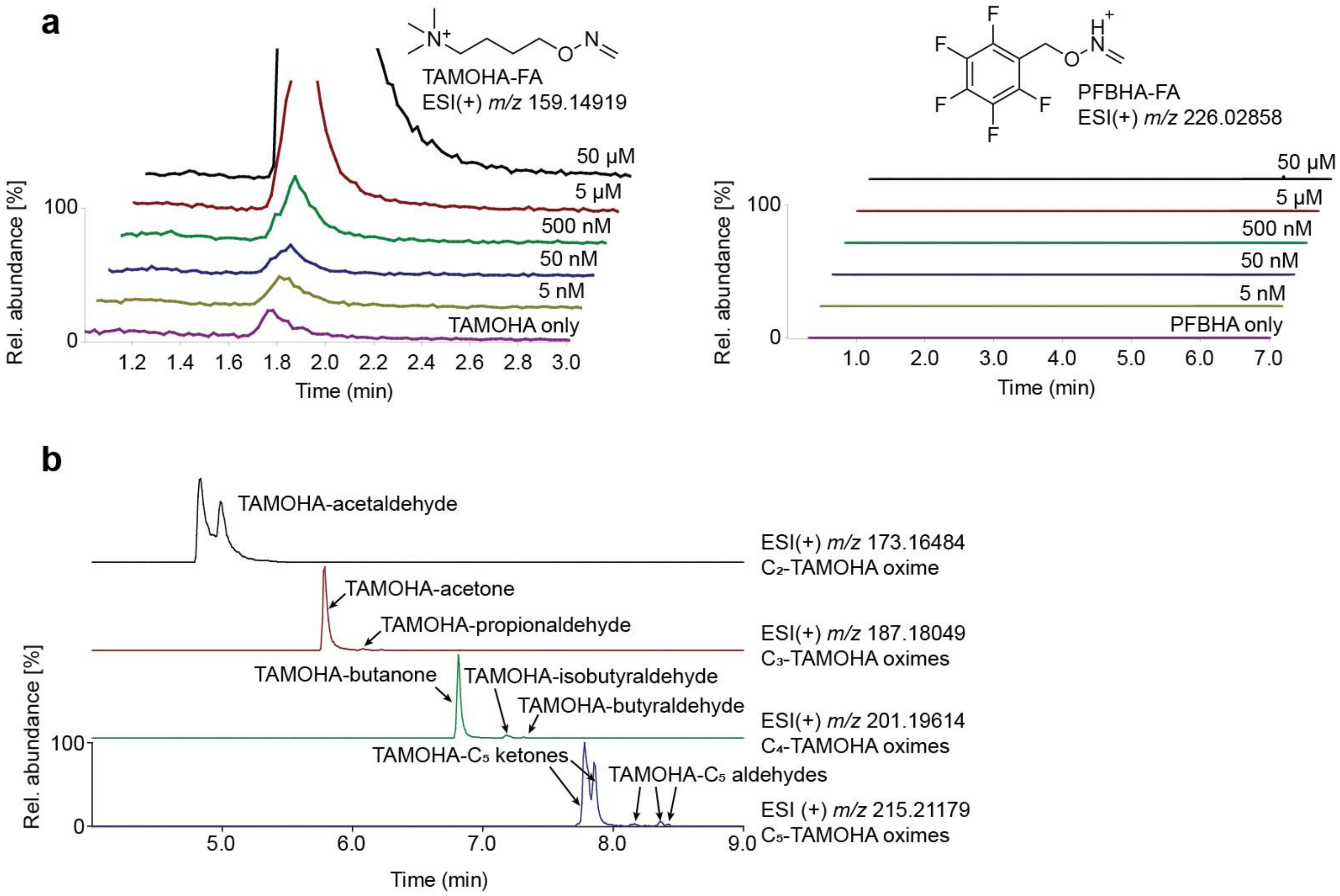


**Figure S6.** Detection of small aliphatic aldehydes. (a) Ion chromatograms of TAMOHA or PFBHA (0.05 M) reacted with a dilution series of formaldehyde ranging from 50 µM (top) to 5 nM (bottom). (b) Annotated extracted ion chromatograms of TAMOHA labeled short chain ketones and aldehydes present in mouse liver.

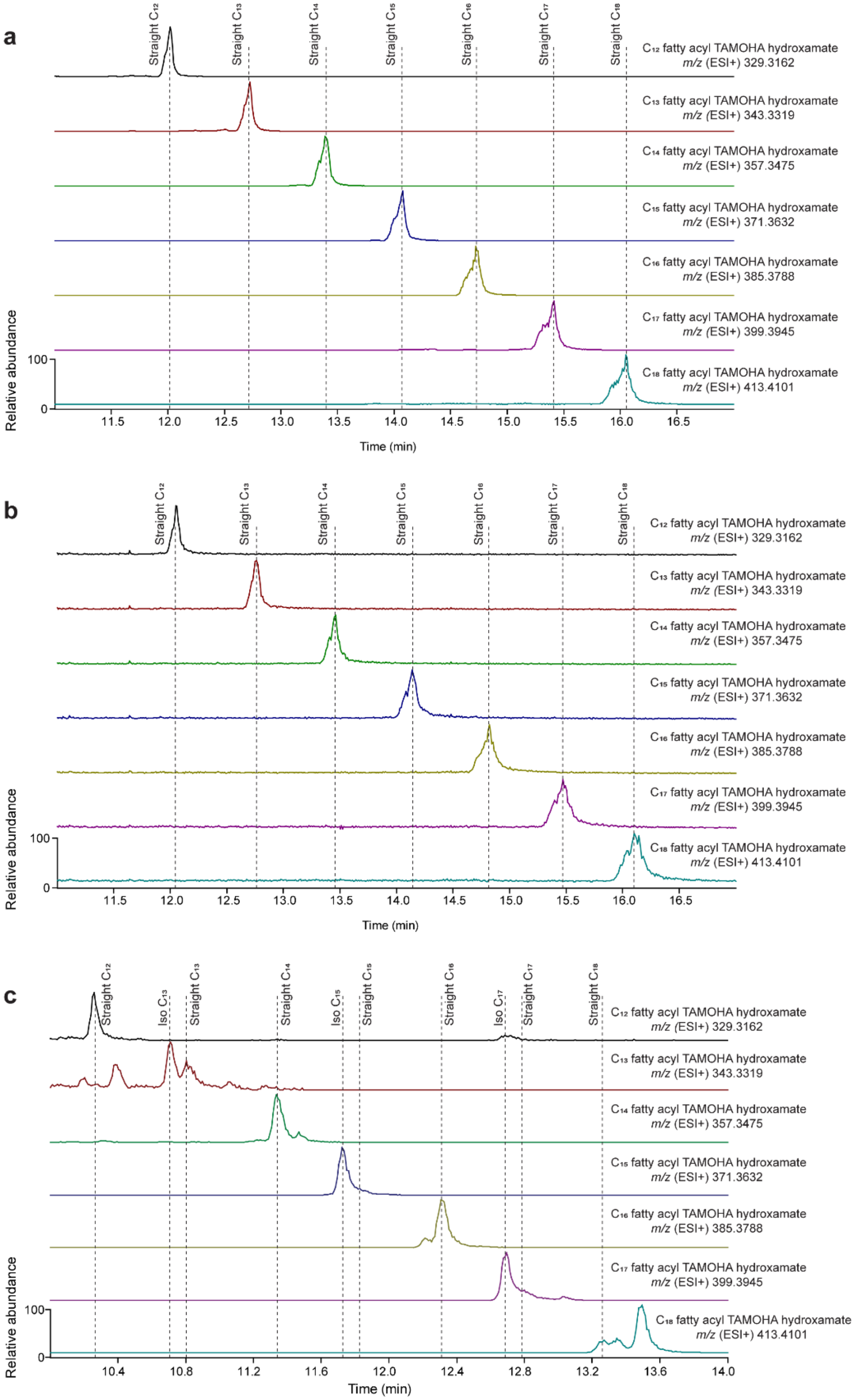


**Figure S7.** Homologous series of TAMOHA fatty hydroxamates. (a) Ion chromatograms of synthetic standards of $C_{12}$ (top) though $C_{18}$ (bottom) saturated, straight chain TAMOHA fatty acid hydroxamates. (b) and (c) show ion chromatograms of $C_{12}$ though $C_{18}$ saturated TAMOHA fatty acid hydroxamates in *E. coli* and in *C. elegans*, respectively. Samples in (c) were run using a different column which results in different retention times.

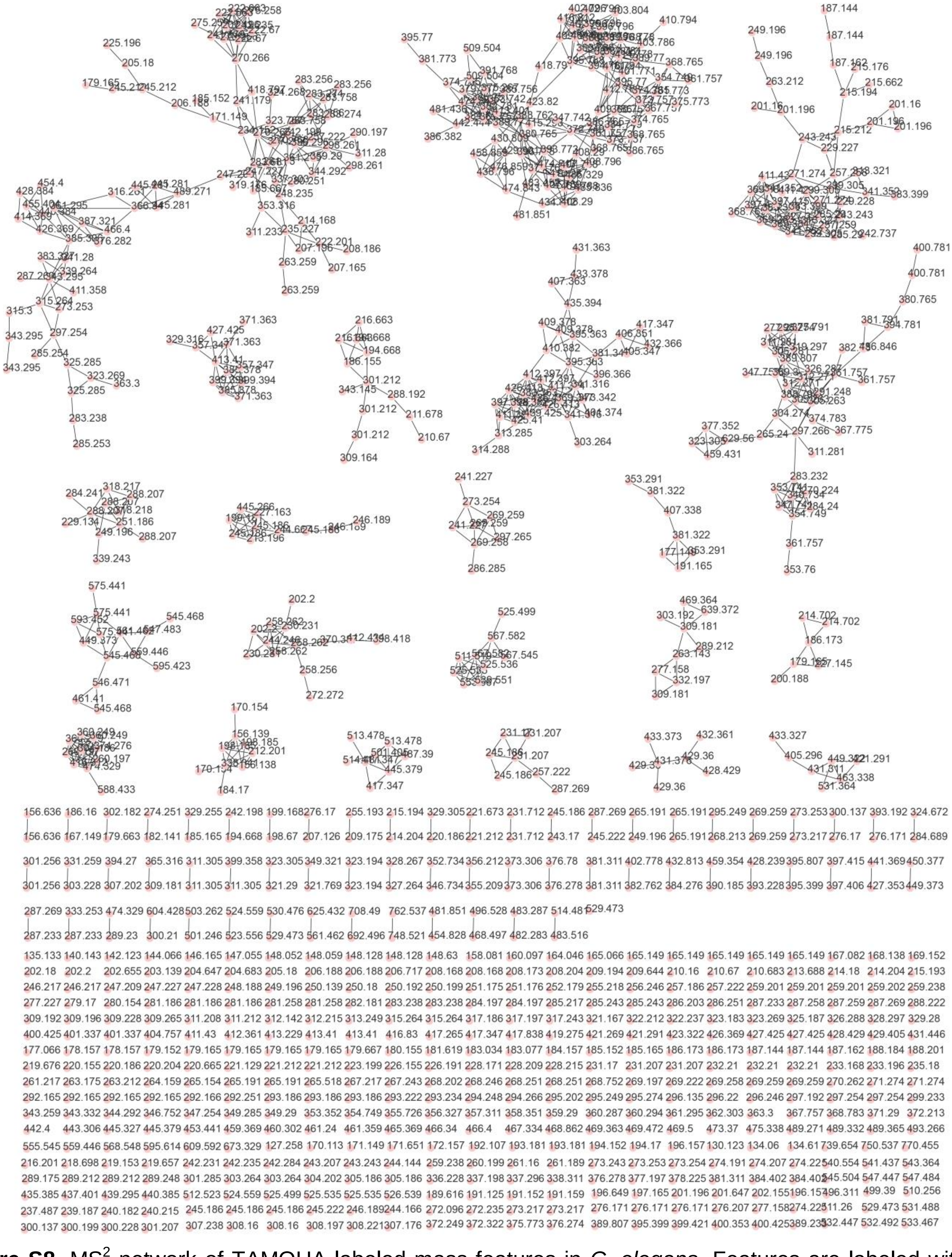

**Figure S8.** MS$^2$ network of TAMOHA labeled mass features in *C. elegans*. Features are labeled with *m/z*.

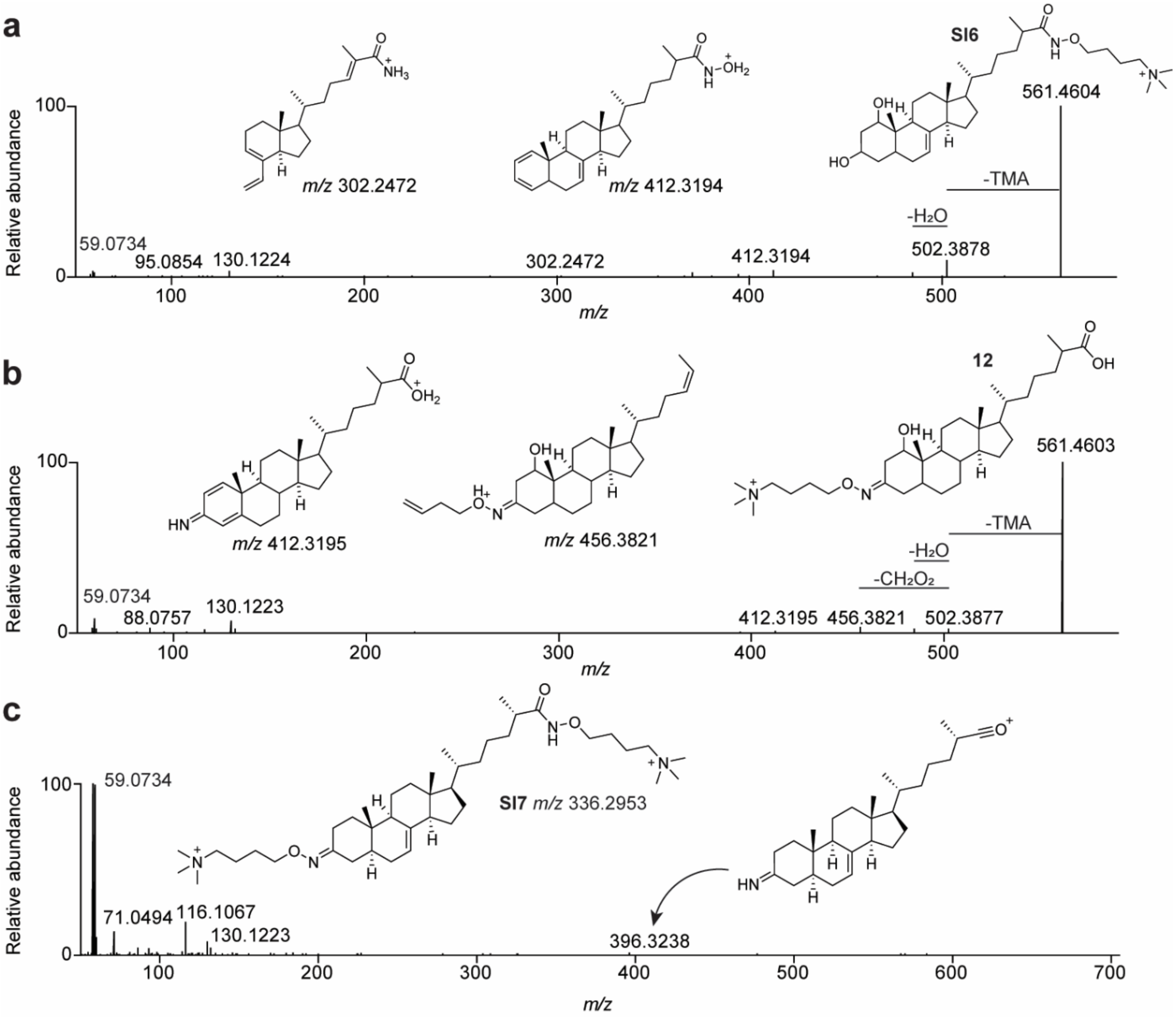


**Figure S9.** Fragmentation of TAMOHA labeled dafachronic acid derivatives. (a) Annotated $MS^2$ spectrum of **SI6** and (b) its isomer, **12**. (c) Annotated $MS^2$ spectrum of **SI7.**

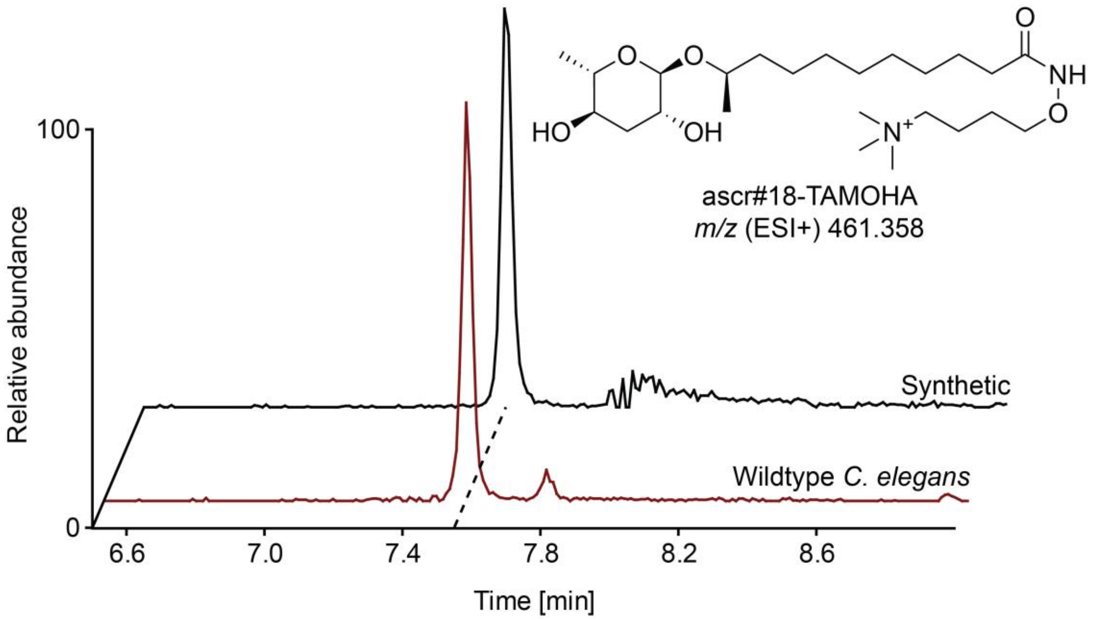


**Figure S10.** Structure confirmation of ascr#18-TAMOHA. (a) Ion chromatograms of a synthetic standard of ascr#18-TAMOHA (black) and the mass feature corresponding to ascr#18-TAMOHA in wildtype *C. elegans* (maroon).

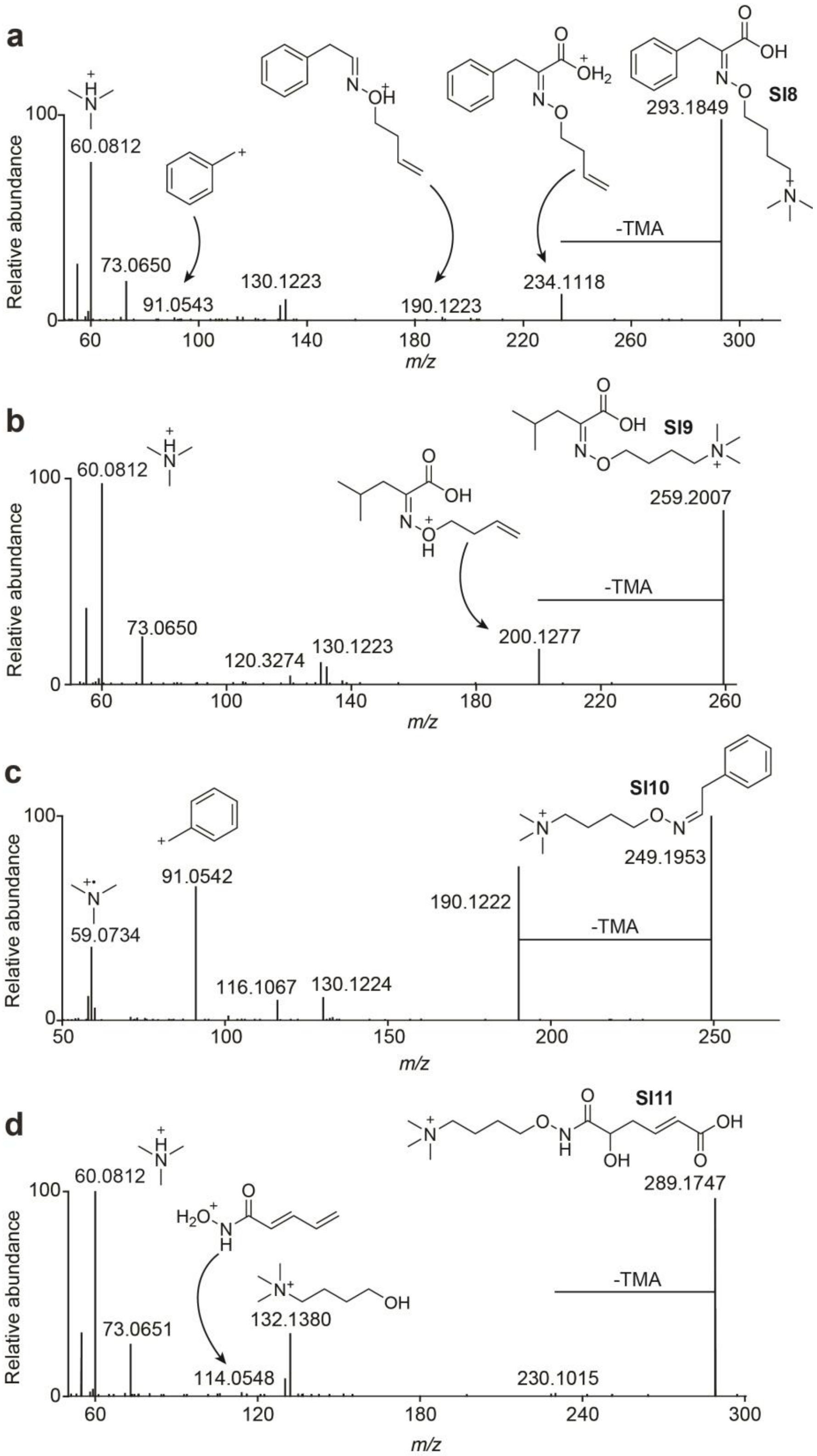


**Figure S11.** Fragmentation of TAMOHA labeled metabolites. Annotated $MS^2$ spectra of **SI8** (a) **SI9** (b), **SI10** (c), and **SI11** (d).

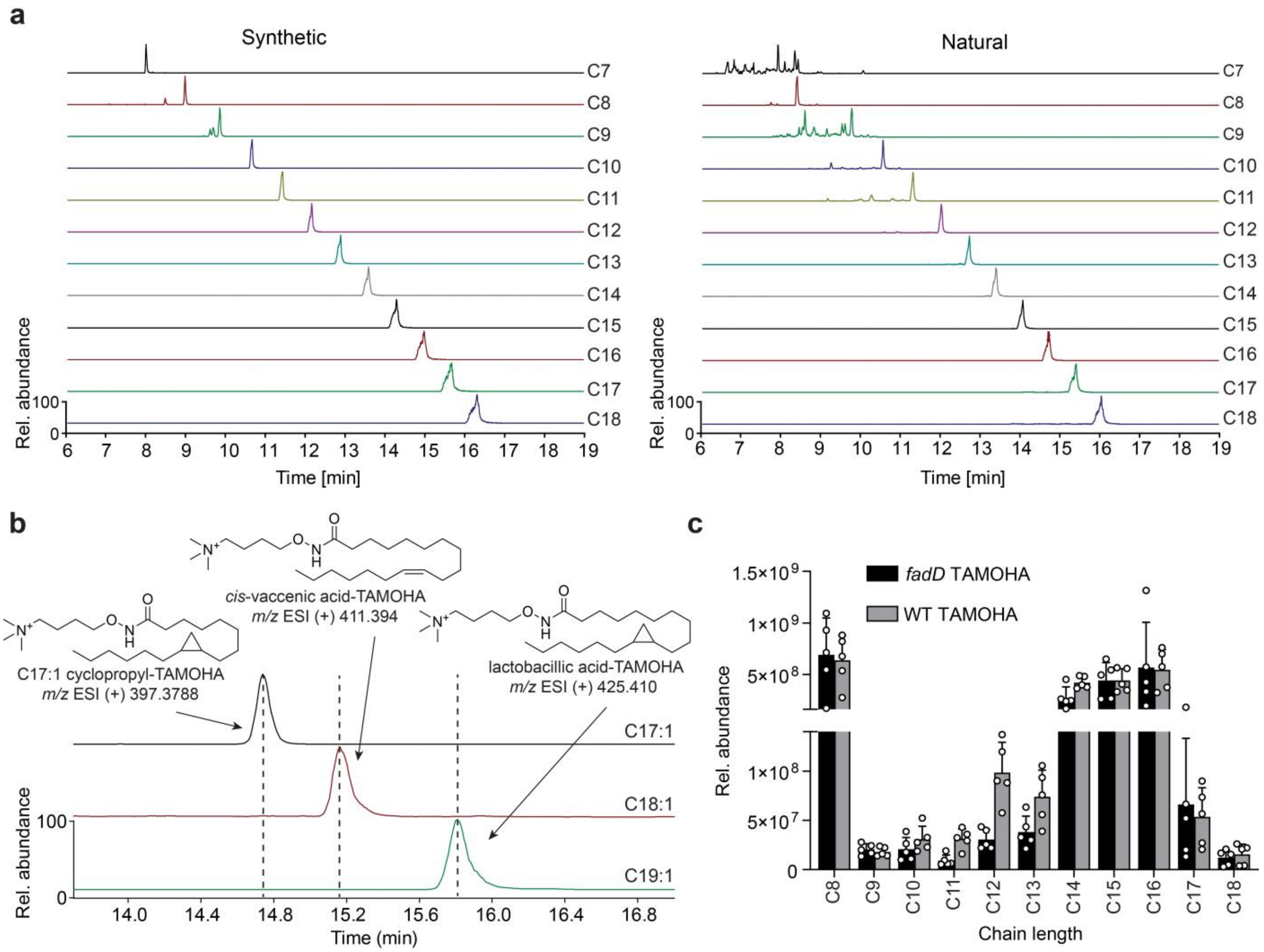


**Figure S12.** TAMOHA labeled fatty acid profiles in *E. coli*. (a) Grid of saturated $C_7$ through $C_{18}$ TAMOHA labeled fatty acid synthetic standards (left) and in TAMOHA treated *E. coli* (right). (b) Ion chromatograms of C17:1 cyclopropane containing TAMOHA fatty hydroxamate (top) *cis*-vaccenic acid TAMOHA fatty hydroxamate (middle), and lactobacillic acid TAMOHA fatty hydroxamate (bottom). (c) Profiles of saturated $C_8$ through $C_{18}$ TAMOHA labeled fatty acids in wildtype (gray) and *fadD* mutant (black) *E. coli*.

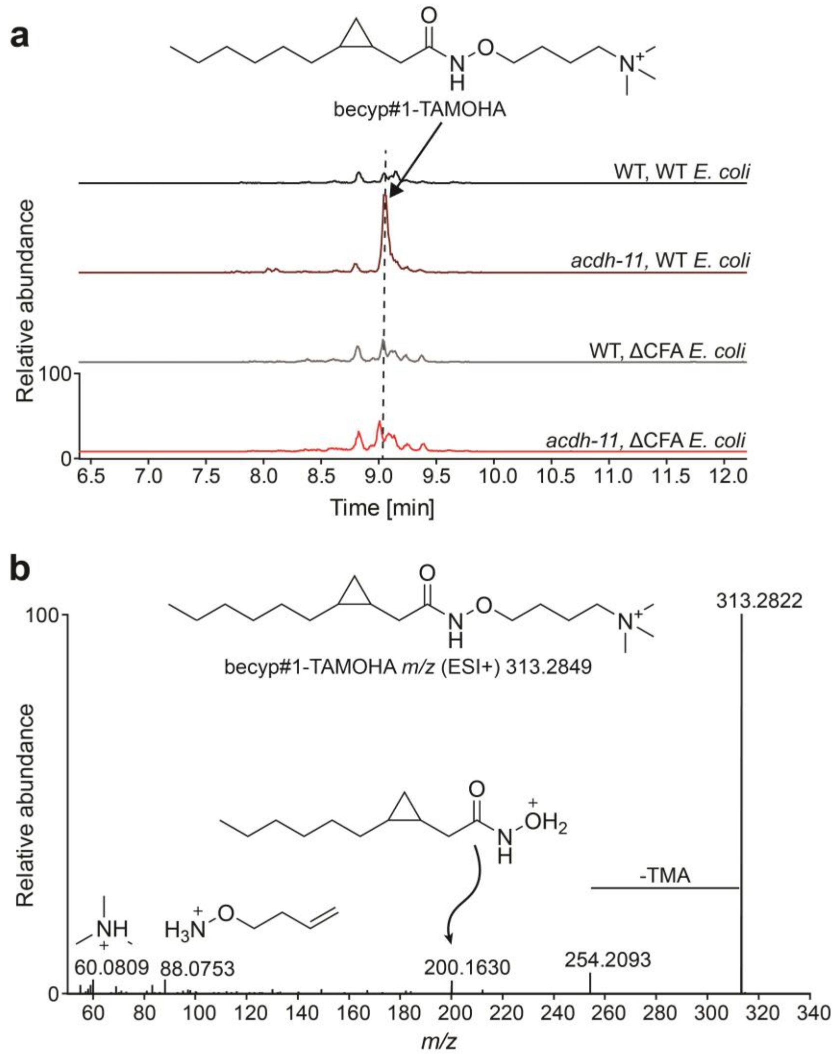


**Figure S13.** TAMOHA labeled becyp#1. (a) Ion chromatograms of becyp#1-TAMOHA in wildtype or *acdh-11* mutant *C. elegans* fed wildtype or ΔCFA *E. coli*. (b) Annotated $MS^2$ spectrum of becyp#1-TAMOHA.

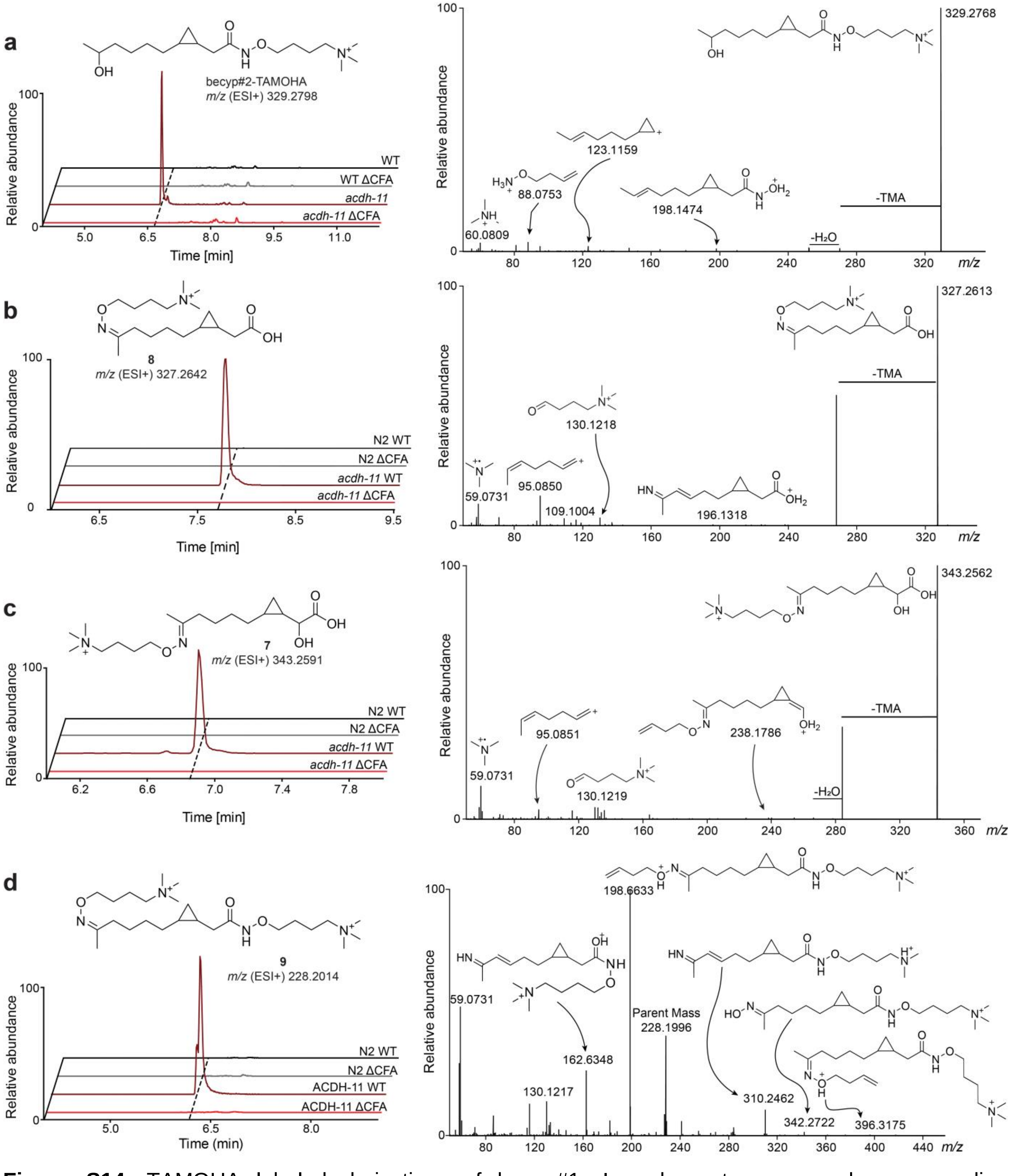


**Figure S14.** TAMOHA labeled derivatives of becyp#1. Ion chromatograms and corresponding annotated $MS^2$ fragmentation spectra of becyp#2-TAMOHA (a), **8** (b), **7** (c), and **9** (d) in wildtype or *acdh-11* mutant *C. elegans* fed wildtype or ΔCFA *E. coli*.

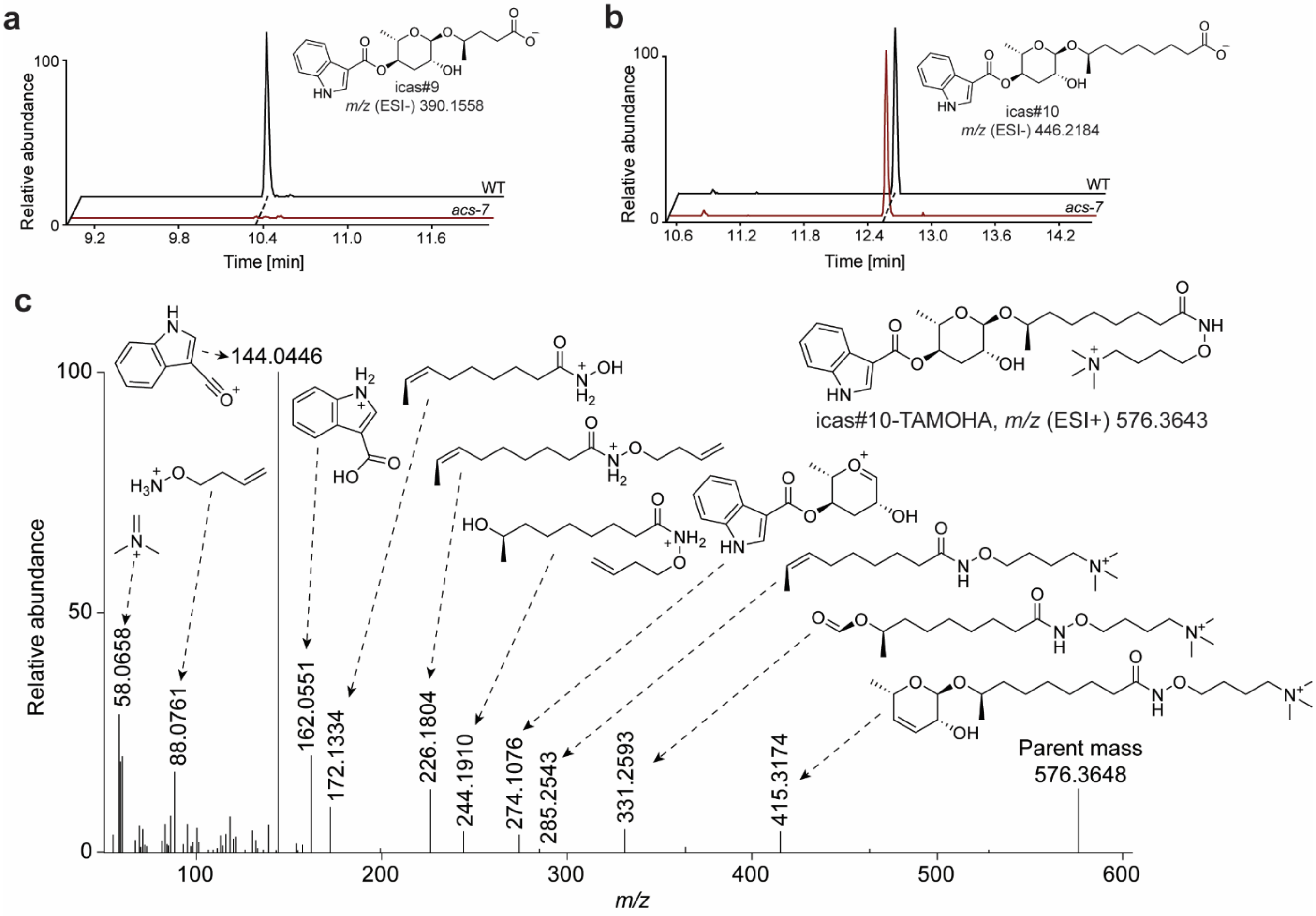


**Figure S15.** TAMOHA labeling in *acs-7* mutant *C. elegans*. Ion chromatograms of icas#9 (a) and icas#10 (b) in wildtype (black) and *acs-7* mutant (red) *C. elegans*. (c) Annotated $MS^2$ spectrum of icas#10-TAMOHA.

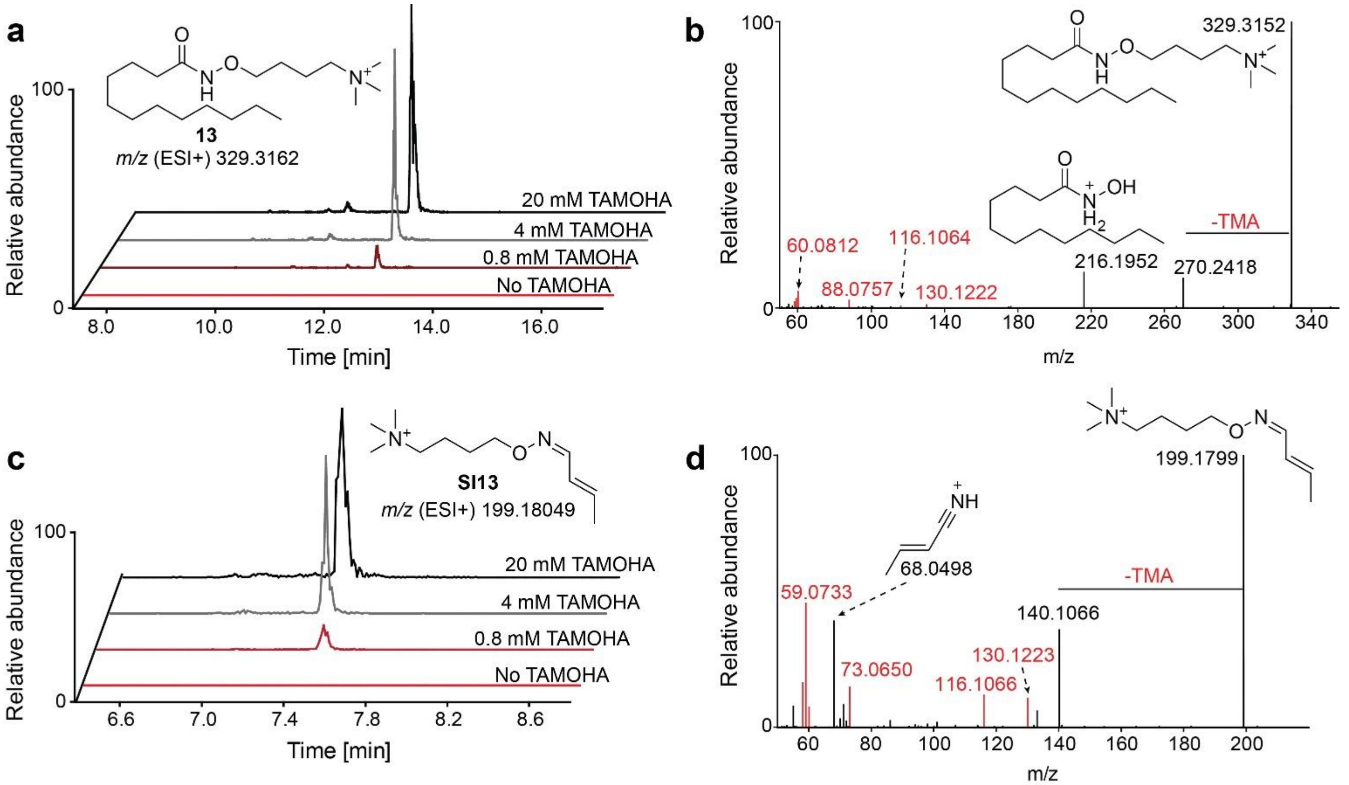


**Figure S16.** Optimization of TAMOHA treatment concentration. (a) Ion chromatograms of $C_{12}$ TAMOHA fatty hydroxamate (**12**) in *E. coli* treated with 20 mM (black), 4 mM (gray), 0.8 mM (maroon), or PBS only control (red). (b) Annotated $MS^2$ spectrum of $C_{12}$ TAMOHA fatty hydroxamate (**12**). (c) Ion chromatograms of the TAMOHA conjugate of crotonaldehyde (**SI13**) in *E. coli* treated with 20 mM (black), 4 mM (gray), 0.8 mM (maroon), or PBS only control (red). (d) Annotated $MS^2$ spectrum of the TAMOHA conjugate of crotonaldehyde (**SI13**). Characteristic TAMOHA fragments are shown in red.

### 4. NMR spectra appendix

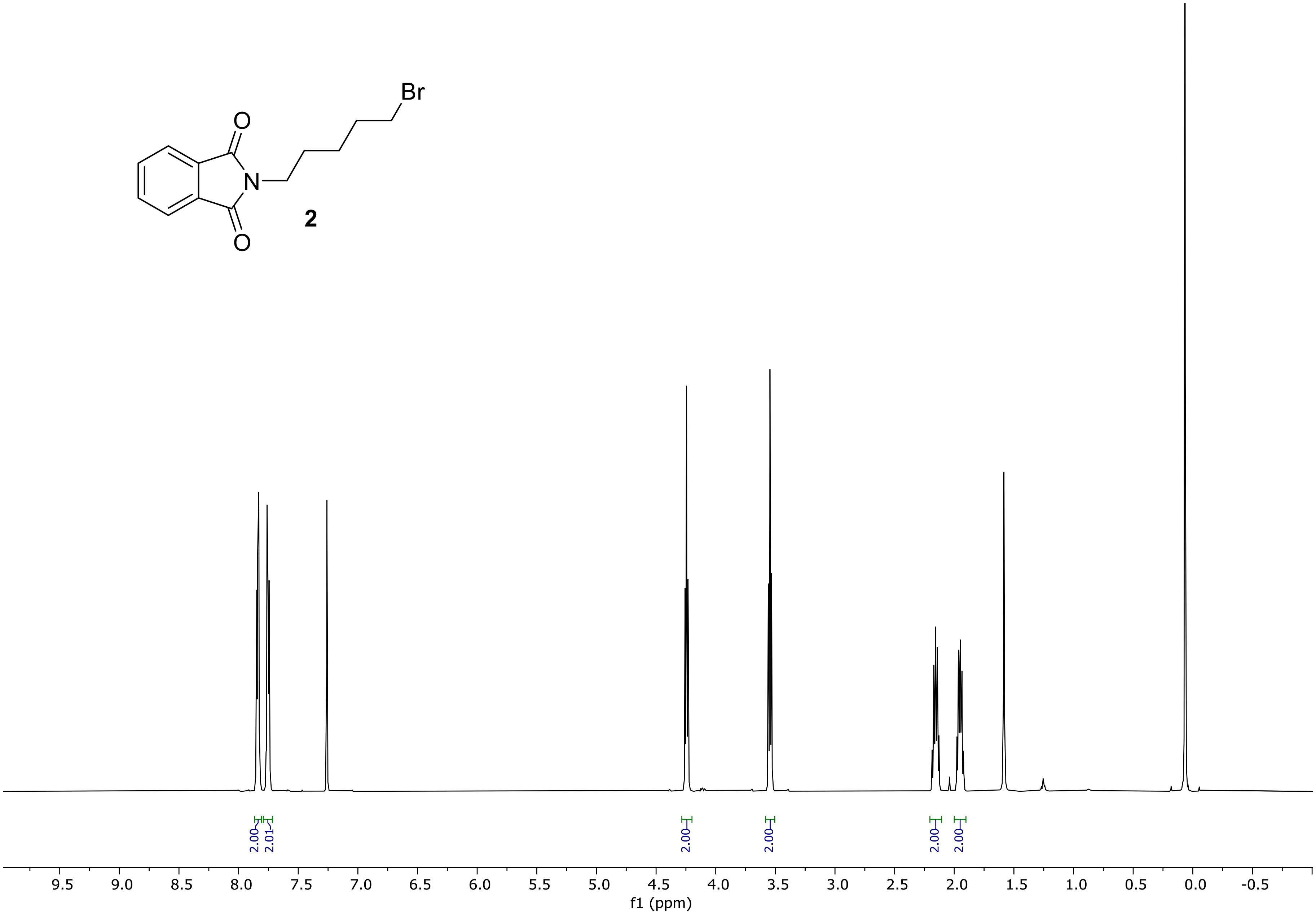


$^1$H NMR spectrum (500 MHz) of 2-(4-bromobutoxy)isoindoline-1,3-dione (**2**) in chloroform-*d*.

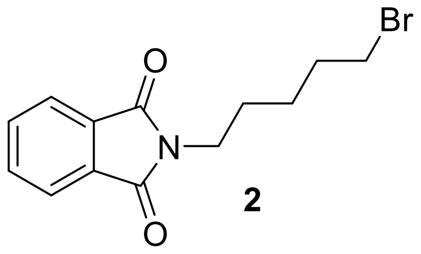

**2**

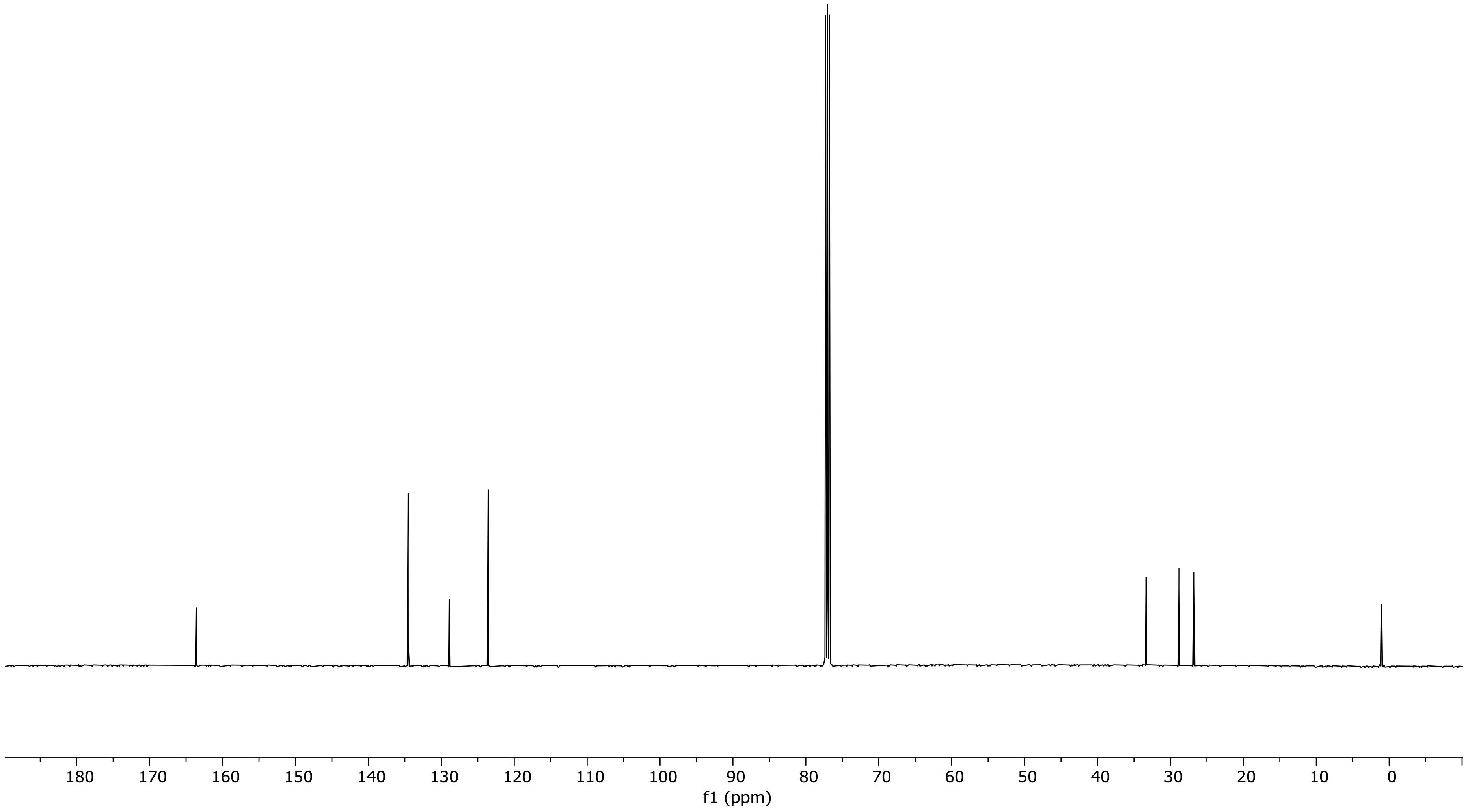


$^{13}$C NMR spectrum (126 MHz) of 2-(4-bromobutoxy)isoindoline-1,3-dione (**2**) in chloroform-*d*.

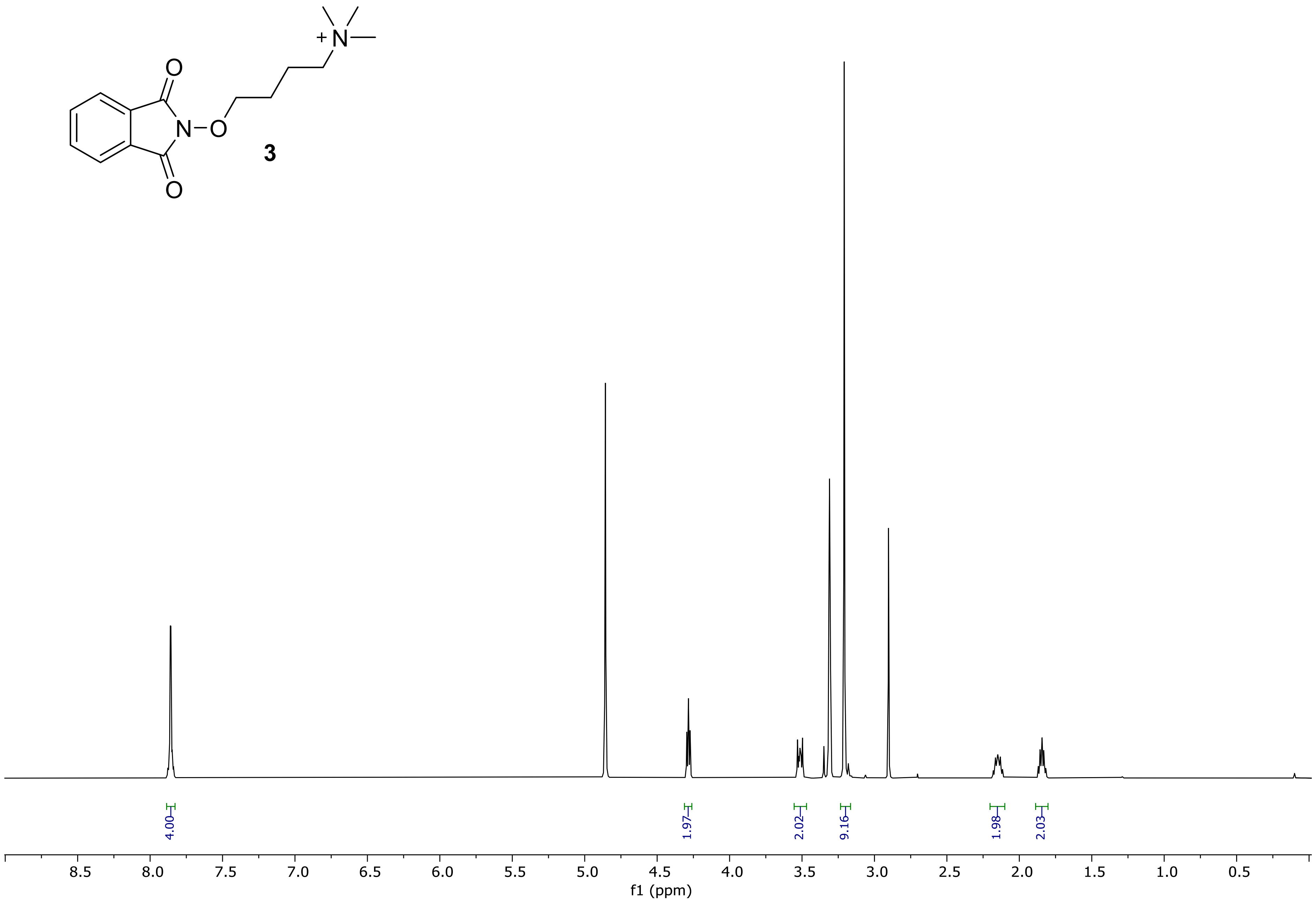


[1]H NMR spectrum (500 MHz) of 4-((1,3-dioxoisoindolin-2-yl)oxy)-*N,N,N*-trimethylbutan-1-aminium bromide (**3**) in methanol-$d_4$.

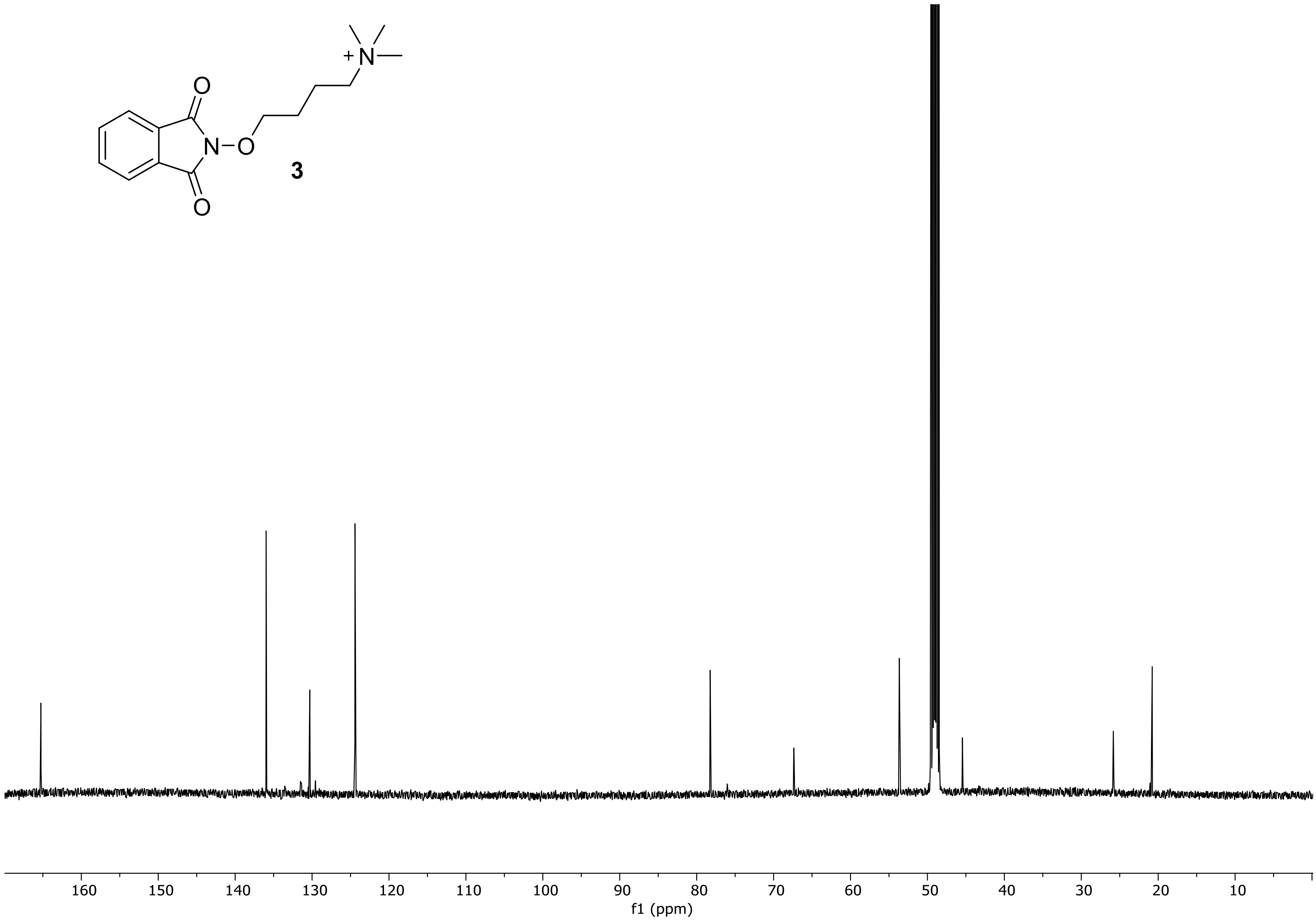


[13]C NMR spectrum (126 MHz) of 4-((1,3-dioxoisoindolin-2-yl)oxy)-*N,N,N*-trimethylbutan-1-aminium bromide (**3**) in methanol-$d_4$.

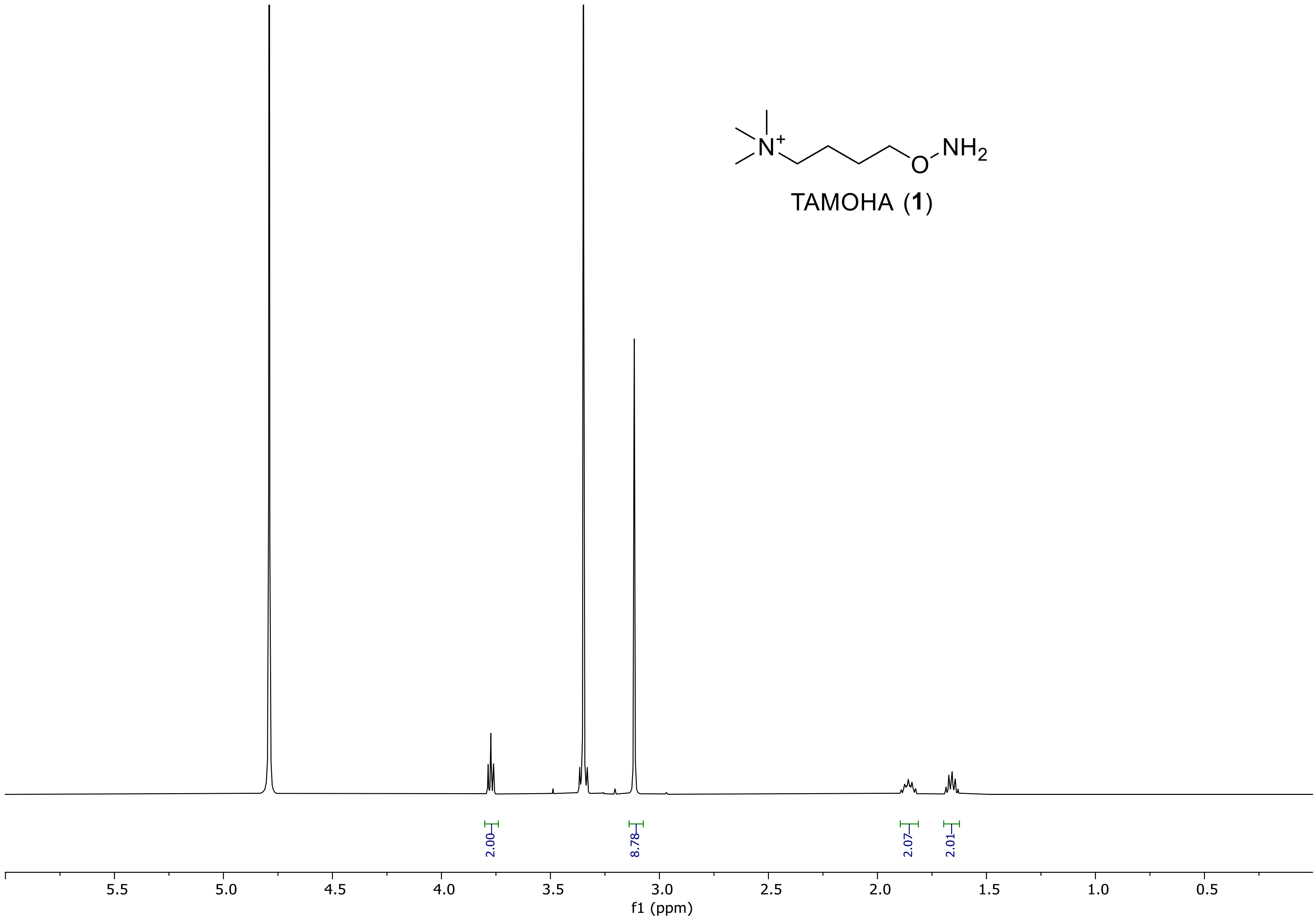


[1]H NMR spectrum (500 MHz) of 4-(aminooxy)-*N,N,N*-trimethylbutan-1-aminium bromide (TAMOHA) (**1**) in $D_2O$.

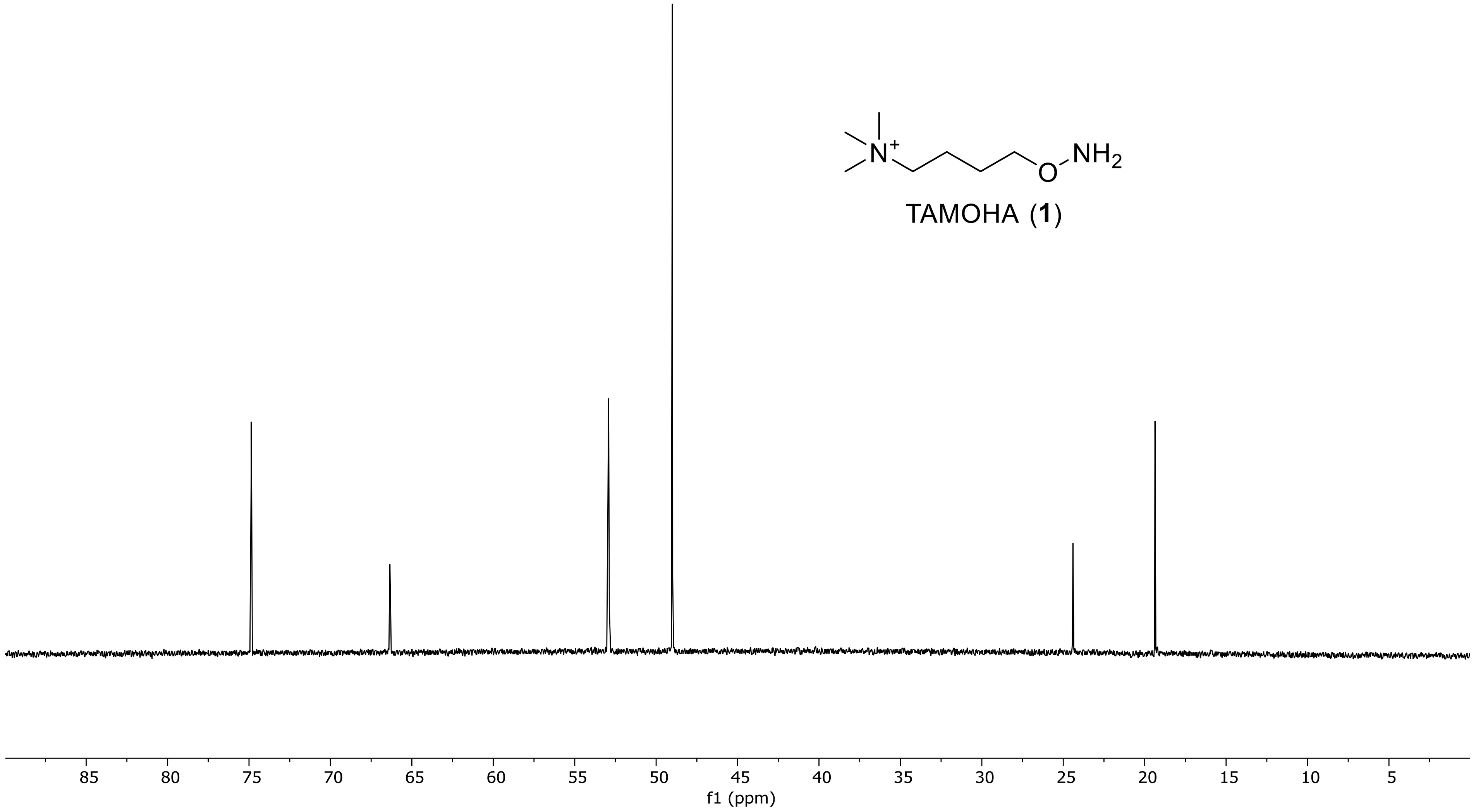


$^{13}$C NMR spectrum (126 MHz) of 4-(aminooxy)-*N,N,N*-trimethylbutan-1-aminium bromide (TAMOHA) (**1**) in $D_2O$.

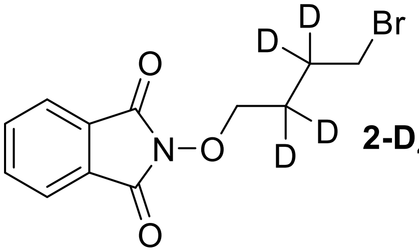

**2-D$_4$**

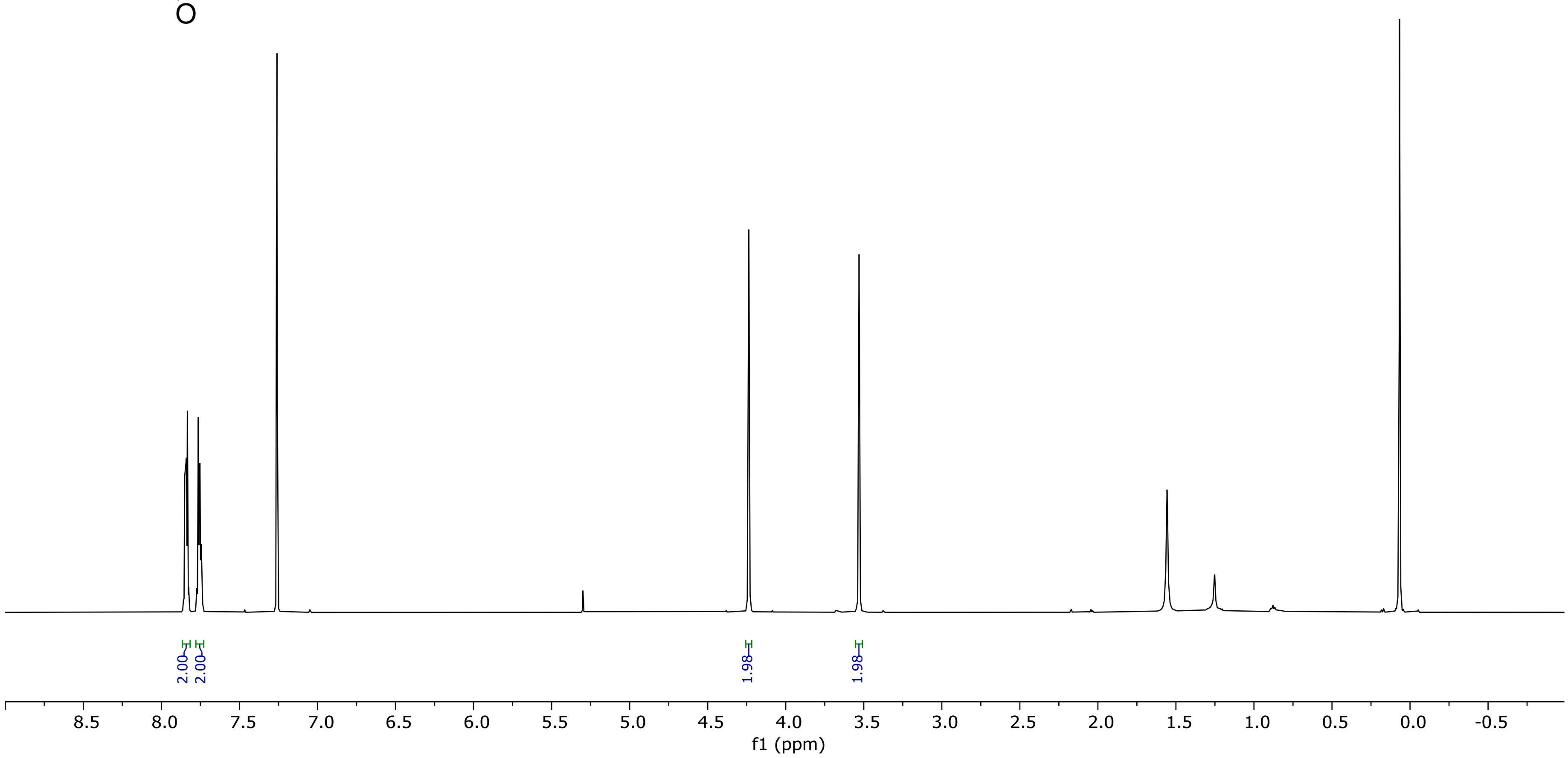


$^{1}$H NMR spectrum (500 MHz) of 2-(4-bromobutoxy-2,2,3,3-D4)isoindoline-1,3-dione (**2-D$_4$**). in chloroform-*d.*

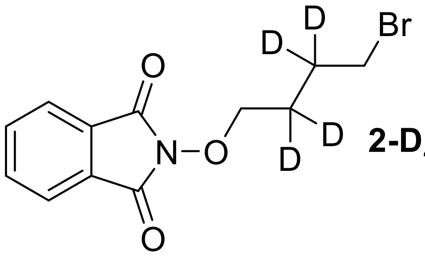


**2-D$_4$**

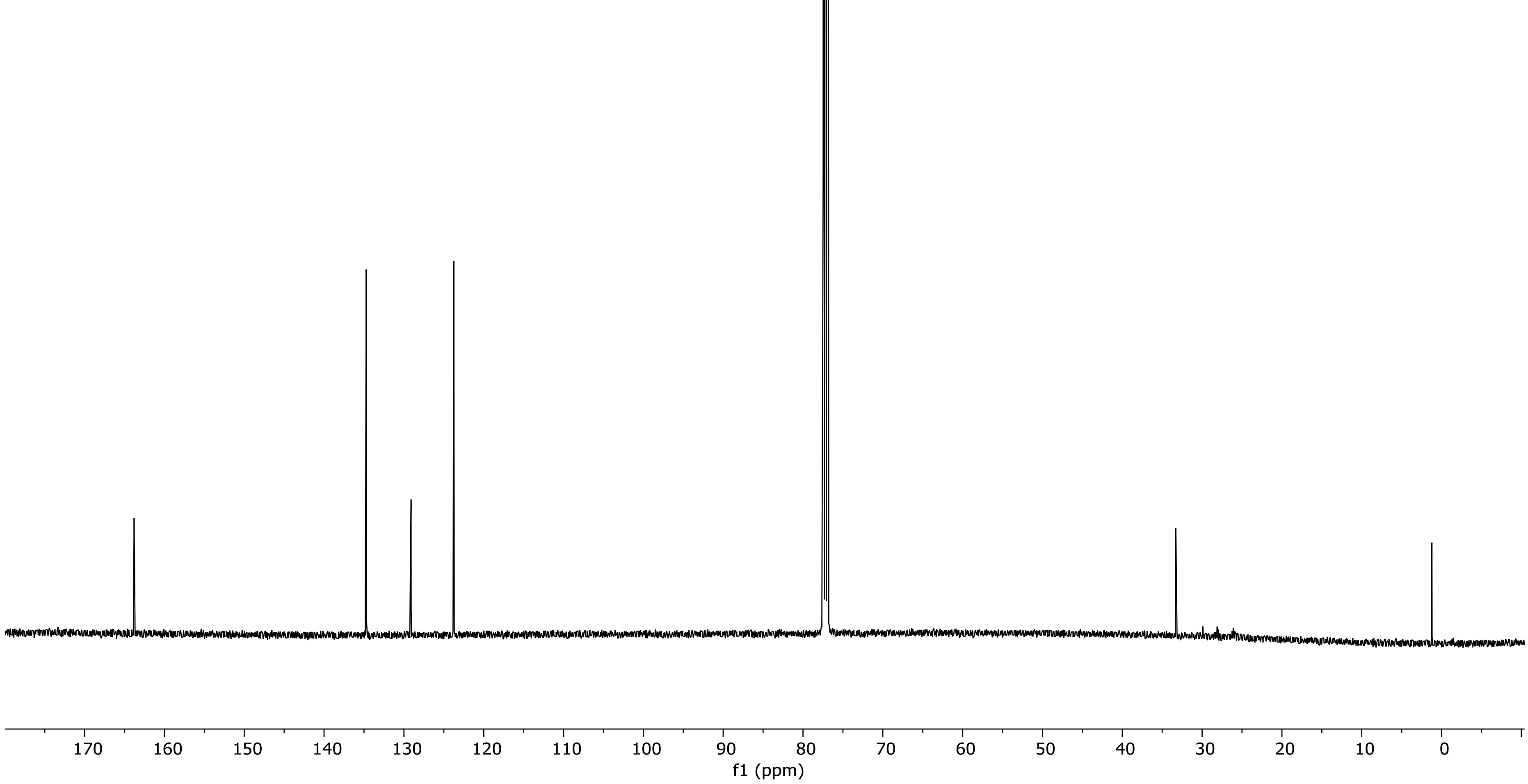


$^{13}$C NMR spectrum (126 MHz) of 2-(4-bromobutoxy-2,2,3,3-D4)isoindoline-1,3-dione (**2-D$_4$**). in chloroform-*d.*

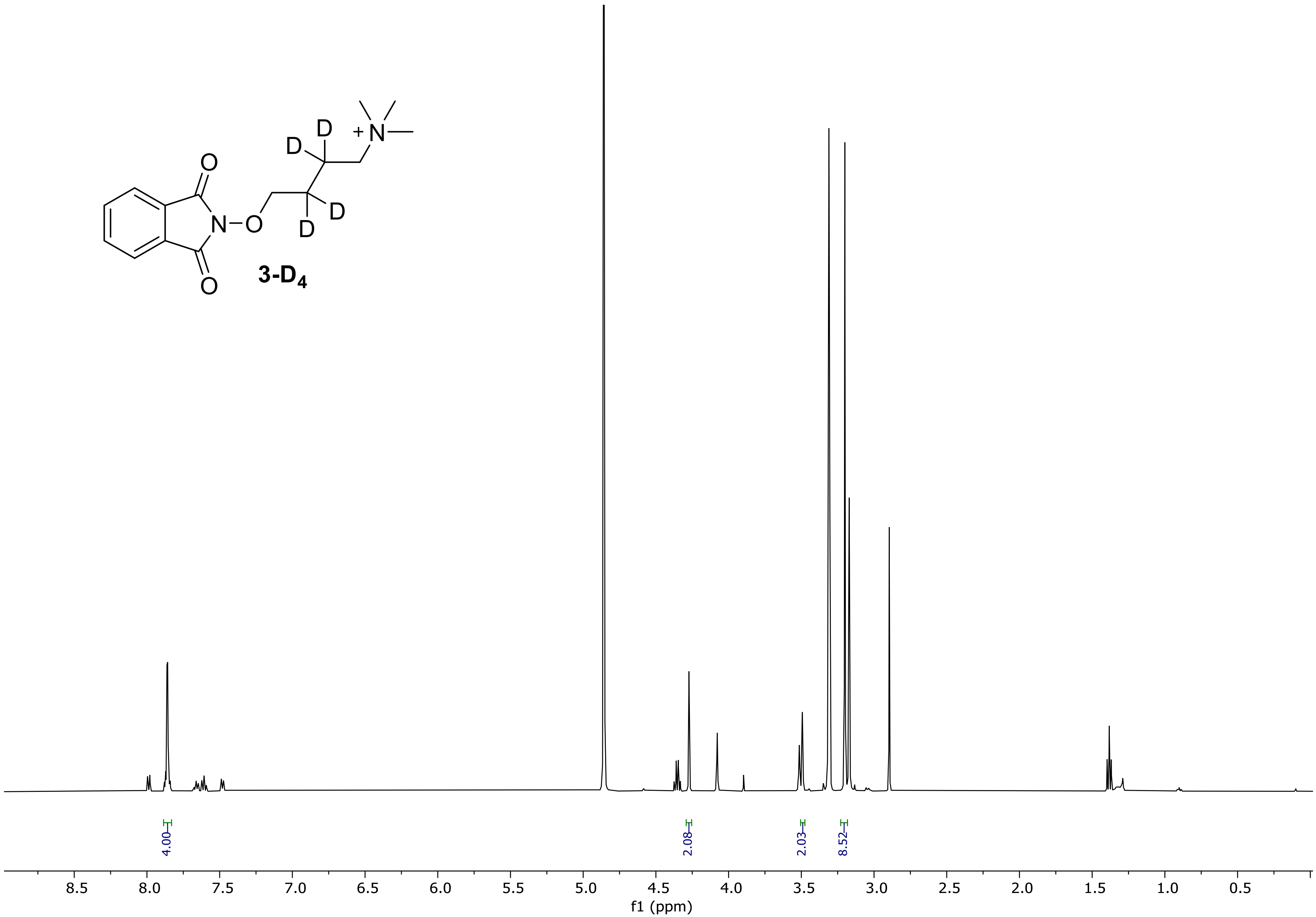


**3-D$_4$**



$^1$H NMR spectrum (500 MHz) of 4-((1,3-dioxoisoindolin-2-yl)oxy)-*N*,*N*,*N*-trimethylbutan-1-aminium-2,2,3,3-D4 (**3 D$_4$**) in methanol-*d$_4$*.

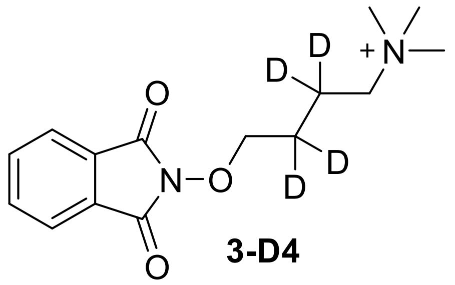


**3-D4**

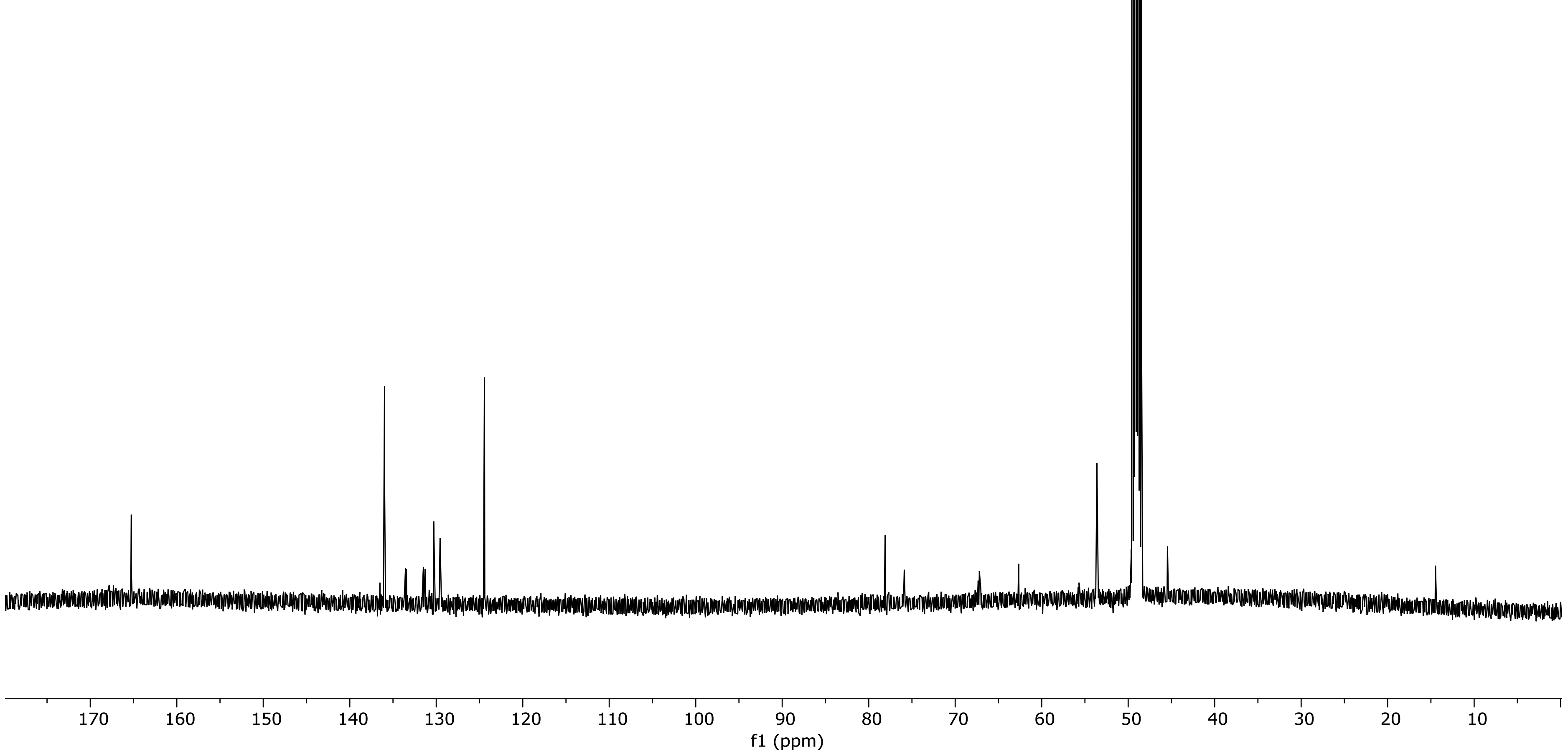


[13]C NMR spectrum (126 MHz) of 4-((1,3-dioxoisoindolin-2-yl)oxy)-*N*,*N*,*N*-trimethylbutan-1-aminium-2,2,3,3-D4 (**3-D$_4$**) in methanol-$d_4$.

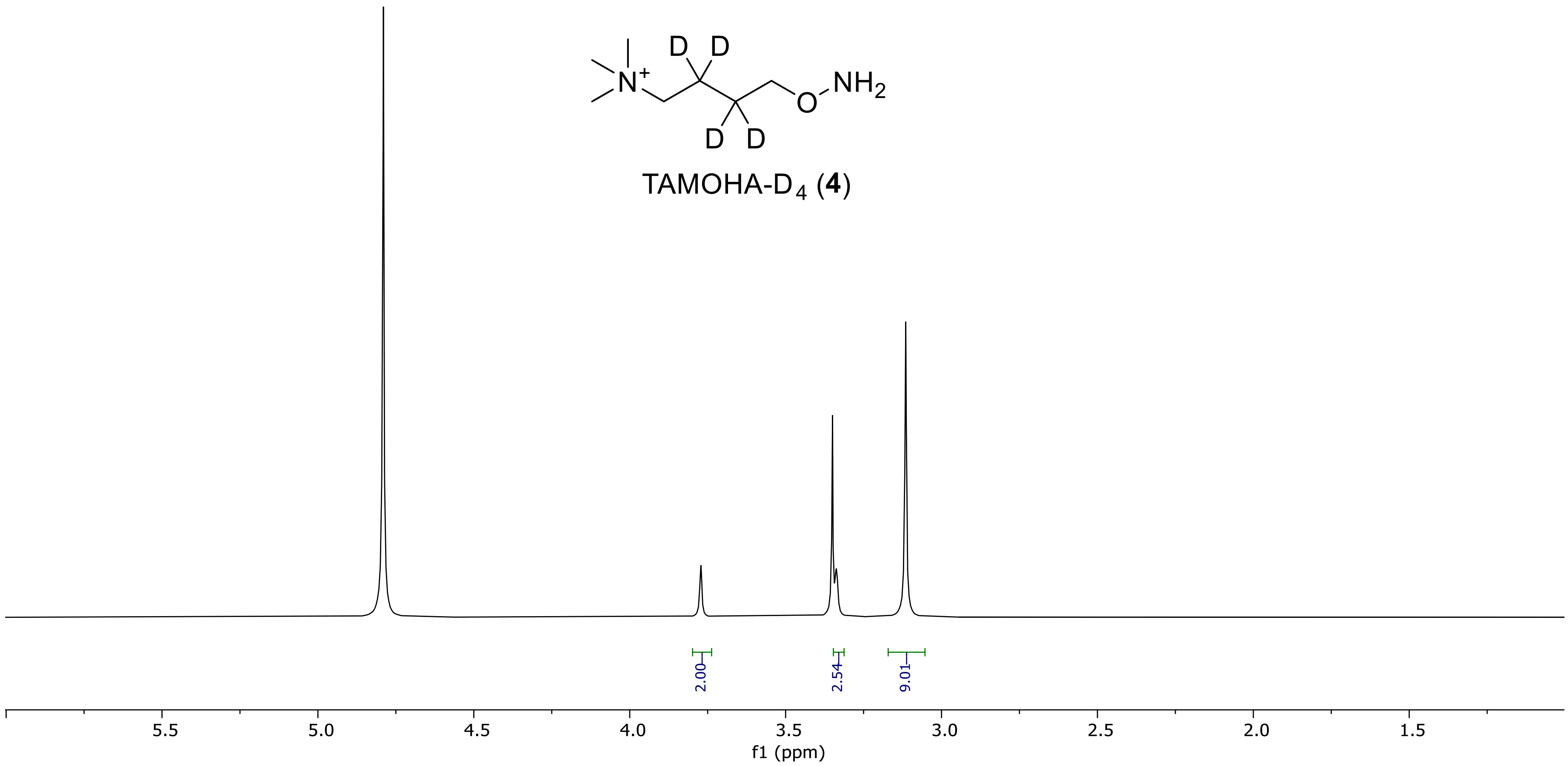


[1]H NMR spectrum (500 MHz) of 4-(aminooxy)-*N,N,N*-trimethylbutan-1-aminium-2,3,3,3-D4 bromide (TAMOHA-$D_4$, 4). in $D_2O$.

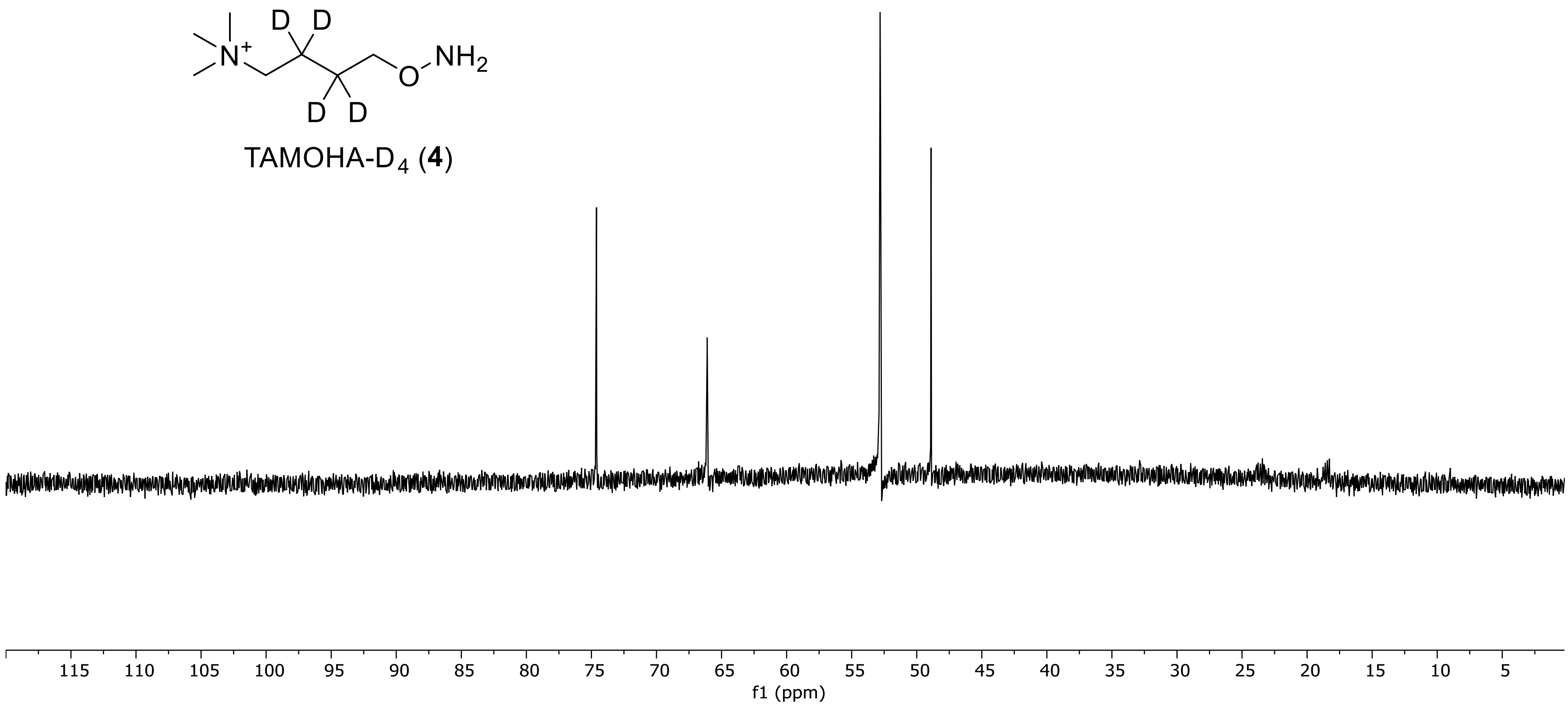


$^{13}$C NMR spectrum (126 MHz) of 4-(aminooxy)-*N,N,N*-trimethylbutan-1-aminium-2,3,3,3-D4 bromide (TAMOHA-$D_4$, **4**). in $D_2O$.

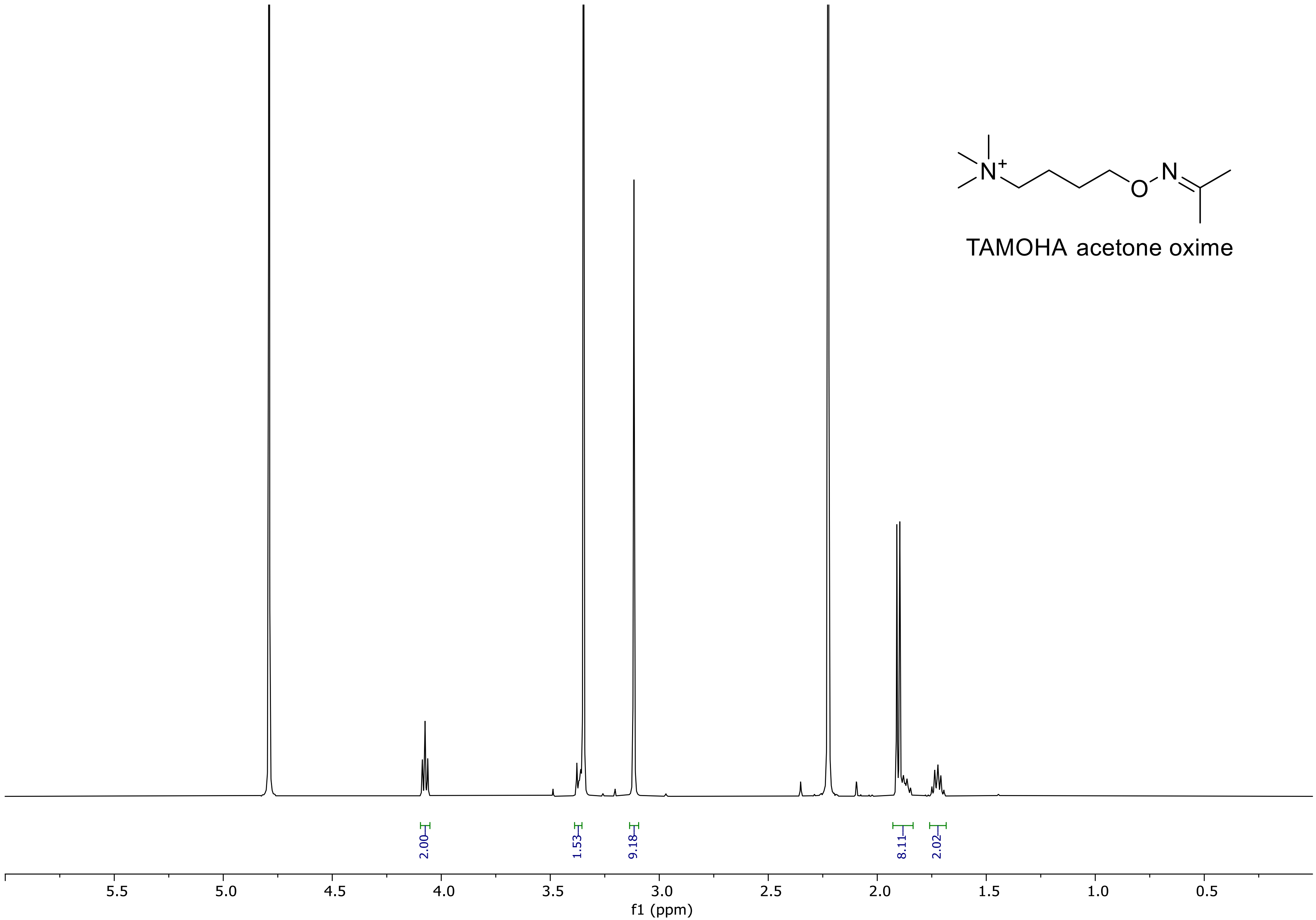


[1]H NMR spectrum (500 MHZ) of *N*,*N*,*N*-trimethyl-4-((propan-2-ylideneamino)oxy)butan-1-aminium (TAMOHA acetone oxime). in $D_2O$.

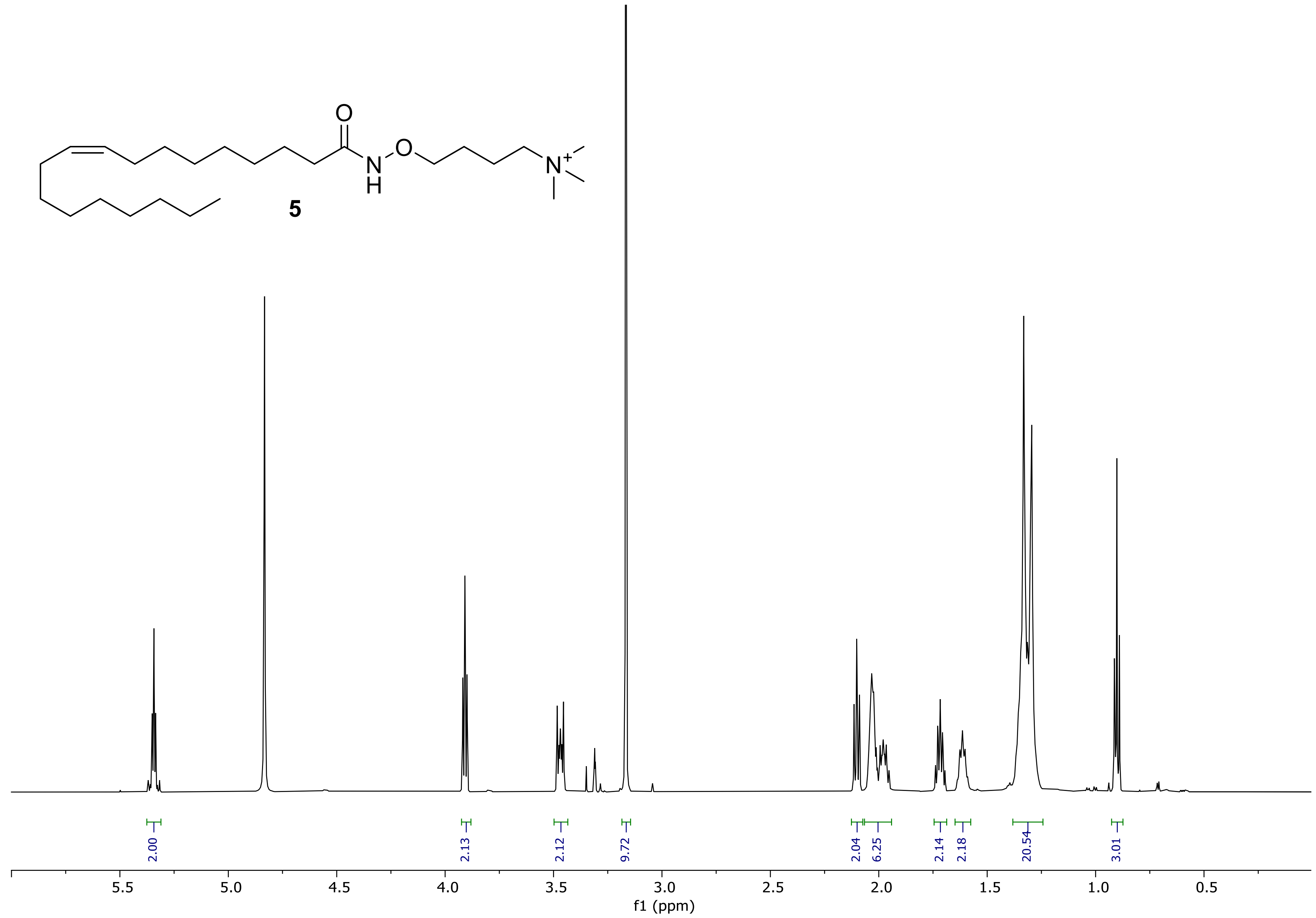


[1]H NMR spectrum (500 MHz) of *N*,*N*,*N*-trimethyl-4-(oleamidooxy)butan-1-aminium (**5**) in methanol-$d_4$.

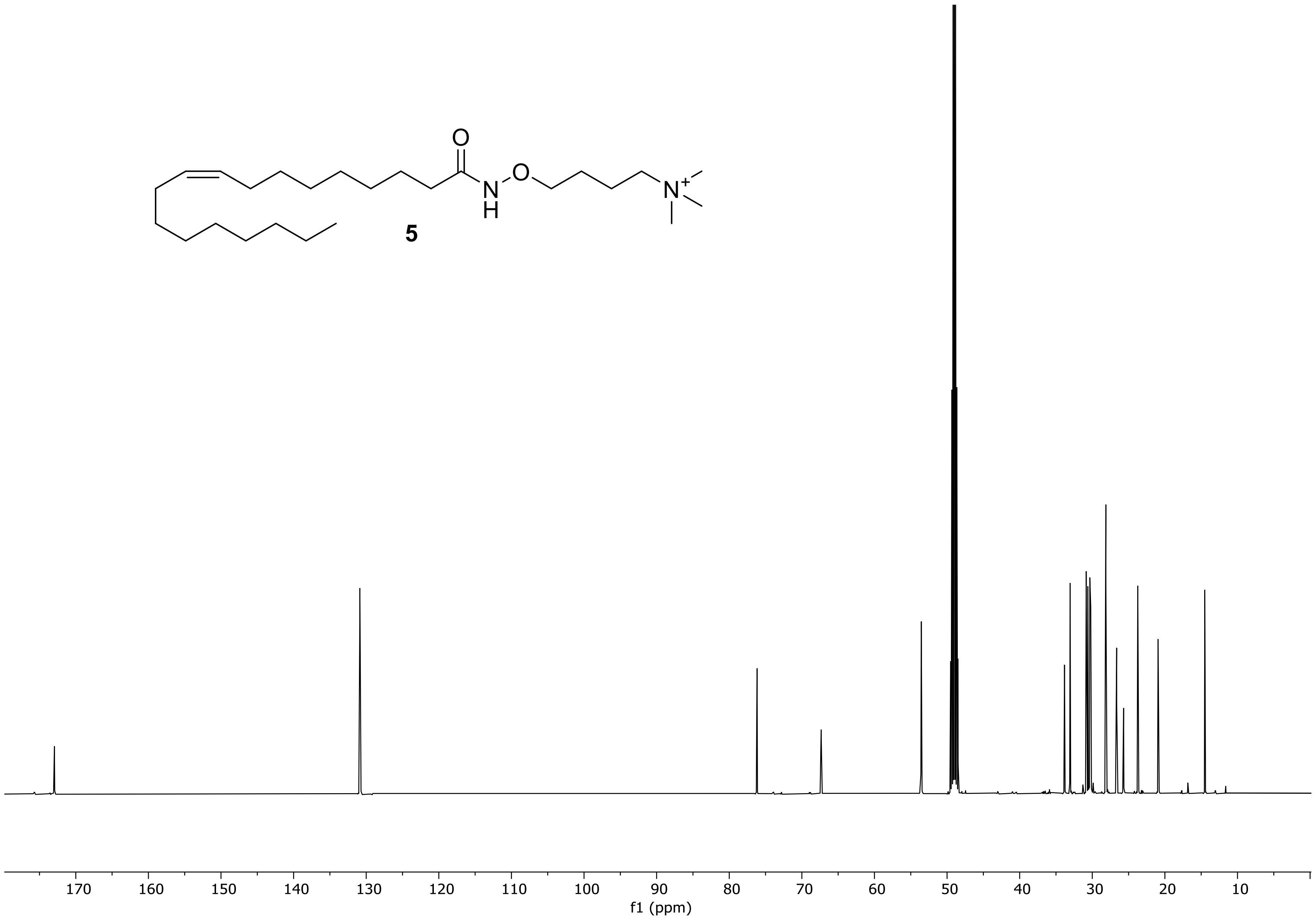


$^{13}$C NMR spectrum (126 MHz) of *N*,*N*,*N*-trimethyl-4-(oleamidooxy)butan-1-aminium (**5**) in methanol-$d_4$.

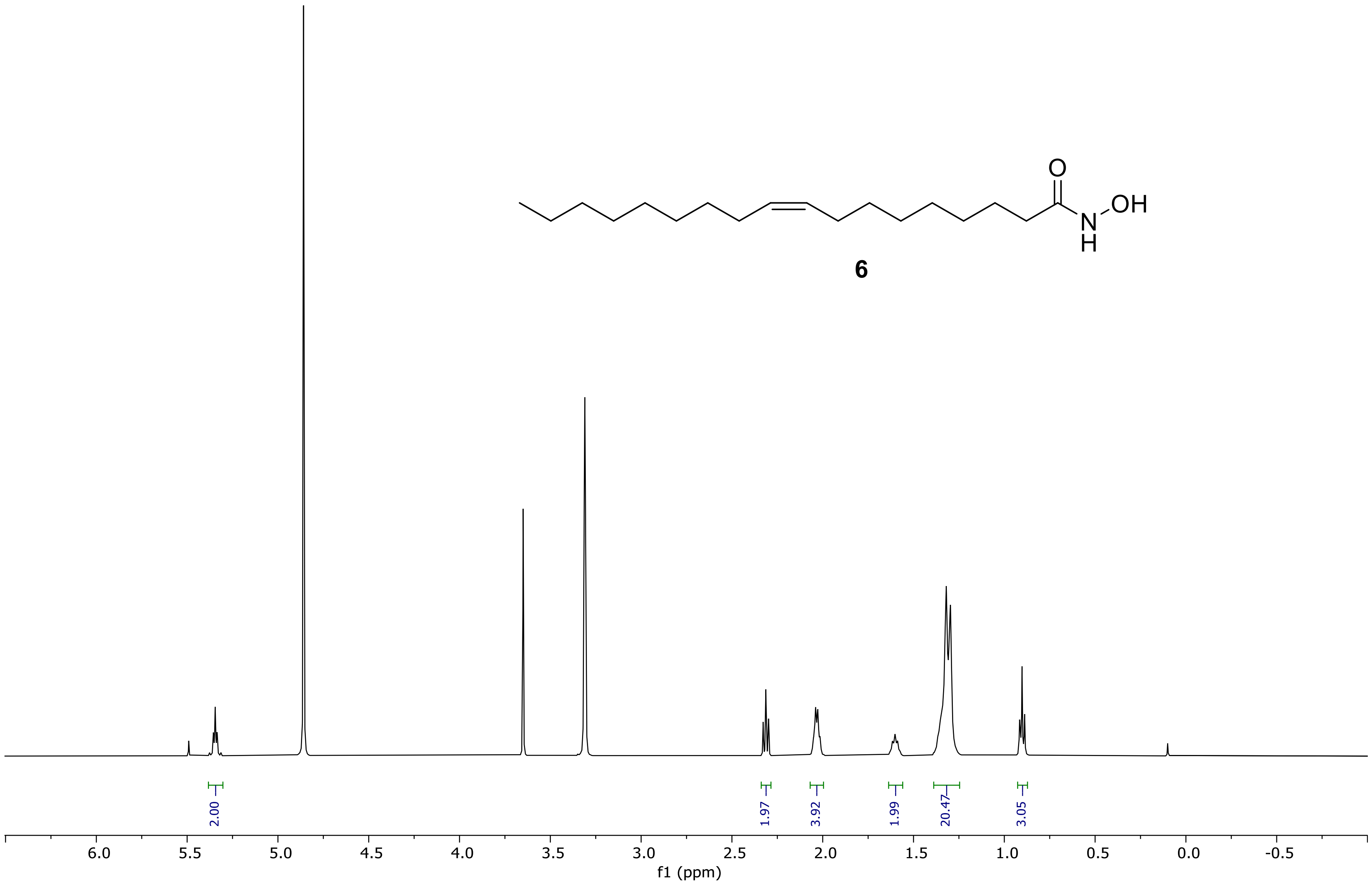


[1]H NMR spectrum (500 MHz) of *N*-hydroxyoleamide (**6**) in methanol-$d_4$.

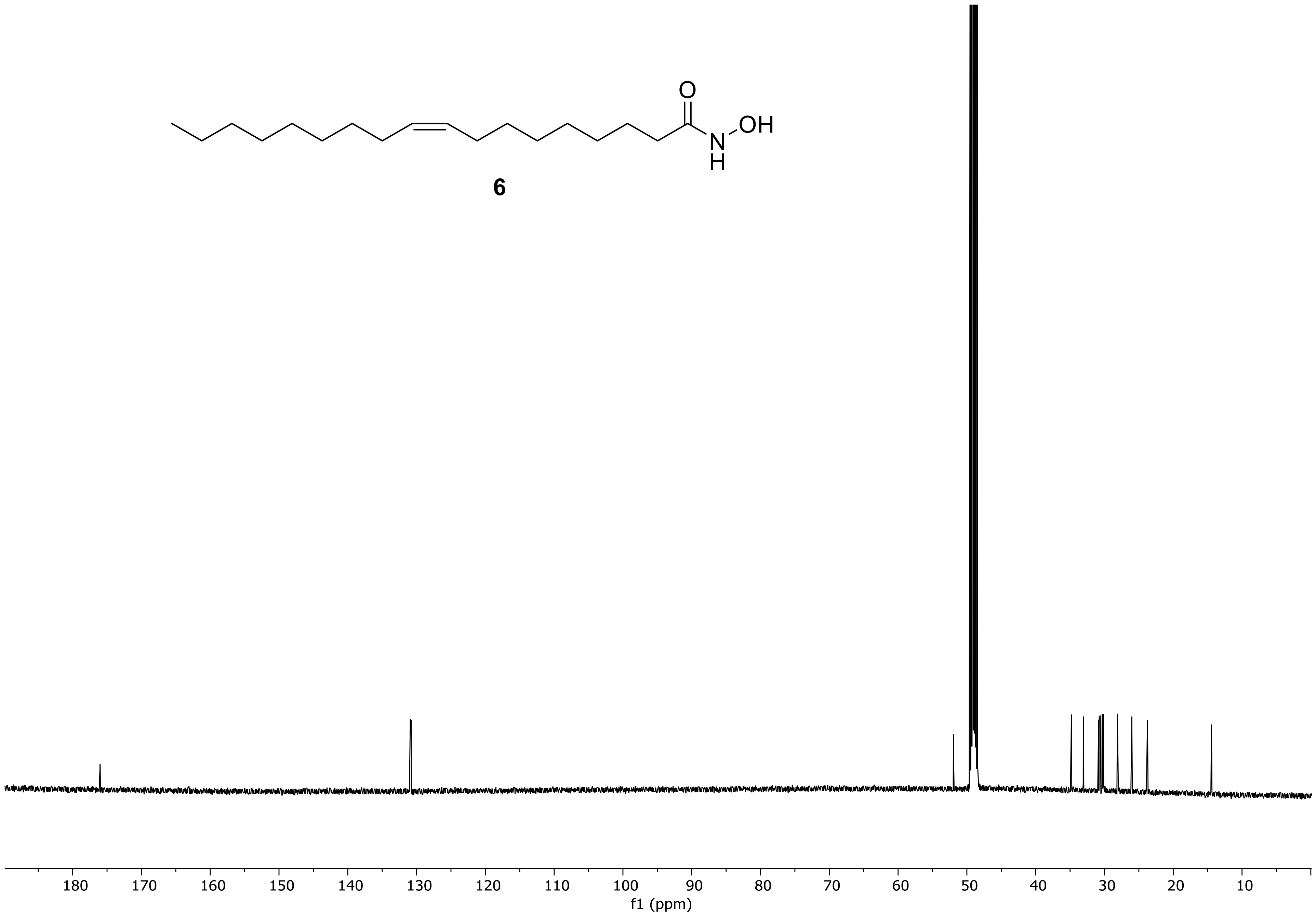


$^{13}$C NMR spectrum (126 MHz) of *N*-hydroxyoleamide (6) in methanol-$d_4$.

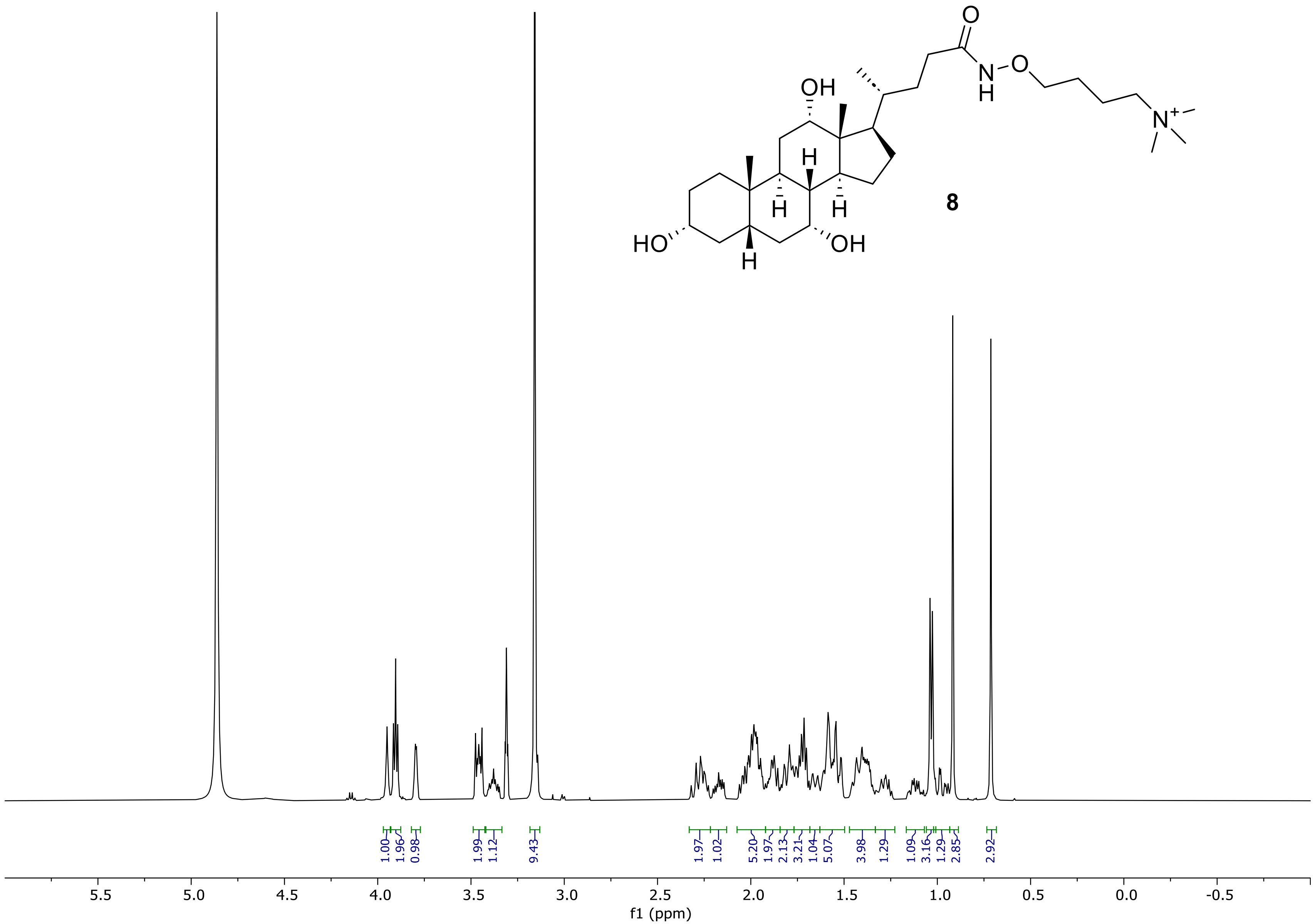


[1]H NMR spectrum (500 MHz) of *N*,*N*,*N*-trimethyl-4-(((*R*)-4-((3*R*,5*S*,7*R*,8*R*,9*S*,10*S*,12*S*,13*R*,14*S*,17*R*)-3,7,12-trihydroxy-10,13-dimethylhexadecahydro-1H-cyclopenta[a]phenanthren-17-yl)pentanamido)oxy)butan-1-aminium (**8**) in methanol-$d_4$.

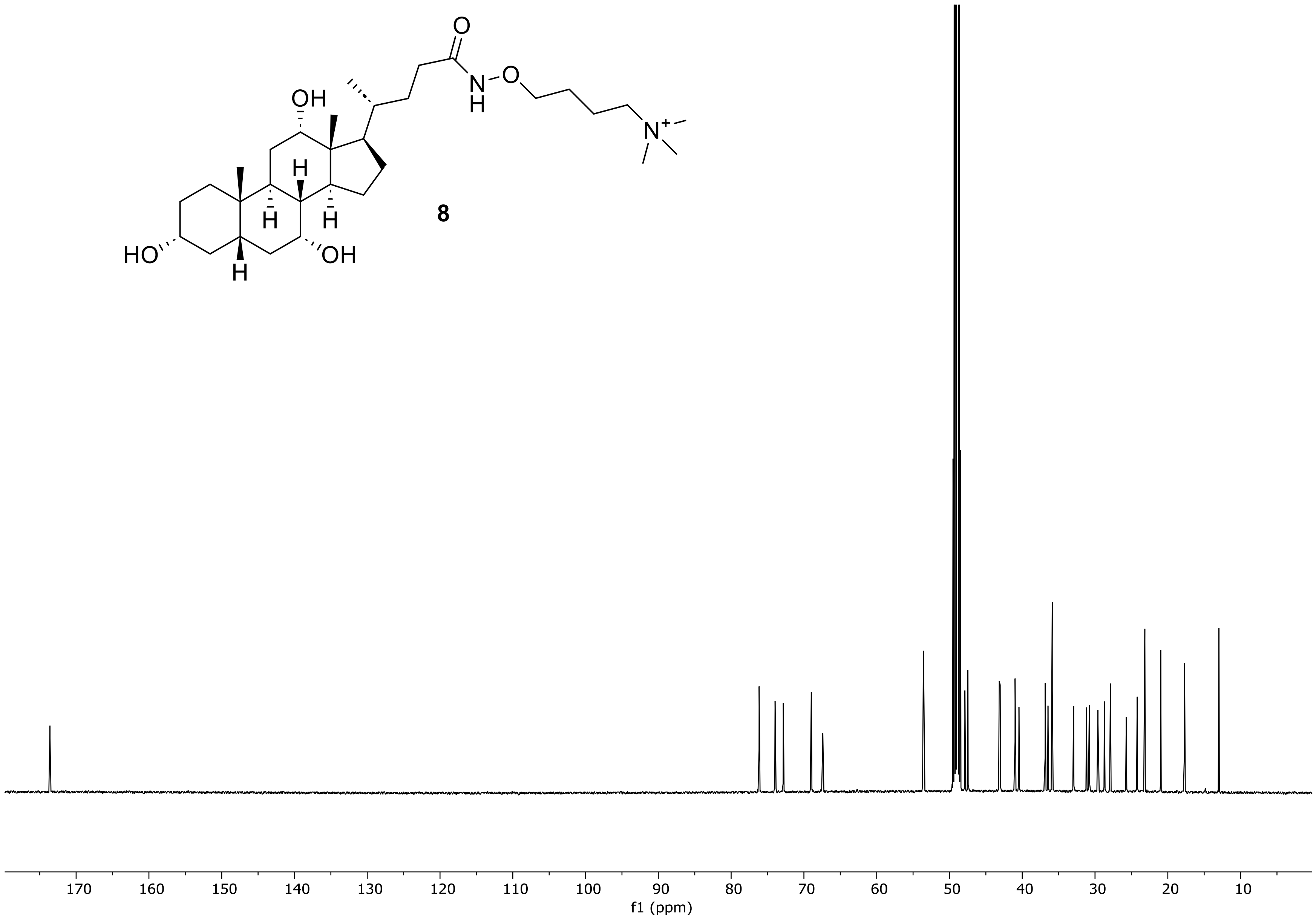


$^{13}$C NMR spectrum (126 MHz) of *N*,*N*,*N*-trimethyl-4-(((*R*)-4-((3*R*,5*S*,7*R*,8*R*,9*S*,10*S*,12*S*,13*R*,14*S*,17*R*)-3,7,12-trihydroxy-10,13-dimethylhexadecahydro-1H-cyclopenta[a]phenanthren-17-yl)pentanamido)oxy)butan-1-aminium (**8**) in methanol-$d_4$.
